# Model estimates of metazoans’ contributions to the biological carbon pump

**DOI:** 10.1101/2021.03.22.436489

**Authors:** Jérôme Pinti, Tim DeVries, Tommy Norin, Camila Serra-Pompei, Roland Proud, David A. Siegel, Thomas Kiørboe, Colleen M. Petrik, Ken H. Andersen, Andrew S. Brierley, André W. Visser

## Abstract

The daily vertical migrations of fish and other metazoans actively transport organic carbon from the ocean surface to depth, contributing to the biological carbon pump. We use an oxygen-constrained, game-theoretic food-web model to simulate diel vertical migrations and estimate global carbon fluxes and sequestration by fish and zooplankton due to respiration, fecal pellets, and deadfalls. Our model provides estimates of the carbon export and sequestration potential for a range of pelagic functional groups, despite uncertain biomass estimates of some functional groups. While the export production of metazoans and fish is modest (~20% of global total), we estimate that their contribution to carbon sequestered by the biological pump (~ 800 PgC) is conservatively more than 50% of the estimated global total (~1300 PgC) and have a significantly longer sequestration time scale (~250 years) than previously reported for other components of the biological pump. Fish and multicellular zooplankton contribute about equally to this sequestered carbon pool. This essential ecosystem service could be at risk from both unregulated fishing on the high seas and ocean deoxygenation due to climate change.

## 1 Introduction

Many marine organisms – from zooplankton to fish – perform diel vertical migrations (DVM) (McLaren, 1963; Klevjer et al., 2016), as they seek both to access food and to avoid predators. Small planktivorous zooplankton, for instance, feed close to the surface at night, and migrate to depth during daytime to reduce their predation risk from visual predators (Zaret and Suffern, 1976; Lampert, 1993). In turn, higher trophic levels organise their vertical migrations to take advantage of their migrating prey while themselves avoiding predators, a process that can penetrate deep into the oceans interior as a staggered trophic relay known as Vinogradov’s ladder (Vinogradov, 1962; Hernández-León et al., 2019). DVM within a marine pelagic community is therefore the product of a co-adaptive “game” where many animals seek to optimize their migration patterns relative to the migration patterns of their respective prey, predators, and conspecifics (Hugie and Dill, 1994; Pinti and Visser, 2019; Pinti et al., 2019). These interacting DVM patterns govern trophic interactions (Bandara et al., 2021) and affect global biogeochemical cycles (Buesseler and Boyd, 2009). In particular, migrating organisms consume organic carbon on average at shallower depths and transport it to deeper depths where it is released through respiration or excretion. This process, termed the active biological pump (or migrant pump), is highly efficient at sequestering carbon, as it injects carbon directly at depth and bypasses the remineralization experienced by passively sinking particles in the upper ocean (Boyd et al., 2019). We call carbon injection the depth-dependent biologically-mediated source of DIC, in contrast to carbon export that refers to organic carbon transported below a reference depth. Even though these two notions are closely related, carbon export and injection can differ as, for example, vertical migrants can consume detritus at depths and bring it back to surface waters.

The effects of DVM on carbon export, particularly that mediated by zooplankton, has garnered considerable attention (Longhurst et al., 1990; Steinberg et al., 2000; Hansen and Visser, 2016; Archibald et al., 2019; Gorgues et al., 2019), and recent biogeochemical models indicate that migrating organisms transport between 1 and 30 mgCm^−2^day^−1^ through the base of the euphotic zone (Archibald et al., 2019; Aumont et al., 2018), typically constituting around 15 - 20% of local export flux. These studies, however, do not address carbon sequestration, an arguably more relevant metric in assessing the biological carbon pump and its various components. In particular, a recent biogeochemical analysis suggests that 1300 PgC is stored in the ocean as a consequence of organic matter remineralization (Carter et al., 2021) and can be directly compared to other important carbon reservoirs in the earth system (e.g. atmosphere 870 PgC, soils 1700 PgC, marine sediments 1700 PgC, dissolved organic carbon in the oceans 700 PgC, and marine biota 3 PgC; IPCC AR6 2021 (Canadell et al., 2021)). More recently, in a bio-geochemically constrained model, Nowicki et al. (2022) were able to decompose the relative pathways of ocean carbon sequestration as (approximately) 100 PgC by subducting water masses, 200 PgC by sinking phytodetritus aggregates, 850 PgC by sinking fecal pellets and 150 PgC by the metabolism of vertical migrants.

Here we seek to refine these estimates, specifically disentangling the contribution of the various taxa of migrants (mesozooplanton, macrozooplankton, mesopelagic fish, forage fish, large pelagic fish and jellyfish), their interlinked migrations and trophic interactions (Vinogradov’s ladder), and the different transport pathways involved (respiration, fecal pellets, and dead-falls). The role of fish, for instance, is currently poorly resolved (Davison et al., 2013; Saba et al., 2021). Mesopelagic fish are potentially of particular importance, in part because of their high biomass (Irigoien et al., 2014; Proud et al., 2019), their conspicuous DVMs, and their excretion of fast-sinking fecal pellets (Klevjer et al., 2016; Davison et al., 2013). Our motivation goes beyond simply a better understanding of the biological carbon pump. Notably, the role of marine biota in sequestering carbon in the ocean is well in excess of their biomass (1300 PgC vrs 3 PgC). Perhaps even more revealing, 1000 PgC (more than 75% total sequestered carbon in the ocean) seems to have passed through zooplankton (Nowicki et al., 2022). Therefore, human disruptions to metazoan communities, for example due to fishing or pollution, could potentially have a large impact on ocean carbon sequestration well beyond the loss of biomass alone. For instance, it has been recently noted that global fisheries have had a comparable impact on ocean biogeochemistry as the anthropogenic release of greenhouse gasses (Bianchi et al., 2021). Our interest is in identifying those components of the marine ecosystem, taxa and pathways, that are particularly important in ocean carbon storage.

Here, we use a pelagic food-web model to investigate the potential impact of different metazoan functional groups and pathways on global ocean carbon budgets. We specifically consider how groups and pathways directly inject respired and egested carbon at depth, and their contribution to ocean carbon sequestration. We use a game theoretic approach to determine optimal DVM patterns for all migrant taxa, and compare these to acoustic observations. Throughout we are mindful of uncertainties, particularly in model estimates of fish biomasses, and provide confidence intervals for all estimates. The global biomass of mesopelagic fish for instance remains poorly quantified and we impose a 10 fold range in its uncertainty in accordance with best estimates from observations. As is done in previous studies (Carter et al., 2021; Nowicki et al., 2022) we use oxygen observations as a constraint on carbon sequestration.

## 2 Methods

### 2.1 Vertical migration model

The behavioural part of our model represents a 1D pelagic community, from surface waters to mesopelagic depths (figure 1). The model resolves migrating functional groups: meso-zooplankton, macro-zooplankton, forage fish, large pelagic fish, tactile predators (i.e. jellyfish), and mesopelagic fish, as well as non-migrating resources of phytoplankton and micro-zooplankton. We consider behavioural responses to be rapid compared to population dynamics (Pinti et al., 2021) so that the biomass of all groups can be considered fixed. The vertical distribution of phytoplankton depends on the mixed layer depth. Large pelagic fish are assumed to be uniformly distributed vertically as they are proficient swimmers that are able to move up and down the water column several times a day (Holland et al., 1992; Thygesen et al., 2016). This also implies that the predation risk they impose on other groups depends only on their total biomass and on physical depth effects (e.g. light attenuation). All other functional groups can move in the water column and our model computes the optimal day and night distribution of all organisms in the water column simultaneously. Detritus is created by organisms (through fecal pellet production or by natural mortality), sinks, and is degraded by bacteria or ingested by macro zooplankton along the way.

**Figure 1.**
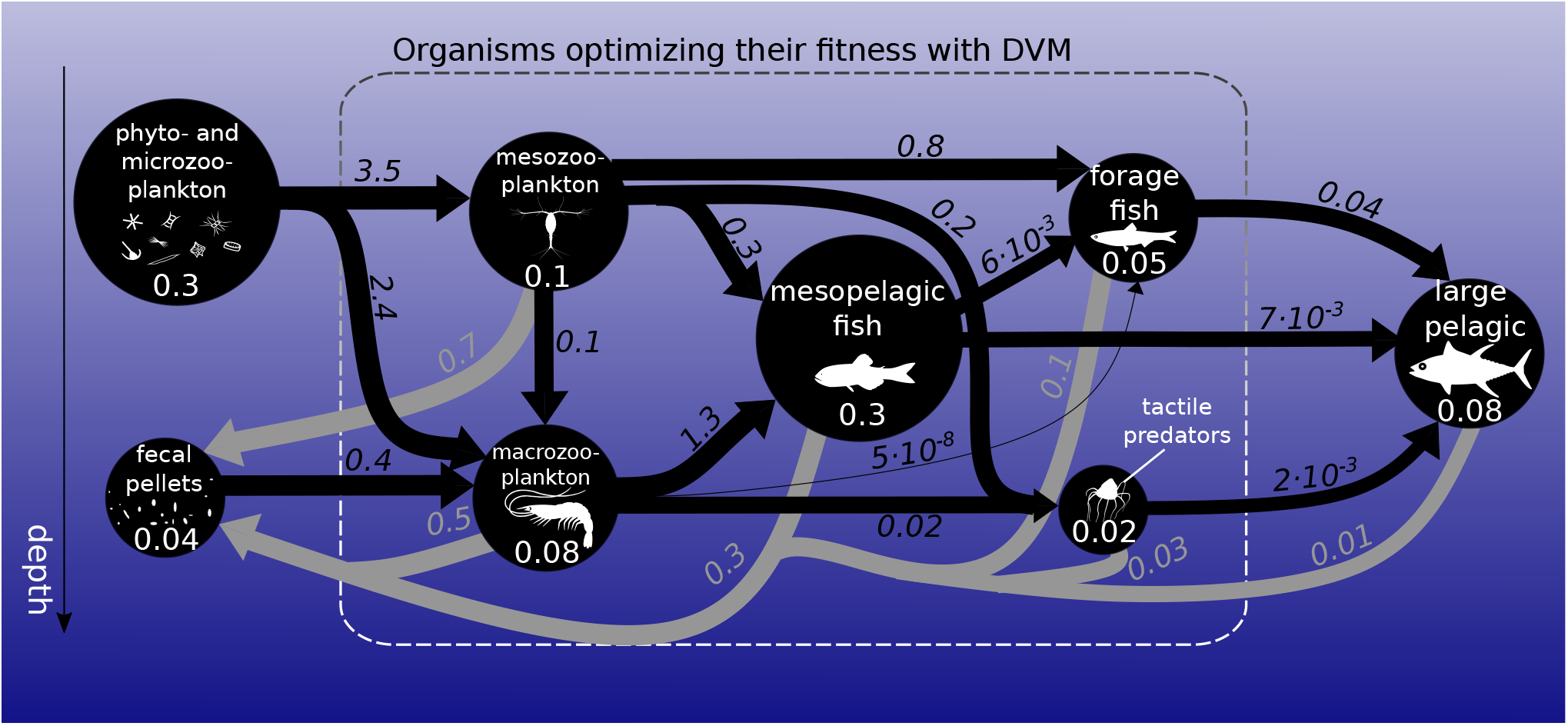
Biomass (circles) and fluxes (arrows) in the food-web integrated over the global ocean. Biomasses are in PgC (white numbers). Black arrows represent ingestion while grey arrows represent fecal pellet excretion in PgC yr^−1^. Arrow widths and circle diameters are proportional to the logarithm of the fluxes and biomasses they represent. Respiration losses are not represented here. The dashed box surrounds the functional groups that optimize their day and night vertical distribution with DVM.

An organism’s optimal strategy (i.e. day and night positions) maximises its fitness given the position of all other organisms in the water column. As an individual selects a strategy, the fitness of its prey, predators and conspecifics also varies. Hence, the optimal strategy of all individuals is intrinsically linked to the optimal strategy of all other players. The optimal strategies for all individuals is attained at the Nash equilibrium (Nash, 1951), where no individual can increase its fitness by changing its strategy. The Nash equilibrium is found using the replicator equation (Hofbauer and Sigmund, 2003; Pinti and Visser, 2019; Pinti et al., 2019). In short, the fraction of the population following a particular strategy grows proportionally to the fitness related to that strategy, and the algorithm is run until steady state is reached. Importantly, this computational scheme allows for mixed strategies to emerge (Sainmont et al., 2013; Pinti and Visser, 2019) so that varying proportions of each population can follow different strategies. Thus, the percentage of each population that performs vertical migrations is an emergent property of the model.

We use Gilliam’s rule (Houston et al., 1993) as a fitness measure, i.e. maximizing the ratio of growth over mortality. In a steady environment, this approximates the optimal behaviour that will ensure maximum life-time reproductive success assuming uniform future conditions. While incomplete in a full accounting of life history strategies, this simple heuristic provides a reasonably accurate estimate for fitness maximizing behaviour (Sainmont et al., 2015). Fitness is affected by the availability of and competition for food, by the presence or absence of predators, and by environmental conditions such as light levels, temperature, and oxygen concentrations. Light levels also vary between day and night, creating the possibility for organisms to perform DVM – if the optimal strategy is to change vertical position during day and night.

The growth rate of organisms is the assimilation rate minus standard metabolic rate and migration cost. The mortality rate is the mortality due to predation plus a small natural mortality from non-modelled sources. Predators and prey swim at a constant speed and encounter each other depending on the clearance rate of the predator (for visual predators, this varies vertically due to light attenuation in the water and between day and night). The probability of capture in each encounter event depends on the escape speed of prey and the attack speed of predators, both varying with the aerobic scope of the corresponding organism (which depends on the local oxygen and temperature conditions). The ingestion rate of each organism is modelled by a type II functional response, except for jellyfish that follow a type I functional response with no saturation at high prey concentrations (Holling, 1959; Titelman and Hansson, 2006). An ingested prey is then assimilated with a certain efficiency. The fraction not assimilated is egested as fecal pellets. Moreover, organisms dying of natural mortality sink as carcasses with a fast sinking velocity, bringing carbon to depths which released as CO_2_ as carcasses get degraded by bacteria. All details, equations and parameters for fitness calculations are given in the supplementary material.

### 2.2 Global simulations and carbon sequestration estimates

This 1D behavioural model is run at the scale of an individual 2-degrees by 2-degrees ocean water column, with vertical limits at the surface and the seafloor and a vertical resolution of 20 m. The model is run repeatedly by varying the spatial location of the water column within a 2-degrees global grid. Each grid box is assumed to be laterally homogeneous and to not interact with neighbouring grid cells. Inputs for the model at each location come from global gridded datasets of global biomass, temperature, oxygen levels and mixed layer depths. Global estimates of biomass distributions come from previously published model results (COBALT (Stock et al., 2014, 2017) and FEISTY (Petrik et al., 2019)). We use the results from COBALT to ascribe the biomass distribution of unicellular plankton (phytoplankton and microzooplankton), mesozooplankton (e.g. copepods) and macrozooplankton (e.g. krill). We use FEISTY model results to provide the biomass distributions of forage fish and large pelagic fish. While the biomass of mesopelagic fish is currently poorly constrained (within an order of magnitude), its distribution can be estimated from acoustic backscatter (Proud et al., 2017, 2019). We use a range of mesopelagic fish biomass which underlies much of the uncertainty in our results. Finally, we use a conservative constant estimate of jellyfish biomass. Environmental conditions (temperature, oxygen, light attenuation coefficient, and mixed layer depth) are taken from the World Ocean Atlas 2018 (Locarnini et al., 2019; Garcia et al., 2019). Additional information on biomass estimates and environmental parameters can be found in the supplementary material.

We do not include coastal areas (shallower than 500 m) nor latitudes higher than ±45^*°*^. Coastal areas were not included because our model is unsuited to shallow continental shelf regions, and high latitudes were not included because of their seasonality – although they can have important consequences for carbon export, through, e.g., the seasonal migration of zooplankton (Jónasdóttir et al., 2015). Our modelled domain constitutes about 65% of the global ocean’s areal extent, and nearly 80% of the pelagic ocean with depths > 500 m.

Once the global behaviour of organisms is computed, we compute the amount of carbon respired, egested as fecal pellets, or sinking as carcasses for each functional group. This directly provides us with global carbon export and injection estimates. The animal respiration rates (basal respiration and other losses – an aggregate of all processes not accounted for in the model, such as specific dynamic action and reproduction) and bacterial respiration rates (due to the degradation of fecal pellets and carcasses) are then used to compute the carbon sequestration by each pathway using a data-constrained steady-state ocean circulation inverse model (OCIM, DeVries and Primeau, 2011; DeVries, 2014; Holzer et al., 2021), providing estimates of the amount of carbon sequestered in the oceans via the different pathways, assuming equilibrium conditions. Dividing the amount of carbon sequestered by the corresponding global injection yields the sequestration time of respired carbon, a measure of the time scale on which carbon is sequestered.

## 3 Results

Global estimates of biomass from published sources (COBALT (Stock et al., 2014, 2017) and FEISTY (Petrik et al., 2019)) as well as mesopelagic fish and jellyfish sum to about 1 (0.6-1.3) PgC (figure 1). Here and throughout the manuscript, numbers in parenthesis refer to the most extreme results of the sensitivity analysis. Given a total marine biomass estimate of 3 PgC (IPCC AR6 2021) and that we do not include either shelf seas or high latitudes seas, this seems like a reasonable estimate. The spatial variation of the biomass of each metazoan group can be found in SI (figure S5). The relatively large uncertainty in total biomass (± 40%) is due to the order of magnitude uncertainty in mesopelagic fish biomass. Results from the game-theoretic DVM model provide the day and night vertical distributions of each metazoan group (figure 2). These vertical distributions are largely driven by resource availability (living and dead organic material) and predation risk, i.e. depth dependent clearance rates, and the vertical distribution overlap of predators and prey. As such, both trophic transfer to higher trophic levels, and the egested flux to fecal pellets are emergent estimates from the DVM model (figure 1).

**Figure 2.**
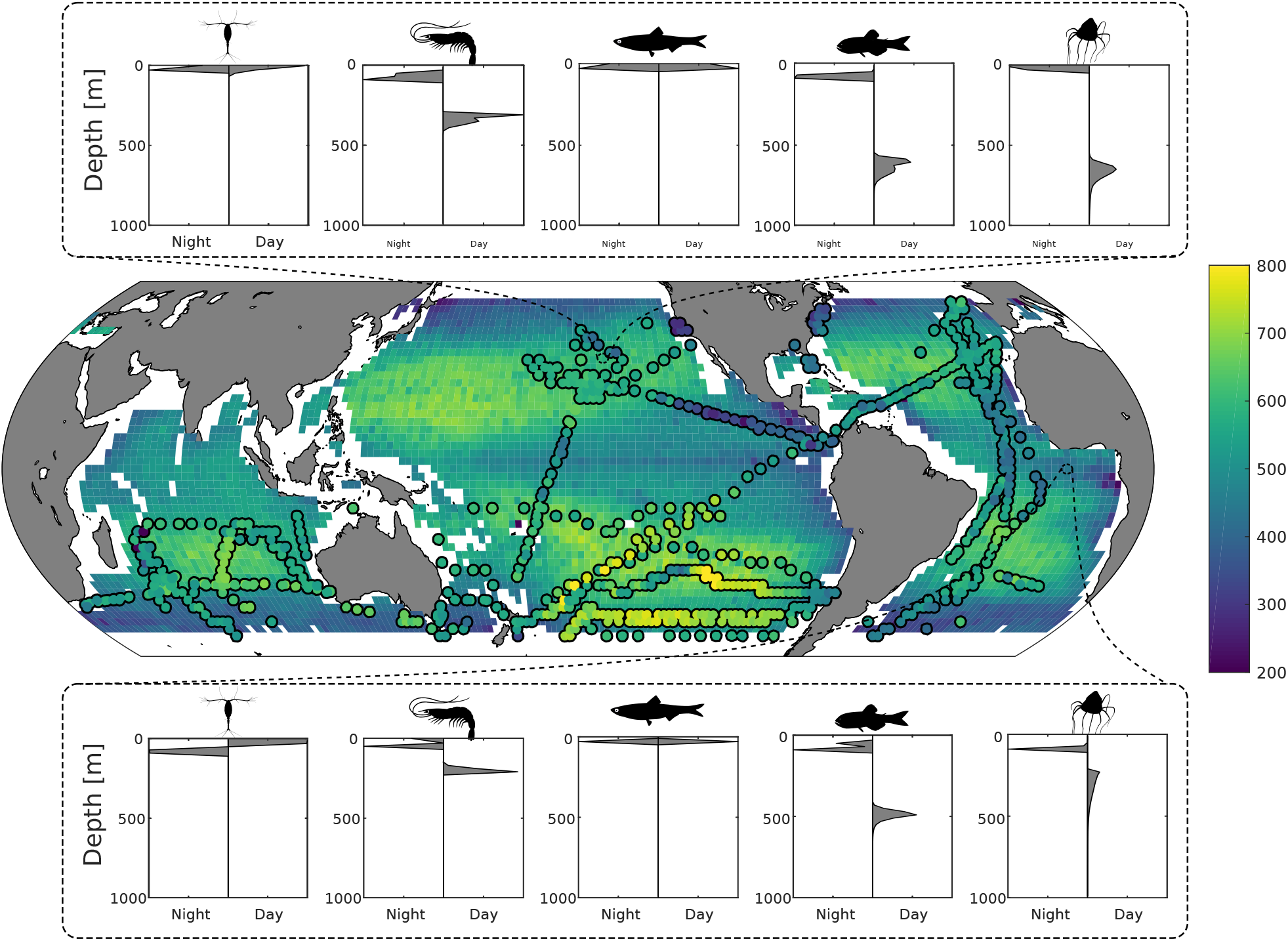
Top & bottom panels: predicted day and night depth distribution of meso-zooplankton, macro-zooplankton, forage fish, mesopelagic fish and jellyfish at 26^*°*^N 152^*°*^W and 7^*°*^S 0^*°*^E respectively. Middle panel: Predicted fish mean depth (in meters) during day-time, weighted by biomass. Circles overlaid are the observed mean depths weighted by echo intensity recorded using 38 kHz echosounders (Klevjer et al., 2016; IMOS, 2021; Polar Data Centre British Antarctic Survey, 2020).

We find the strongest trophic coupling between mesopelagic fish (total biomass of 0.32 (0.06-0.64) PgC) and macro-zooplankton, with mesopelagic fish ingesting 1.3 (0.3-2.4) PgC of macro-zooplankton annually. Deadfalls and fecal pellets produced by metazoans in the euphotic zone contribute to a sinking flux of 1.0 (0.6-1.5) PgC yr^−1^ at the base of the euphotic zone (see figure 4 for local estimates). Additionally, 0.4 (0.2-0.7) PgC yr^−1^ of fecal pellets and 0.1 (0.1-0.3) PgC yr^−1^ of deadfalls are produced below the euphotic zone by metazoans, which also respire 1.1 PgC yr^−1^ (0.6 - 1.7) PgC yr^−1^ through basal respiration and 0.4 (0.1-0.8) PgC yr^−1^ through other losses below the euphotic zone globally (see figures S11 and S12 for local estimates). Table 1 provides a summary of carbon injection rates due to the different pathways – basal respiration, fecal pellets, deadfalls, and other losses – for all functional groups.

**Table 1.**
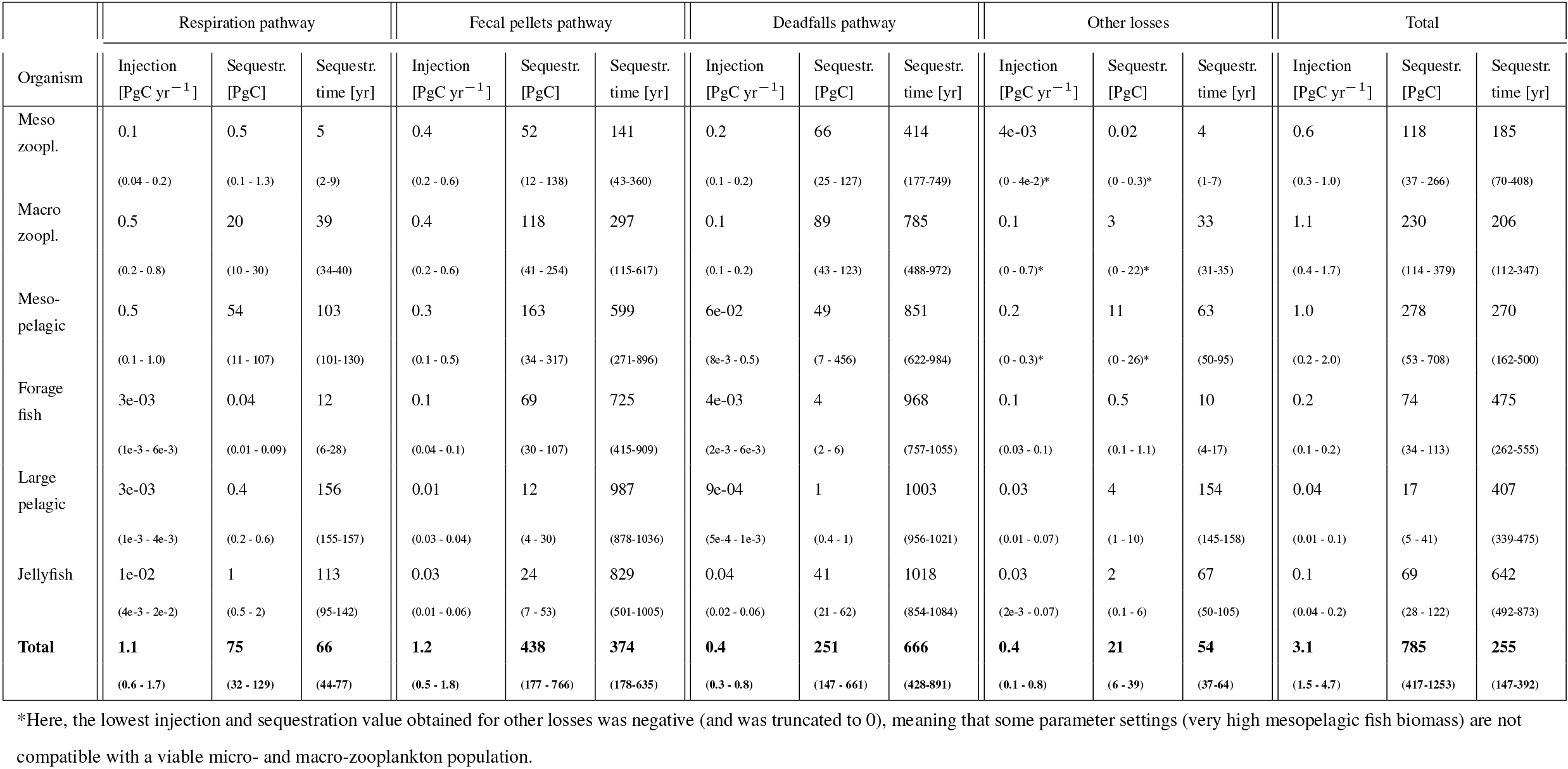
Total injection, corresponding sequestration, and sequestration time for the different pathways considered in the model. Respiration pathway corresponds to animal respiration, fecal pellets pathway to bacterial respiration of metazoan fecal pellets, deadfalls to natural mortality, and other losses to all other losses. The range is obtained from the different scenarios of the sensitivity analysis, reported in section S7.1.

Results from the game-theoretic model of vertical migration provides a global map of the day and night distribution of biomass of each metazoan group (figure 2). The predicted biomass-weighted mean depth (figure 2), taken as the model-predicted mean daytime depth of all fish weighted by biomass, is deeper in oceanic gyres (between 500-700 m deep), and shallower along the ocean margins and at the Equator (between 200-400 m). Predicted DVM patterns of the different functional groups (figure 2) can be compared to echosounder observations (figure 3a, (Klevjer et al., 2016; IMOS, 2021; Polar Data Centre British Antarctic Survey, 2020)). Even though low frequency (e.g. 38 kHz) echosounder observations can be biased (Proud et al., 2019), they can be used as a proxy for estimating the mean depth of water-column communities (figure 2 and S7). Our simulations generally match echosounder observations: meso-zooplankton and forage fish remain close to the surface, whereas macro-zooplankton and mesopelagic fish (as well as jellyfish) perform vertical migrations everywhere (Figure 2). At temperate latitudes, our model predicts shallower migrations than observed, in particular in the Southern Ocean where seasonality can lead to large annual variations in DVM behaviour (Prihartato et al., 2015) coupled to zooplankton dormancy (Bandara et al., 2016).

**Figure 3.**
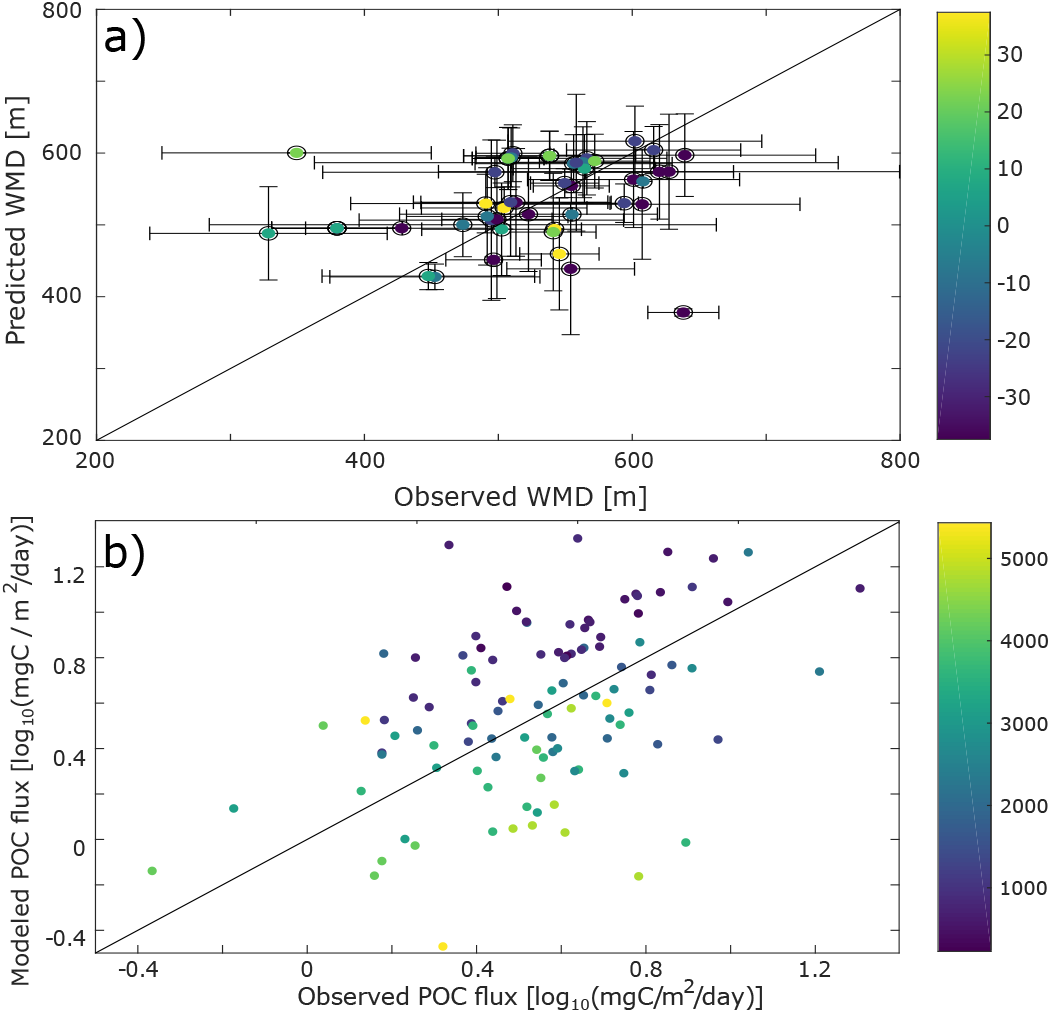
Comparison with data. a) Scatter plot of the differences between the observed and the model-predicted weighted mean depth (WMD) of the deep scattering layer. b) Difference between the observed (from sediment traps data, Lutz et al. (2007)) and modeled POC flux. To decrease possible biases due to localized blooms, only fixed sediment traps deeper than 500m and with an annual coverage were selected for this comparison.

We estimate the contributions of the different functional groups to carbon sequestration, as well as the corresponding residence times of respired carbon (figure 4 and table 1). Mesopelagic fish are the most important contributors to carbon se-questration with a total of 280 (50-710) PgC sequestered, followed by zooplankton (meso- and macro-zooplankton contribute 120 (40-270) and 230 (110-380) PgC respectively), forage fish (70 (30-110) PgC), and jellyfish (70 (30-120) PgC). Carbon sequestered via the fecal pellets pathway resides in the ocean longer than carbon sequestered via respiration (370 (180-640) years vs. 70 (40-80) years for all functional groups, table 1). In addition, carbon sequestered via degradation of fast-sinking fish fecal pellets or carcasses is stored on much longer time scales (up to 970 (760-1060) years for forage fish carcasses, and more than a thousand years for jellyfish and large pelagic fish carcasses) than carbon sequestered via degradation of slower-sinking fecal pellets such as meso-zooplankton (140 (40-360) years). While zooplankton produce the largest carbon fluxes globally, carbon injected via fish respiration and degradation of detritus originating from fish are stored more efficiently in the ocean’s interior – all because larger organisms tend to remain deeper and because they produce larger particles that sink faster and thus escape remineralization in the upper parts of the water columns.

**Figure 4.**
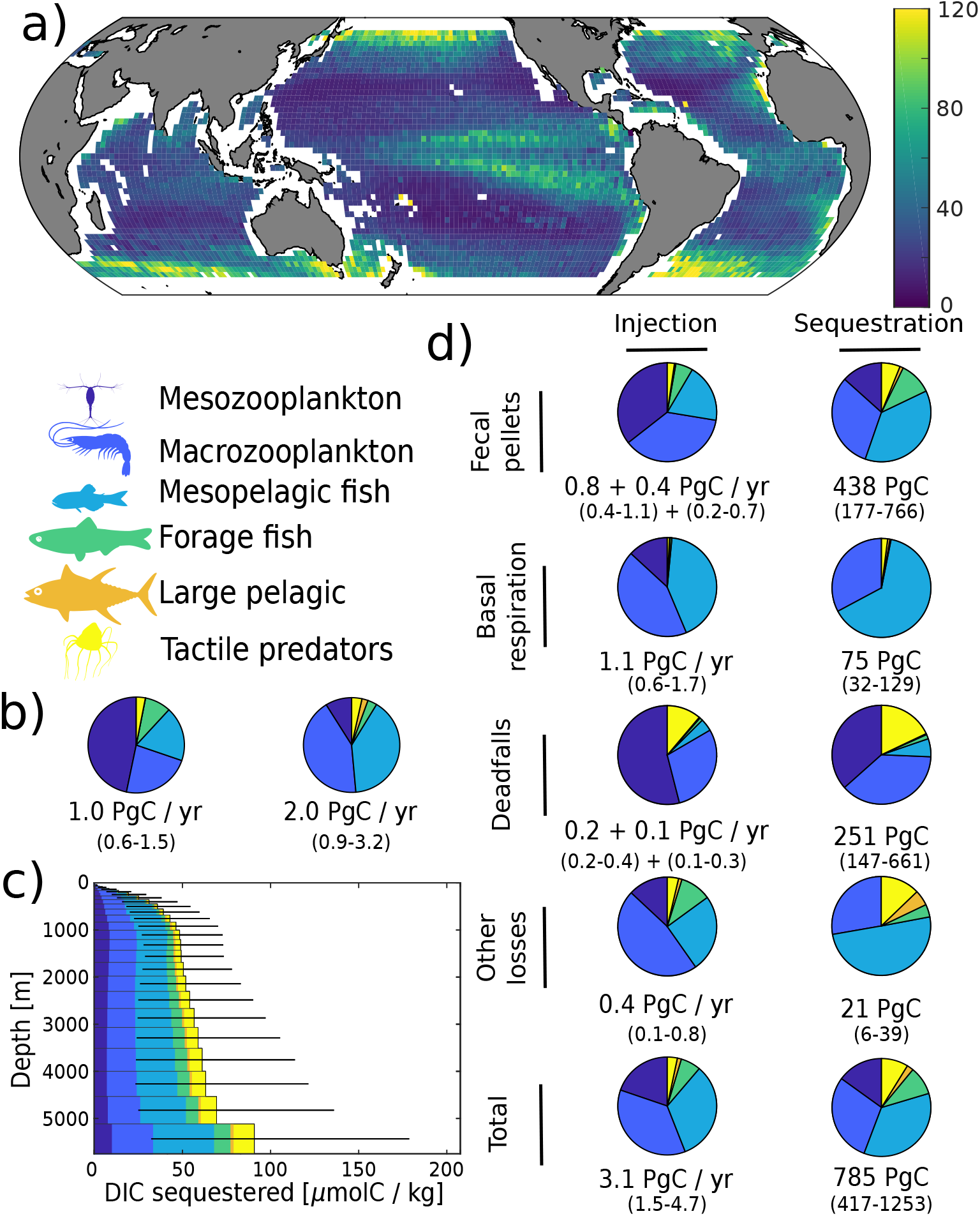
Simulated carbon injection and sequestration by metazoans. a) Simulated injection below the euphotic zone (in mgC m^−2^ day^−1^). b) Relative contribution of the simulated functional groups to injection below the euphotic zone. Left pie chart corresponds to the degradation of organic carbon that was produced in the euphotic zone and subsequently sank below the euphotic zone, right pie chart corresponds to the degradation of organic carbon that was injected by metazoans directly below the euphotic zone. c) Globally averaged concentration of DIC derived from the activity of metazoans according to the depth at which it is sequestered. d) Relative contribution of the functional groups to injection and sequestration (left and right column respectively) via the different pathways. For fecal pellets and carcass degradation, the first flux value corresponds to the degradation of organic carbon that was produced in the euphotic zone and subsequently sank below the euphotic zone, while the second corresponds to the degradation of organic carbon that was injected by metazoans directly below the euphotic zone. For basal respiration and other losses, only the direct injection below the euphotic zone is reported as there is no sinking flux for these pathways.

On a regional level, the absolute magnitude of carbon injected by metazoans below the euphotic zone varies significantly, from less than 10 to around 120 mgC m^−2^ day^−1^ (figure 4a). Subtropical gyres have the lowest injection, followed by the tropics, the Southern Ocean, the North Atlantic and North Pacific. The relative contribution of mesopelagic fish varies per geographic zone (figure S10), consistent with previous observations (Davison et al., 2013; Saba et al., 2021). Mesopelagic fish dominate carbon sequestration via the respiration pathway (more than 70% (30-80%) of the total) due to their deep daytime residence depths (figure 2).

Some aspects of our predictions can be compared to independent observations or constraints. We already commented on the DVM predictions. Our predictions of sinking particle fluxes can be compared to observations from sediment traps (figure 3b). While there are large differences between data and observations for some locations (up to 15 mgC m^−2^ day^−1^), the predicted fluxes are of the same order of magnitude as those observed. There is no global or regional bias in these differences (figure S9) and the depth bias in modeled vs. observed sediment trap flux is consistent with biases usually witnessed for this type of data (Schlitzer et al., 2003).

Oxygen demand places a strong constraint on how much carbon the oceans can sequester by the remineralization of organic matter (i.e. the primary path of the biological carbon pump). Predicted global ocean carbon sequestration constrained by apparent oxygen utilization (AOU) data is 1770 PgC across the global ocean, while a recent study taking into account variations in the concentration of oxygen subducted into the interior ocean (Carter et al., 2021) estimated that the interior ocean stores 1300 (± 230) PgC. Our estimate of 780 (420-1250) PgC for metazoans, certainly comes under this constraint, and can be compared directly to Nowicki et al. (2022)’s estimate for metazoans of 980 (830-1140), although that model only represented zooplankton and did not include fish or jellyfish. The simulated AOU values (SI; figure S13) show a deeper maximum than the observed AOU because we resolve processes with faster sinking speed, whereas remaining processes (e.g. remineralization of DOC, aggregates and small fecal pellets from micro-zooplankton) would be concentrated in the upper oceans. The difference arises because we do not consider all export pathways (e.g. phytoplankton and aggregate sinkings and DOC transport), and that our spatial coverage accounts for only 63% of the global ocean (no coastal areas nor latitudes higher than ±45^*°*^). Overall, our predictions of DVM, fluxes at depth, and AOU are compatible with available independent observations.

Because the large number of parameters and high computational cost of each simulation prohibit an exhaustive sensitivity analysis, we focused model sensitivity tests to nine poorly-constrained parameters: bacterial degradation rate of fecal pellets and carcasses, fecal pellet sinking speeds, biomass of mesopelagic fish, biomass of all functional groups, assimilation efficiencies, assimilation efficiency for detritus only, swimming speeds of all organisms, swimming speeds of mesopelagic fish only, and reference and maximum temperatures for all temperature-dependent rates. These parameters are anticipated to be those to which carbon injection and sequestration are most sensitive. Overall, the DVM patterns observed are robust (figure S14). Carbon injected and sequestered vary significantly between sensitivity scenarios, but are mostly of the same order as the ranges of the parameter variations (table S2-S22). This highlights the need to better understand mid-water animal ecology and to refine pelagic biomass estimates, in order to constrain these parameters more. In addition, we ran a more detailed Monte-Carlo sensitivity analysis for five different ecoregions (subtropical gyres, tropical area, North Pacific, North Atlantic and Southern Ocean). This analysis confirms that the behaviour of organisms and passive and active injections are relatively robust to changes in parameter values (figures S16-S20). Respiration due to other losses, and to a lesser degree the sinking flux below the euphotic zone, is more sensitive to small changes in parameters than basal respiration and the production of detritus (fecal pellets or carcasses) below the euphotic zone. The sensitivity to changes in parameter values was similar within ocean ecoregions (figures S16-S20).

## 4 Discussion

Present global estimates of total organic carbon export are roughly 10 PgC yr^−1^ (Schlitzer, 2002; Dunne et al., 2005; DeVries and Weber, 2017; Bisson et al., 2020; Siegel et al., 2023). Of this, the export production directly associated with POC aggregates from phytoplankton and unicellular organisms accounts for 1-2 PgC yr^−1^ (Siegel et al., 2014; Bisson et al., 2020; Nowicki et al., 2022), sinking of fecal pellets and carcasses from microzooplankton accounts for 3-4 PgC yr^−1^ (Bisson et al., 2020)), and the export of DOC by ocean mixing for another 1-2 PgC/yr (Hansell et al., 2009; Roshan and DeVries, 2017). Our estimate that metazoans contribute about 2.0 (1.2-3.0) PgC yr^−1^ to this export production (table S1) is consistent with the deficit, and appears to be mostly facilitated by macro-zooplankton and mesopelagic fish. Our results indicate that the metazoan contribution to export production is roughly evenly split between the production of POC (fecal pellets and carcasses) and active transport by migrants.

This export flux of organic carbon supplies pelagic and benthic ecosystems, where it is further transported and processed by bacteria and metazoans before it is eventually converted to dissolved inorganic carbon (DIC). We introduce the term “carbon injection” as the production of DIC via various respiratory pathways, and indicates when it becomes unavailable for further processing. Nearly all of the carbon exported from the surface is eventually injected into the ocean’s interior, but the depth at which it is injected can vary significantly depending on sinking speeds and how it is cycled through the various trophic links in Vinogradov’s ladder. Our results indicate that 3 (1.5 - 4.7) PgC yr^−1^ carbon injected into the oceans is mediated by metazoans.

Whether in terms of export production or carbon injection, the role of metazoans seems relatively modest (20% to 30% of global estimates). However, carbon sequestration by the biological pump is more than just export; it depends critically on how long respired carbon remains below the surface. It is in this respect, particularly through their vertical migrations, that metazoans occupy key roles in ocean carbon budgets. Our results indicate that roughly ~800 PgC of the total ~1300 PgC of sequestered respired DIC is due to metazoans, mainly by mesopelagic fish and macrozooplankton. This is consistent with the estimate of 1000 PgC (Nowicki et al., 2022) for zooplankton (both migrant respiration and fecal pellets), although that estimate included microzooplankton fecal pellets, and their model did not include fish nor jellyfish. Our study is able to disentangle the various pathways for sequestration by metazoans as ~60% by fecal pellets, ~30% by deadfall, and ~10% by basal respiration. Again this is consistent with Nowicki et al. (2022) who found about a ~5 fold difference between the contribution of migrant fecal pellets and respiration with regards their contribution to sequestration.

While our estimates carry considerable uncertainty, the relative contribution to carbon sequestration budgets are relatively robust. For instance, mesopelagic fish and macrozooplankton (e.g. krill) each contribute ~30% to total metazoan sequestration, a relative proportion also exhibited in their contribution to sequestration via the fecal pellet pathway. These two groups also constitute close to all of the sequestration via basal respiration, likely due to their near exclusive residency below the euphotic zone. Mesozooplankton (e.g. copepods) contribute modestly to sequestration via fecal pellets (~ 10%), but have a larger proportional impact on the deadfall pathway (~25%). Jellyfish contribute a relative minor fraction (~10%) to overall sequestration, but their deadfalls appear to be responsible for much greater carbon sequestration than those of pelagic or mesopelagic fish (Table 1).

Our results suggest that, despite large uncertainties, fish play an important role in the global carbon cycle – a hypothesis supported by local estimates (Saba and Steinberg, 2012; Davison et al., 2013; Hudson et al., 2014) and by an analysis of observed data in a recent review (Saba et al., 2021). Our model is not built on observations of DVM or carbon flux, but on fundamental mechanistic principles defining the interactions between individuals within different functional groups. These interactions lead to realistic vertical migration patterns and carbon fluxes that are coupled to a global ocean circulation model to assess global carbon sequestration inventories. In our model, fish (including mesopelagic fish, forage fish, and large pelagic fish) account for 40% (14-60%) of the carbon injected by metazoans below the euphotic zone. Compared to global carbon export due to all processes (i.e. including phytoplankton and microzooplankton) of around ~10 PgC yr^−1^ (Schlitzer, 2002; Dunne et al., 2005; DeVries and Weber, 2017), this suggests that fish are responsible for 12% (4-23%) of total export. This figure is in line with a recent literature review of local studies that estimated that fish were responsible for around 16% (± 13%) of carbon flux out of the euphotic zone (Saba et al., 2021). More importantly, our analysis suggests that fish are responsible for 50% (25-65%) of simulated carbon sequestration by metazoans (table 1). The large influence of fish on carbon sequestration (relative to export or injection) is due to the deep migration depths of mesopelagic fish, and the production of large fast-sinking fecal pellets, both of which lead to long sequestration times for the resulting respired DIC. While these first global mechanistic estimates of DVM patterns and fish carbon sequestration are subject to uncertainty, they provide a baseline for future assessments and for evaluating the carbon sequestration impact of fishing.

In addition to the passive sinking of fecal pellets and carcasses, our model also predicts carbon export by active diel vertical migration. Other modelling studies that have assessed the role of DVM on carbon export have relied on heuristics rather than mechanistic principles (Aumont et al., 2018; Archibald et al., 2019), and rarely consider functional groups separately to assess their relative importance (Archibald et al., 2019). Their resulting estimates mostly align with ours. Aumont et al. (2018) estimated that all migrating organisms export about 1.0 PgC yr^−1^ below 150 m (this depth is always deeper than the euphotic zone limit, so this result is hard to relate directly to ours), while Archibald et al. (2019) found that zooplankton are responsible for the export of about 0.8 PgC yr^−1^ below the euphotic zone. Global carbon export measurements estimate that mesopelagic fish are responsible for a carbon flux of 1.5 ±1.2 PgC yr^−1^ (Saba et al., 2021), a figure in agreement with our simulated injection of 1.0 (0.2-2.0) PgC yr^−1^. Note that here we are using carbon injection and not carbon export. Carbon injection is a more relevant metric when it comes to metazoan-driven carbon transport, as (organic) carbon exported can be uptaken again by detritivorous organisms, while carbon injected (in the form of dissolved inorganic carbon) cannot be reused by metazoans.

As we compute only one DVM cycle per geographical location, we neglect seasonality and assume that the ocean is at steady state. While this assumption is justified in tropical environments, especially when it comes to behaviours, this approach does not allow capturing peaks in export fluxes that can sometimes account for a significant fraction of the total annual export over short time periods. For example, at station M in the California Current System, 30 ± 10 % of the annual POC flux arrives at 3400m in episodic pulses 0 to 70 days after satellite-based estimates of maximum POC flux (Smith et al., 2018), with a single pulse reported to account for up to 44 % of the yearly POC flux at ~ 3900 m (Preston et al., 2020). Modelling the seasonality of pelagic environments in the water column would tremendously increase computing cost in a game theoretic setting with no guarantee of accurately picturing DVM behaviours as the model complexity would also increase – making it very sensitive to minor changes throughout the year, especially when episodic high pulses of primary production and POC flux arise throughout the year. This is why we decided not to include seasonality in our model and to provide baseline estimates of carbon export and sequestration instead.

Within this framework, our model results are relatively robust, as a factor 2 change in the most sensitive parameter values leads to a twofold change in export. The relative importance of fish for carbon sequestration remains high throughout the sensitivity analysis. However, more research is needed to validate these estimates. One of the most sensitive inputs of our model is biomass, and global estimates are highly uncertain. For example, mesopelagic fish global biomass estimates vary between 20 and 200% of the reference estimate due to the uncertainty in translating echosounder observations into biomass estimates (Proud et al., 2019). Other functional groups could not be included in the model because knowledge of their biology and abundance is even scarcer than for mesopelagic fish. Cephalopods, and squids in particular, occupy important mesopelagic niches. Depending on their sizes, they can feed on zooplankton or mesopelagic fish, and they are also valuable food resources for large predatory fish (Choy et al., 2013; Rodhouse, 2013). Their presence as a functional group in our model would probably slightly modify trophic couplings (Ariza et al., 2016), and, consequently, carbon export estimates.

In addition, gelatinous zooplankton estimates are still highly imprecise (Lucas et al., 2014), but potentially of considerable importance. A recent study (Luo et al., 2020) estimated that gelatinous zooplankton were responsible for a global export of 1.6-5.2 PgC yr^−1^ below 100 m. Even though that study included coasts and high latitudes and had a fixed depth horizon, their estimate is still much higher than our estimate of a total injection of 0.1 (0.04-0.2) PgC yr^−1^ below the euphotic zone for jellyfish, perhaps because their study –unlike this one– included gelatinous zooplankton that can also feed on phytoplankton and micro-zooplankton. Further, our model only considers one functional group for jellyfish, whereas in reality there is a vast diversity of gelatinous predators with a wide range of behaviours. When more data become available (i.e. global abundances and metabolic rates of salps, cnidarians, appendicularians), we will be able to refine the description of that group to gain better insight of the respective importance of these different taxa. Even though mixed strategies (i.e. a functional group with a sub-population performing DVM while another sub-population remains resident at depth) can emerge in our model, describing specifically these different functional groups will enable a better representation of the different behaviours that are observed in the ocean.

An omitted functional group of this model is bathypelagic fish. These fish constantly live below ~1000 m, potentially migrating daily between bathyal depths (up 4000 m deep) and the mesopelagic zone, taking up the lower rungs of Vinogradov’s ladder (Vinogradov, 1962). These organisms, feeding on mesopelagic fish (that can also, sometimes, migrate below 1000 m (Ariza et al., 2015; Hudson et al., 2014)), would tend to increase the time scales on which carbon is sequestered. The biomass of bathypelagic fish is, however, even less well known than the biomass of mesopelagic fish. Therefore, their potential contribution to global carbon sequestration is hard to assess. We can only conjecture that carbon sequestered because of bathypelagic fish respiration and excretion would be sequestered on very long time scales given the depths at which these organisms live. This consideration emphasizes further the importance of considering carbon injection and sequestration in addition to carbon export. While carbon export is an important metric, it only gives a partial idea of ocean carbon budgets. Carbon injection – the depth dependent biologically mediated source of DIC – is a more relevant metric that all biological pump studies should strive to estimate, whether focusing on the degradation of sinking POC (i.e. bacterial respiration) or respiration from vertical migrants.

As anthropogenic pressures increase, the last realm to remain relatively undisturbed by human activities is the deep sea. This may change because of commercial incentives to fish on the vast resource that mesopelagic fish represents (St. John et al., 2016). It has been suggested that 50% of the existing mesopelagic biomass can be sustainably extracted (St. John et al., 2016). However, fishing may have implications for carbon sequestration (Mariani et al., 2020; Bianchi et al., 2021), and there is a trade-off between economic gain of developing mesopelagic fishing and the cost of the forgone carbon sequestration.

Finally we note that the oxygen demand imposed by the biological pump in the ocean’s interior is not simply a useful constraint for estimating carbon sequestration; it is an issue of real concern. Simulated changes in ocean circulation and bio-geochemisty under climate change indicate a significant decrease in global ocean oxygen levels even with reduced export production (Koeve et al., 2020). Reduced oxygen will render increasing volumes of the ocean inaccessible to aerobic animals (Deutsch et al., 2011) and disrupt their vertical migrations with repercussions on the efficiency of the biological pump (sequestration time scale).

## Code availability

The source code (written in MATLAB) supporting this article has been uploaded as part of the supplementary material and is available at: https://gitlab.gbar.dtu.dk/jppi/global-fish-biological-carbon-pump

## Author contributions

JP designed the study with help from AWV and TK. RP, CMP, and ASB contributed biomass data. JP conducted the study with technical assistance from TDV, TN, CSP, DAS, TKA, KHA, and AWV. JP analysed results with help from TDV, DAS, CSP, TK, KHA, and AWV. JP wrote the manuscript with contributions from all authors. All authors approved the manuscript and agreed to be held personally accountable for their own contributions.

## Acknowledgements

This work was supported by the Centre for Ocean Life, a VKR Centre of excellence funded by the Villum Foundation, and by the Gordon and Betty Moore Foundation (grant #5479). ASB and RP were funded in part through the EU BG3 project ‘SUMMER’ and BG8 project ‘Mission Atlantic’. Collated echosounder data obtained from the BODC included observations made during the Atlantic Meridional Transect. The Atlantic Meridional Transect is funded by the UK Natural Environment Research Council through its National Capability Long-term Single Centre Science Programme, Climate Linked Atlantic Sector Science (grant number NE/R015953/1). This study contributes to the international IMBeR project and is contribution number 378 of the AMT programme.

## References

Archibald, K. M., Siegel, D. A., and Doney, S. C.: Modeling the impact of zooplankton diel vertical migration on the carbon export flux of the biological pump, Global Biogeochemical Cycles, 33, 181–199, 2019.

Ariza, A., Garijo, J. C., Landeira, J. M., Bordes, F., and Hernández-León, S.: Migrant biomass and respiratory carbon flux by zooplankton and micronekton in the subtropical northeast Atlantic Ocean (Canary Islands), Progress in Oceanography, 134, 330–342, https://doi.org/10.1016/j.pocean.2015.03.003, 2015.

Ariza, A., Landeira, J. M., Escánez, A., Wienerroither, R., Aguilar de Soto, N., Røstad, A., Kaartvedt, S., and Hernández-León, S.: Vertical distribution, composition and migratory patterns of acoustic scattering layers in the Canary Islands, Journal of Marine Systems, 157, 82–91, https://doi.org/10.1016/j.jmarsys.2016.01.004, 2016.

Aumont, O., Maury, O., Lefort, S., and Bopp, L.: Evaluating the Potential Impacts of the Diurnal Vertical Migration by Marine Organisms on Marine Biogeochemistry, Global Biogeochemical Cycles, 32, 1–22, https://doi.org/10.1029/2018GB005886, 2018.

Bandara, K., Varpe, Ø., Søreide, J. E., Wallenschus, J., Berge, J., and Eiane, K.: Seasonal vertical strategies in a high-Arctic coastal zooplankton community, Marine Ecology Progress Series, 555, 49–64, https://doi.org/10.3354/meps11831, 2016.

Bandara, K., Varpe, Ø., Wijewardene, L., Tverberg, V., and Eiane, K.: Two hundred years of zooplankton vertical migration research, Biological Reviews, https://doi.org/10.1111/brv.12715, 2021.

Bianchi, D., Carozza, D. A., Galbraith, E. D., Guiet, J., and DeVries, T.: Estimating global biomass and biogeochemical cycling of marine fish with and without fishing, Science Advances, 7, https://doi.org/10.1126/sciadv.abd7554, 2021.

Bisson, K., Siegel, D. A., and DeVries, T.: Diagnosing Mechanisms of Ocean Carbon Export in a Satellite-Based Food Web Model, Frontiers in Marine Science, 7, 1–15, https://doi.org/10.3389/fmars.2020.00505, 2020.

Boyd, P. W., Claustre, H., Levy, M., Siegel, D. A., and Weber, T.: Multi-faceted particle pumps drive carbon sequestration in the ocean, Nature, 568, 327–335, https://doi.org/10.1038/s41586-019-1098-2, 2019.

Buesseler, K. O. and Boyd, P. W.: Shedding light on processes that control particle export and flux attenuation in the twilight zone of the open ocean, Limnology and Oceanography, 54, 1210–1232, https://doi.org/10.4319/lo.2009.54.4.1210, 2009.

Canadell, J., Monteiro, P., Costa, M., Cotrim da Cunha, L., Cox, P., Eliseev, A., Henson, S., Ishii, M., Jaccard, S., Koven, C., Lohila, A., Patra, P., Piao, S., Rogelj, J., Syampungani, S., Zaehle, S., and Zickfeld, K.: Global Carbon and other Biogeochemical Cycles and Feedbacks, in: Climate Change 2021: The Physical Science Basis. Contribution of Working Group I to the Sixth Assessment Report of the Intergovernmental Panel on Climate Change, edited by Masson-Delmotte, V., Zhai, P., Pirani, A., Connors, S., Péan, C., Berger, S., Caud, N., Chen, Y., Goldfarb, L., Gomis, M., Huang, M., Leitzell, K., Lonnoy, E., Matthews, J., Maycock, T., Waterfield, T., Yelekçi, O., Yu, R., and Zhou, B., pp. 673–816, Cambridge University Press, Cambrige, UK and New York, NY, USA, 2021.

Carter, B. R., Feely, R. A., Lauvset, S. K., Olsen, A., DeVries, T., and Sonnerup, R.: Preformed Properties for Marine Organic Matter and Carbonate Mineral Cycling Quantification, Global Biogeochemical Cycles, 35, https://doi.org/10.1029/2020GB006623, 2021.

Choy, C. A., Portner, E., Iwane, M., and Drazen, J. C.: Diets of five important predatory mesopelagic fishes of the central North Pacific, Marine Ecology Progress Series, 492, 169–184, https://doi.org/10.3354/meps10518, 2013.

Davison, P. C., Checkley, D. M., Koslow, J. A., and Barlow, J.: Carbon export mediated by mesopelagic fishes in the northeast Pacific Ocean, Progress in Oceanography, 116, 14–30, https://doi.org/10.1016/j.pocean.2013.05.013, 2013.

Deutsch, C., Brix, H., Ito, T., and Thompson, L.: Climate-forced variability of ocean hypoxia, Science, 333, 336–340, 2011.

DeVries, T.: The oceanic anthropogenic CO2 sink: Storage, air-sea fluxes, and transports over the industrial era, Global Biogeochemical Cycles, 28, 631–647, https://doi.org/10.1002/2013GB004739, 2014.

DeVries, T. and Primeau, F.: Dynamically and observationally constrained estimates of water-mass distributions and ages in the global ocean, Journal of Physical Oceanography, 41, 2381–2401, https://doi.org/10.1175/JPO-D-10-05011.1, 2011.

DeVries, T. and Weber, T.: The export and fate of organic matter in the ocean: New constraints from combining satellite and oceanographic tracer observations, Global Biogeochemical Cycles, 31, 535–555, https://doi.org/10.1002/2016GB005551, 2017.

Dunne, J. P., Armstrong, R. A., Gnnadesikan, A., and Sarmiento, J. L.: Empirical and mechanistic models for the particle export ratio, Global Biogeochemical Cycles, 19, 1–16, https://doi.org/10.1029/2004GB002390, 2005.

Garcia, H., Weathers, C., Paver, C., Smolyar, I., Boyer, T., Locarnini, R., Zweng, M., Mishonov, A., Baranova, O., Seidov, D., and Reagan, J.: WORLD OCEAN ATLAS 2018 Volume 3: Dissolved Oxygen, Apparent Oxygen Utilization, and Dissolved Oxygen Saturation, Tech. rep., Silver Spring, MD, https://www.nodc.noaa.gov/OC5/woa18/pubwoa18.html, 2019.

Gorgues, T., Aumont, O., and Memery, L.: Simulated changes in the particulate carbon export efficiency due to diel vertical migration of zooplankton in the North Atlantic, Geophysical Research Letters, p. 2018GL081748, https://doi.org/10.1029/2018GL081748, 2019.

Hansell, D., Carlson, C., Repeta, D., and Schlitzer, R.: DISSOLVED ORGANIC MATTER IN THE OCEAN: A CONTROVERSY STIMULATES NEW INSIGHTS, Oceanography, 22, 202–211, http://www.jstor.org/stable/24861036, 2009.

Hansen, A. N. and Visser, A. W.: Carbon export by vertically migrating zooplankton: An optimal behavior model, Limnology and Oceanography, 61, 701–710, https://doi.org/10.1002/lno.10249, 2016.

Hernández-León, S., Olivar, M. P., Fernández de Puelles, M. L., Bode, A., Castellón, A., López-Pérez, C., Tuset, V. M., and González-Gordillo, J. I.: Zooplankton and Micronekton Active Flux Across the Tropical and Subtropical Atlantic Ocean, Frontiers in Marine Science, 6, https://doi.org/10.3389/fmars.2019.00535, 2019.

Hofbauer, J. and Sigmund, K.: Evolutionary Game Dynamics, Bulletin (New Series) of the American mathematical society, 40, 479–519, 2003.

Holland, K. N., Brill, R. W., Chang, R. K., Sibert, J. R., and Fournier, D. A.: Physiological and behavioural thermoregulation in bigeye tuna (Thunnus obesus), Nature, 358, 410–412, https://doi.org/10.1038/358410a0, 1992.

Holling, C.: Some characteristics of simple types of predation and parasitism, The Canadian Entomologist, 91, 385–398, 1959.

Holzer, M., DeVries, T., and de Lavergne, C.: Diffusion controls the ventilation of a Pacific Shadow Zone above abyssal overturning, Nature Communications, 12, 1–13, https://doi.org/10.1038/s41467-021-24648-x, 2021.

Houston, A. I., McNamara, J. M., and Hutchinson, J. M. C.: General results concerning the trade-off between gaining energy and avoiding predation, Philosophical Transactions of the Royal Society of London. Series B: Biological Sciences, 341, 375–397, 1993.

Hudson, J. M., Steinberg, D. K., Sutton, T. T., Graves, J. E., and Latour, R. J.: Myctophid feeding ecology and carbon transport along the northern Mid-Atlantic Ridge, Deep-Sea Research Part I: Oceanographic Research Papers, 93, 104–116, https://doi.org/10.1016/j.dsr.2014.07.002, 2014.

Hugie, D. M. and Dill, L. M.: Fish and Game: a game theoretic approach to habitat selection by predators and prey, Journal of Fish Biology, 45, 151–169, 1994.

IMOS: IMOS BASOOP sub facility, imos.org.au, 2021.

Irigoien, X., Klevjer, T. A., Røstad, A., Martinez, U., Boyra, G., Acuña, J. L., Bode, A., Echevarria, F., Gonzalez-Gordillo, J. I., Hernandez-Leon, S., Agusti, S., Aksnes, D. L., Duarte, C. M., and Kaartvedt, S.: Large mesopelagic fishes biomass and trophic efficiency in the open ocean, Nature communications, 5, 3271, https://doi.org/10.1038/ncomms4271, 2014.

Jónasdóttir, S. H., Visser, A. W., Richardson, K., and Heath, M. R.: Seasonal copepod lipid pump promotes carbon sequestration in the deep North Atlantic, Proceedings of the National Academy of Sciences, 112, 12 122–12 126, https://doi.org/10.1073/pnas.1512110112, 2015.

Klevjer, T. A., Irigoien, X., Røstad, A., Fraile-Nuez, E., Benítez-Barrios, V. M., and Kaartvedt, S.: Large scale patterns in vertical distribution and behaviour of mesopelagic scattering layers, Scientific Reports, 6, 19 873, 2016.

Koeve, W., Kähler, P., and Oschlies, A.: Does Export Production Measure Transient Changes of the Biological Carbon Pump’s Feedback to the Atmosphere Under Global Warming?, Geophysical Research Letters, 47, https://doi.org/10.1029/2020GL089928, 2020.

Lampert, W.: Ultimate causes of diel vertical migration of zooplankton: New evidence for the predator-avoidance hypothesis, Archiv her Hydrobiologie, Beiheft Ergebnisse der Limnologie, 39, 79–88, https://pure.mpg.de/pubman/faces/ViewItemOverviewPage.jsp?itemId=item{_}1508686http://www.crcnetbase.com/doi/10.1081/E-EEE2-120046011, 1993.

Locarnini, R., Mishonov, A., Baranova, O., Boyer, T., Zweng, M., Garcia, H., Reagan, J., Seidov, D., Weathers, K., Paver, C., and Smolyar, I.: World Ocean Atlas, Volume 1: Temperature, http://www.nodc.noaa.gov/OC5/indprod.html, 2019.

Longhurst, A., Bedo, A., Harrison, W., Head, E., and Sameoto, D.: Vertical flux of respiratory carbon by oceanic diel migrant biota, Deep Sea Research Part A. Oceanographic Research Papers, 37, 685–694, https://doi.org/10.1016/0198-0149(90)90098-G, 1990.

Lucas, C. H., Jones, D. O., Hollyhead, C. J., Condon, R. H., Duarte, C. M., Graham, W. M., Robinson, K. L., Pitt, K. A., Schildhauer, M., and Regetz, J.: Gelatinous zooplankton biomass in the global oceans: Geographic variation and environmental drivers, Global Ecology and Biogeography, 23, 701–714, https://doi.org/10.1111/geb.12169, 2014.

Luo, J. Y., Condon, R. H., Stock, C. A., Duarte, C. M., Lucas, C. H., Pitt, K. A., and Cowen, R.: Gelatinous zooplankton-mediated carbon flows in the global oceans: a data-driven modeling study, Global Biogeochemical Cycles, 34, 2020.

Lutz, M. J., Caldeira, K., Dunbar, R. B., and Behrenfeld, M. J.: Seasonal rhythms of net primary production and particulate organic carbon flux to depth describe the efficiency of biological pump in the global ocean, Journal of Geophysical Research: Oceans, 112, https://doi.org/10.1029/2006JC003706, 2007.

Mariani, G., Cheung, W. W., Lyet, A., Sala, E., Mayorga, J., Velez, L., Gaines, S. D., Dejean, T., Troussellier, M., and Mouillot, D.: Let more big fish sink: Fisheries prevent blue carbon sequestration-half in unprofitable areas, Science advances, 6, 1–9, https://doi.org/10.1126/sciadv.abb4848, 2020.

McLaren, I. A.: Effects of Temperature on Growth of Zooplankton, and the Adaptive Value of Vertical Migration, Journal of the Fisheries Research Board of Canada, 20, 685–727, https://doi.org/10.1139/f63-046, 1963.

Nash, J.: Non-Cooperative Games, The Annals of Mathematics, 54, 286, https://doi.org/10.2307/1969529, 1951.

Nowicki, M. E., Devries, T., and Siegel, D. A.: Quantifying carbon export and sequestration pathways in the global ocean, Global Biogeochemical Cycles, 36, e2021.B007 083, 2022.

Petrik, C. M., Stock, C. A., Andersen, K. H., van Denderen, P. D., and Watson, J. R.: Bottom-up drivers of global patterns of demersal, forage, and pelagic fishes, Progress in Oceanography, 176, 102 124, https://doi.org/10.1016/j.pocean.2019.102124, 2019.

Pinti, J. and Visser, A. W.: Predator-Prey Games in Multiple Habitats Reveal Mixed Strategies in Diel Vertical Migration, The American Naturalist, 193, E65–E77, 2019.

Pinti, J., Kiørboe, T., Thygesen, U. H., and Visser, A. W.: Trophic interactions drive the emergence of diel vertical migration patterns: a game-theoretic model of copepod communities, Proceedings of the Royal Society B: Biological Sciences, 286, 20191 645, 2019.

Pinti, J., Andersen, K. H., and Visser, A. W.: Co-adaptive behavior of interacting populations in a habitat selection game significantly impacts ecosystem functions, Journal of Theoretical Biology, p. 110663, https://doi.org/10.1016/j.jtbi.2021.110663, 2021.

Polar Data Centre British Antarctic Survey: British Antarctic Survey, Raw acoustic data collected by ship-borne EK60 echosounder in the Atlantic Ocean (AMT24, AMT25, AMT26, AMT29)., 2020.

Preston, C. M., Durkin, C. A., and Yamahara, K. M.: DNA metabarcoding reveals organisms contributing to particulate matter flux to abyssal depths in the North East Pacific ocean, Deep-Sea Research Part II: Topical Studies in Oceanography, 173, 104 708, https://doi.org/10.1016/j.dsr2.2019.104708, 2020.

Prihartato, P. K., Aksnes, D. L., and Kaartvedt, S.: Seasonal patterns in the nocturnal distributionand behavior of the mesopelagic fish Maurolicus muelleri at high latitudes, Marine Ecology Progress Series, 521, 189–200, 2015.

Proud, R., Cox, M. J., and Brierley, A. S.: Biogeography of the Global Ocean’s Mesopelagic Zone, Current Biology, 27, 113–119, https://doi.org/10.1016/j.cub.2016.11.003, 2017.

Proud, R., Handegard, N. O., Kloser, R. J., Cox, M. J., and Brierley, A. S.: From siphonophores to deep scattering layers: uncertainty ranges for the estimation of global mesopelagic fish biomass, ICES Journal of Marine Science, 76, 718–733, https://doi.org/10.1093/icesjms/fsy037, 2019.

Rodhouse, P. G.: Role of squid in the Southern Ocean pelagic ecosystem and the possible consequences of climate change, Deep-Sea Research Part II: Topical Studies in Oceanography, 95, 129–138, https://doi.org/10.1016/j.dsr2.2012.07.001, 2013.

Roshan, S. and DeVries, T.: Efficient dissolved organic carbon production and export in the oligotrophic ocean, Nature Communications, 8, 1–8, https://doi.org/10.1038/s41467-017-02227-3, 2017.

Saba, G. K. and Steinberg, D. K.: Abundance, composition, and sinking rates of fish fecal pellets in the santa barbara channel, Scientific Reports, 2, 1–6, https://doi.org/10.1038/srep00716, 2012.

Saba, G. K., Burd, A. B., Dunne, J. P., Hernández-león, S., Martin, A. H., Rose, K. A., Salisbury, J., Steinberg, D. K., Trueman, C. N., Wilson, R. W., and Wilson, S. E.: Toward a better understanding of fish-based contribution to ocean carbon flux, Limnology and Oceanography, pp. 1–26, https://doi.org/10.1002/lno.11709, 2021.

Sainmont, J., Thygesen, U. H., and Visser, A. W.: Diel vertical migration arising in a habitat selection game, Theoretical Ecology, 6, 241–251, https://doi.org/10.1007/s12080-012-0174-0, 2013.

Sainmont, J., Andersen, K. H., Thygesen, U. H., Fiksen, Ø., and Visser, A. W.: An effective algorithm for approximating adaptive behavior in seasonal environments, Ecological Modelling, 311, 20–30, 2015.

Schlitzer, R.: Carbon export fluxes in the Southern Ocean: Results from inverse modeling and comparison with satellite-based estimates, Deep-Sea Research Part II: Topical Studies in Oceanography, 49, 1623–1644, https://doi.org/10.1016/S0967-0645(02)00004-8, 2002.

Schlitzer, R., Usbeck, R., and Fischer, G.: Inverse modeling of particulate organic carbon fluxes in the South Atlantic, in: The South Atlantic in the Late Quaternary: Reconstruction of material budgets and current systems, edited by Wefer, G., Mulitza, S., and Ratmeyer, V., pp. 1–19, Springer-Verlag, https://doi.org/10.1007/978-3-642-18917-3, 2003.

Siegel, D. A., Buesseler, K. O., Doney, S. C., Sailley, S. F., Behrenfeld, M. J., and Boyd, P. W.: Global assessment of ocean carbon export by combining satellite observations and food-web models, Global Biogeochemical Cycles, 28, 181–196, https://doi.org/10.1002/2013GB004743, 2014.

Siegel, D. A., DeVries, T., Cetinić, I., and Bisson, K. M.: Quantifying the Ocean’s Biological Pump and Its Carbon Cycle Impacts on Global Scales, Annual Review of Marine Science, 15, null, https://doi.org/10.1146/annurev-marine-040722-115226, pMID: 36070554, 2023.

Smith, K. L., Ruhl, H. A., Huffard, C. L., Messié, M., and Kahru, M.: Episodic organic carbon fluxes from surface ocean to abyssal depths during long-term monitoring in NE Pacific, Proceedings of the National Academy of Sciences of the United States of America, 115, 12 235–12 240, https://doi.org/10.1073/pnas.1814559115, 2018.

St. John, M. A., Borja, A., Chust, G., Heath, M. R., Grigorov, I., Mariani, P., Martin, A. P., and Santos, R. S.: A Dark Hole in Our Understanding of Marine Ecosystems and Their Services: Perspectives from the Mesopelagic Community, Frontiers in Marine Science, 3, 1–6, https://doi.org/10.3389/fmars.2016.00031, 2016.

Steinberg, D. K., Carlson, C. A., Bates, N. R., Goldthwait, S. A., Madin, L. P., and Michaels, A. F.: Zooplankton vertical migration and the active transport of dissolved organic and inorganic carbon in the Sargasso Sea, Deep Sea Research Part I, 47, 137–158, https://doi.org/10.1016/S0967-0637(99)00052-7, 2000.

Stock, C. A., Dunne, J. P., and John, J. G.: Global-scale carbon and energy flows through the marine planktonic food web: An analysis with a coupled physical-biological model, Progress in Oceanography, 120, 1–28, https://doi.org/10.1016/j.pocean.2013.07.001, 2014.

Stock, C. A., John, J. G., Rykaczewski, R. R., Asch, R. G., Cheung, W. W., Dunne, J. P., Friedland, K. D., Lam, V. W., Sarmiento, J. L., and Watson, R. A.: Reconciling fisheries catch and ocean productivity, Proceedings of the National Academy of Sciences of the United States of America, 114, E1441–E1449, https://doi.org/10.1073/pnas.1610238114, 2017.

Thygesen, U. H., Sommer, L., Evans, K., and Patterson, T. A.: Dynamic optimal foraging theory explains vertical migrations of Bigeye tuna, Ecology, 97, 1852–1861, 2016.

Titelman, J. and Hansson, L. J.: Feeding rates of the jellyfish Aurelia aurita on fish larvae, Marine Biology, 149, 297–306, https://doi.org/10.1007/s00227-005-0200-5, 2006.

Vinogradov, M.: Feeding of the deep-sea zooplankton, ICES reports, 18, 114–120, 1962.

Zaret, T. M. and Suffern, S.: Vertical migration in zooplankton as a predator avoidance mechanism, Limnology and Oceanography, 21, 804–813, https://doi.org/10.4319/lo.1976.21.6.0804, 1976.

