## Supplementary material for "Model estimates of metazoans’ contributions to the biological carbon pump"

### Supplementary material
### The global importance of fish to the biological carbon pump

Jérôme Pinti, Tim DeVries, Tommy Norin, Camila Serra-Pompei, Roland Proud, David A. Siegel, Thomas Kiørboe, Colleen M. Petrik, Ken H. Andersen, Andrew S. Brierley, André W. Visser

**2 Global model** **10**

**3 Carbon injection and sequestration** **14**

**4 Difference between carbon export and injection** **15**

**5 Additional results** **16**

**7 Glossary of parameters** **39**

### 1 1D diel vertical migration model

#### 1.1 Food-web set-up

The model depicts a pelagic community, from surface waters to mesopelagic depths. We consider phytoplankton resources, two zooplankton populations (meso-zooplankton and macro-zooplankton), forage fish, large pelagic fish, tactile predators (e.g. jellyfish), mesopelagic fish, and detritus (figure 1). The water column is discretized in $n$ layers. The average total biomass of all functional groups in the water column is fixed. The vertical distribution of phytoplankton is fixed and distributed in the surface mixed layer. Large pelagic fish are very fast organisms able to move up and down the water column several times every day [18, 49], so we consider their distribution fixed and uniform in the water column. The vertical distribution of all other groups (meso- and macro-zooplankton, forage fish, mesopelagic fish and tactile predators) is the emergent property of the model, as organisms can perform DVM. The day is divided in two periods of time – daytime and nighttime – and organisms can choose their daytime and nighttime positions according to fitness-optimization rules detailed below. Detritus is created by organisms (through fecal pellet and carcasses production), sinks and is degraded along the way, but can also be ingested by macro-zooplankton.

A summary of all parameters and functions used in this document is provided tables S23, S24, S25, S26 and S27.

We call $\tilde{X}$ the mean concentration of population $X$ in the water column in $\text{gC m}^{-3}$. Here and in the rest of this document, $X$ is a placeholder referring to the different groups of individuals considered: meso-zooplankton $C$, macro-zooplankton $P$, forage fish $F$, mesopelagic fish $M$, large pelagic organisms $A$ or tactile predators $J$. $X_{ij}$ is the proportion of population $X$ following strategy $ij$ (we call strategy the set of day ($i$)-night ($j$) positions an individual adopts). In the following of this document, $i$ and $j$ will either refer to a water layer or to the depth of this specific water layer. By definition, we have

$$\sum_{i=1}^{n}\sum_{j=1}^{n} X_{ij} = 1. \tag{1}$$

The concentration of organisms in water layer $i$ during daytime is

$$X(i, day) = \tilde{X} n \sum_{k=1}^{n} X_{ik}, \tag{2}$$

with a similar expression for the concentration of organisms in layer $j$ during nighttime.

In the water column, abiotic conditions (temperature, light levels, oxygen concentration) vary vertically, impacting the fitness $W_X$ of organisms. Light levels also change between day and night, creating the possibility for organisms to perform DVM – if the optimal strategy is to change vertical position during day and night. The goal of performing DVM is to optimize fitness. As an individual selects a strategy, the fitness of its prey, predators and conspecifics also varies. Hence, the optimal strategy of each individual is intrinsically linked to the optimal strategy of the other players. The optimal strategies for all individuals is attained at the Nash equilibrium [32], where no individual can increase its fitness by changing its strategy. The Nash equilibrium is found using the replicator equation [17, 36, see section 1.5]. In short, the fraction of the population following a particular strategy grows proportionally to the fitness related to that strategy, before renormalization to ensure fixed population sizes.

#### 1.2 Effects of temperature and oxygen on standard metabolic rate, maximum metabolic rate and aerobic scope

While the effect of temperature $T$ on vital rates is fairly well understood and modelled (e.g. with $Q_{10}$ temperature coefficients), oxygen concentration is rarely considered in population models of zooplankton and fish. The way oxygen concentration affects the standard metabolic rate (SMR) and the maximum metabolic rate (MMR) of organisms depends on whether they are oxygen regulators or oxygen conformers (figure S1). Oxygen regulators (i.e. fish) have an SMR depending only on temperature [13, 41] – at least at oxygen concentrations above the critical oxygen tension ($p_{crit}$), below which the animal will start accumulating an oxygen debt (see also section 1.2.3) and eventually die if continually exposed to hypoxia or anoxia – whereas oxygen conformers (i.e. zooplankton and jellyfish) have the ability to decrease their oxygen requirements when the oxygen partial pressure $p_{O_2}$ drops below $p_{reg}$ (typically below 60% oxygen

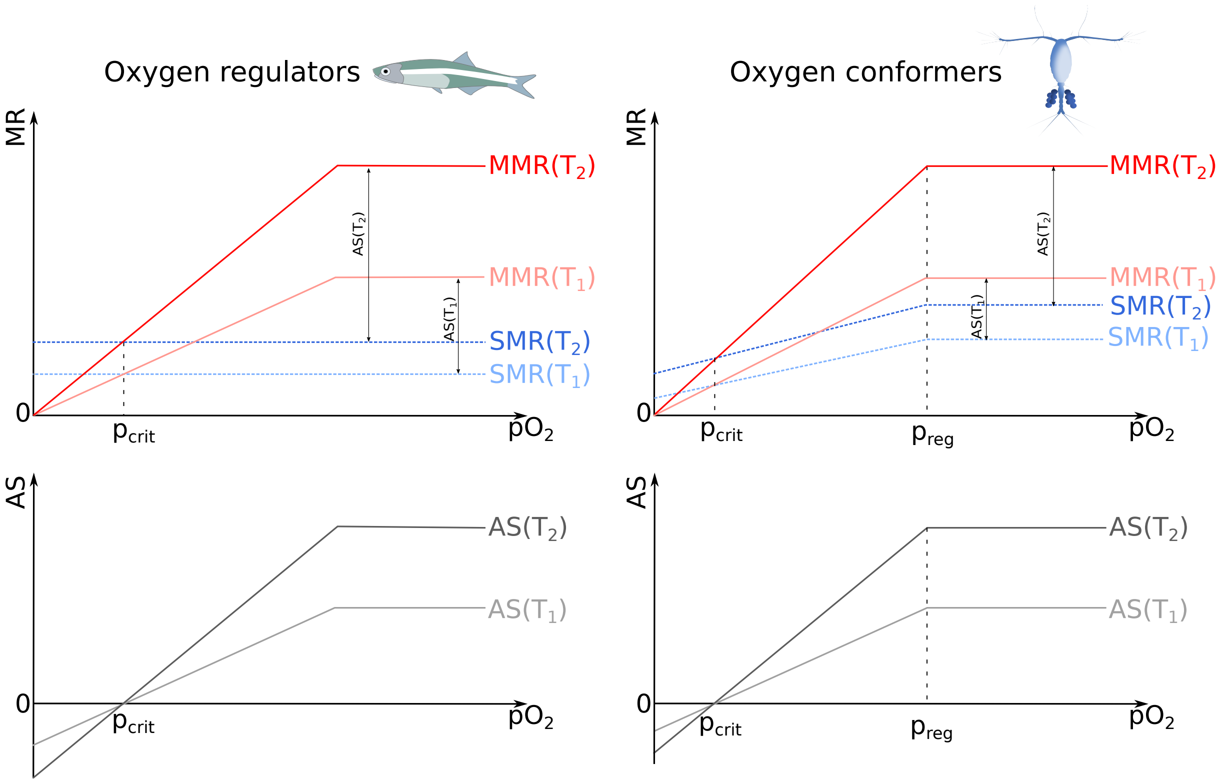

Figure S1: Outline of typical maximum metabolic rates (MMR), standard metabolic rates (SMR) and aerobic scopes (AS=MMR-SMR) of oxygen regulator and oxygen conformer organisms. Here, $T_1 < T_2$.

saturation, [23]). For both oxygen regulators and conformers, the MMR drops when $p_{O_2}$ decreases below $p_{reg}$ [45].

$p_{crit}$ is the oxygen partial pressure at which MMR equals SMR, and $p_{reg}$ the partial pressure above which the MMR cannot increase.

The aerobic scope $AS$ (defined as MMR-SMR [14, 8]) defines the amount of oxygen that organisms can use for their activities. As such, we assume that maximum ingestion rate and activities such as swimming speed scale linearly with it, up to a maximum level. Few expressions of aerobic scope of pelagic fish or zooplankton as a function of both temperature and oxygen are available in the literature, and for simplicity we transformed the expression of Claireaux et al. [8] in a piece-wise linear function (figure S1) for both oxygen conformers regulators.

##### 1.2.1 Rates for oxygen conformers

In the following, we drop the dependencies in $X$, $T$ and $p_{O_2}$ for readability. We also call $s(T)$ the function:

$$s(T) = SMR_0 Q_{10}^{\frac{min(T,T_{max})-T_{ref}}{10}}, \tag{3}$$

where $T$ is temperature, $SMR_0$ the standard metabolic rate at $T_{ref}$, and $T_{max}$ the temperature of maximum aerobic scope.

The following equations are derived from the maximum metabolic rate being equal to the standard metabolic rate at $p_{crit}$ (by definition of $p_{crit}$) and from the maximum metabolic rate being 0 under complete anoxia (figure S1, [50]).

The SMR is:

$$\begin{aligned} SMR &= s(T) & \text{if } p_{O_2} > p_{reg} = 0.6 p_{max} \\ &= s(T)\left[1 + \left(\Delta_{MR}\frac{p_{crit}}{p_{reg}} - 1\right)\frac{p_{reg}-p_{O_2}}{p_{reg}-p_{crit}}\right] & \text{if } p_{O_2} \leq p_{reg}, \end{aligned} \tag{4}$$

with $p_{O_2}$ the oxygen partial pressure and $\Delta_{MR}$ the factor of increase of MMR compared to SMR under normoxia.

Similarly, the MMR of an oxygen conformer is:

$$\begin{aligned} MMR &= \Delta_{MR} s(T) && \text{if } p_{O_2} > p_{reg} = 0.6 p_{max} \\ &= \Delta_{MR} s(T) \frac{p_{O_2}}{p_{reg}} && \text{if } p_{O_2} \leq p_{reg}. \end{aligned} \tag{5}$$

##### 1.2.2 Rates for oxygen regulators

With the same conventions as section 1.2.1, the SMR is (figure S1):

$$SMR = s(T), \tag{6}$$

and the maximum metabolic rate is:

$$MMR = \min\left( s(T) \frac{p_{O_2}}{p_{crit}}, \Delta_{MR} s(T) \right). \tag{7}$$

##### 1.2.3 Diel context

For some combinations of temperature and oxygen concentration, the aerobic scope of the organism is negative (figure S1), meaning that it has a deficit in oxygen supply compared to its oxygen demand.

Organisms can build up and sustain an oxygen debt (typically building lactates [44, 46]) for a period of time. Fish can usually sustain this oxygen deficiency for a few minutes to a few hours [47, 33], whereas zooplankton can withstand such low oxygen conditions for several hours and up to half a day [13]. As the hypoxic tolerance of fish is below the temporal resolution of our model, habitats yielding negative metabolic scopes (and their related strategies) are not available to fish populations (figure S2), except large pelagic fish that are scattered in the water column.

Some zooplankton can, however, maintain an anaerobic metabolism for a period of up to half a day. When in anaerobic mode, they build up an oxygen debt that needs to be repaid later. In our case, we consider that the oxygen debt taken during one part of the day (daytime or nighttime) will be repaid during the other part of the day, decreasing the available aerobic scope. Once again, the aerobic scope is set to a minimum of 0, so oxygen debts cannot be repaid in all cases: for example, a day and a night residency at low oxygen levels is not a viable strategy as it would create a oxygen debt that can never be repaid (not to mention the fact that a constant aerobic scope of 0 means that the organism does not feed, see figure S2). In cases where the oxygen debt can be repaid (for example a day oxygen debt repaid at night), the available aerobic scope during nighttime $S_X(i, j, day = 0)$ is equal to its original value as calculated in sections 1.2.1 and 1.2.2 plus the negative daytime aerobic scope $\tilde{S_X}(i, j, day = 1) = AS(z = i)$, modulated by the proportion of daylight hours in a day:

$$S_X(i, j, 0) = (1 - \sigma)\tilde{S_X}(i, j, 0) + \sigma \tilde{S_X}(i, j, 1). \tag{8}$$

#### 1.3 Fitness

The fitness $W$ of an individual from population $X$ following strategy $ij$ (i.e. being at depth $i$ during day and $j$ during night) is defined, following Gilliam's rule, as its growth rate divided by its mortality rate [16, 19]:

$$W_X(i, j) = \frac{g_X(i, j)}{m_X(i, j)}. \tag{9}$$

The fitness of phytoplankton and of large predatory fish is not considered here as we constrain their vertical distributions.

Growth $g$ is equal to the assimilation rate $\nu$ due to predation minus a standard metabolic cost $Q$ and a migration cost $C_{migr}$. Mortality $m$ is due to predation $\mu$ and to a background mortality $\mu_0$.

##### 1.3.1 Metabolic rate

The standard metabolic cost experienced by an individual $X$ following strategy $ij$ is:

$$Q_X(i, j) = \sigma \tilde{Q_X}(i) + (1 - \sigma)\tilde{Q_X}(j), \tag{10}$$

with $\sigma$ the proportion of daylight hours in a day and $\tilde{Q}(z)$ the metabolic cost experienced at depth $z$. $\tilde{Q}(z)$ is the standard metabolic rate at the conditions encountered at depth $z$:

$$\tilde{Q}(z) = SMR(T(z), p_{O_2}(z)). \tag{11}$$

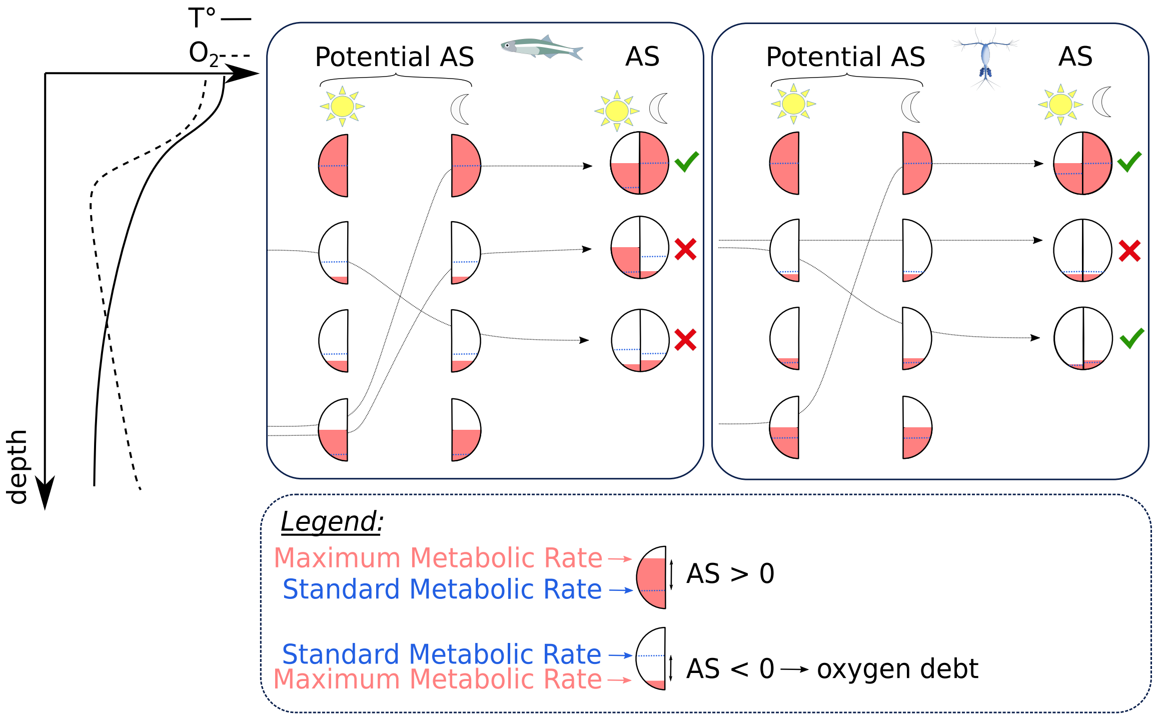

Figure S2: Example of possible (green check marks) and impossible (red crosses) strategies (arrows) for copepods and fish. Red level in half-circles represent the oxygen available, and the blue line the basal oxygen requirements, i.e. the standard metabolic rate. As depth increases, the standard metabolic rate decreases, and the maximum metabolic rate (and hence aerobic scope) scope changes because of changing oxygen and temperature levels. The SMR also decreases because of decreasing temperatures. When the potential aerobic scope gets negative, zooplankton can build up an oxygen debt which can be repaid during the other part of the day, but fish cannot – making any habitat with a negative aerobic scope unsuitable. For zooplankton, the total balance cannot be negative: the oxygen debt contracted during part of the day has to be repaid during the other part of the day.

The reference metabolic rate at $T_{ref}$ is $SMR_0$, defined as [25]:

$$SMR_0 = 0.0014 w_X^{-0.25}, \tag{12}$$

with $w_X$ the body mass measured in gC. Weights of organisms from each functional group are given in table S25.

##### 1.3.2 Assimilation rate

As for the metabolic rate, the total assimilation rate is the mean of the assimilation rate during day and night:

$$\nu_X(i,j) = \sigma\tilde{\nu}_{X,i,j}(i) + (1-\sigma)\tilde{\nu}_{X,i,j}(j), \tag{13}$$

with $\tilde{\nu}_{X,i,j}(z)$ the assimilation rate of $X$ at depth $z$ ($z = i$ during day and $j$ during night). Note that the calculation of $\tilde{\nu}$ is tedious as it depends on the feeding mode of the predator, on the concentration of prey at the considered depth, and on the environmental conditions encountered during both day and night – as the oxygen debt during part of the day can modulate the aerobic scope during the other part of the day and hence the assimilation rate.
The assimilation rate $\tilde{\nu}(z)$ is related to the swimming speed $u$ and the maximum ingestion rate $I_{max}$ of an individual, both of which are impacted by temperature and oxygen concentration in the water column as they scale linearly with the aerobic scope $S$ of the organism (section 1.2.3). We express $u$ and $I_{max}$ as:

$$\begin{aligned} u_{X,i,j}(z) &= u_{0,X}\frac{S_X(i,j,z)}{S_0}, \\ I_{max,X,i,j}(z) &= I_{max0,X}\max\left[10^{-10}, \frac{S_X(i,j,z)}{S_0}\right], \end{aligned} \tag{14}$$

with $u_{X0}$ and $I_{max,X0}$ being swimming speed and maximum ingestion with an aerobic scope $S_0$.

The reference swimming speed $u_{0,X}$ in m day$^{-1}$ is defined as [20]:

$$\begin{aligned} u_{0,X} &= 7.87 \cdot 10^4 l^{0.825} \text{ for zooplankton and fish,} \\ &= 2.68 \cdot 10^4 l^{0.75} \text{ for tactile predators,} \end{aligned} \tag{15}$$

with $l$ in m. Lengths of organisms from each functional group are given in table S25. The reference maximum ingestion rate in gC day$^{-1}$ is [2]:

$$\begin{aligned} I_{max0,X} &= 0.0542 w_X^{3/4} \text{ for zooplankton,} \\ &= 226.6 l_X^{2.55} \text{ for fish,} \end{aligned} \tag{16}$$

with $l$ in m and $w$ in gC.

The assimilation rate $\tilde{\nu}(z)$ can first be expressed as:

$$\tilde{\nu_X}(z) = \sum_{preyY} \varphi_X F_X^Y(z), \tag{17}$$

with $\varphi$ the assimilation efficiency and $F_X^Y(z)$ the ingestion rate of prey $Y$ by predator $X$ at depth $z$. Note that for macro-zooplankton, prey also include the detritus pool.

**Ingestion rate:**

The ingestion rate $F_X^Y(z)$ is defined as:

$$F_X^Y(z) = \frac{1}{w_X}\frac{I_{maz,X}(z)E_X^Y(z)}{I_{maz,X}(z) + \sum_{preyY'} E_X^{Y'}(z)}, \tag{18}$$

with $E_X^Y$ the encounter rate of prey $Y$ by predator $X$. $E_X^Y$ is equal to:

$$E_X^Y(day,z) = \Gamma_X^Y(z,day)V_X^Y(day,z)\Phi(X,Y)Y(day,z), \tag{19}$$

where $\Gamma_X^Y$ is the depth- and time-dependent capture probability of $Y$ by $X$, $V_X^Y$ the clearance rate of $X$ on $Y$, $\Phi$ the preference matrix (whose values are given table S26) and $Y$ the concentration of prey organisms at the time and depth considered. As in eq. 8 and for the rest of this document, *day* is a boolean taking value 1 during daytime and 0 during nighttime.

Large pelagic fish are uniformly distributed, as they can move up and down in the water column several times a day [18, 49]. As these organisms do not remain at fixed (different) depths all the time, their population is assumed to be monomorphic, i.e. all organisms within the population perform the same strategy. As such, the ingestion rate of large pelagic fish is:

$$F_A^Y(z) = \frac{1}{w_A} \frac{I_{max,A}(z) E_A^Y(z)}{I_{max,A}(z) + \int_0^{Z_{MAX}} \sum_{preyY'} E_A^{Y'}(z')\,\mathrm{d}z'}. \tag{20}$$

**Clearance rate:**

For fish, the clearance rate depends on the light level at the time and depth considered. The visual range $\Lambda_X^Y(z, day)$ of a fish $X$ preying on $Y$ is expressed as [3]:

$$\Lambda_X^Y(z, day) = R_{0,X}^Y \sqrt{\frac{light(z, day)}{K_{e,X} + light(z, day)}}, \tag{21}$$

with $R_{0,X}$ the reference visual range, $K_{e,X}$ the half-saturation constant for light and $light(z, day)$ the light level at the time and depth considered. Light levels are expressed as:

$$light(z, day) = \rho_l(day) L_{max} \exp(-\kappa z), \tag{22}$$

with $\rho_l$ the attenuation coefficient between day and night (1 and day, $10^{-5}$ at night), $L_{max}$ the surface daytime irradiance and $\kappa$ the light attenuation coefficient of the water. We assume that the visual range of a fish is always at least 10% of its body length. The clearance rate is the volume of water swept looking for prey per unit of time. For fish, it is:

$$V_X^Y(day, z) = \gamma\pi \max\left[\Lambda_X^Y(z, day)^2, (0.1 l_X)^2\right] u_X(z), \tag{23}$$

with $\gamma$ the cross-sectional area efficiently scanned and $u_X$ the predator swimming speed.

For zooplankton, the clearance rate is:

$$V_C^Y(day, z) = \pi(2l_C)^2 u_C(z), \tag{24}$$

with $2l_C$ the detection distance [51]. Finally, for tactile predators, the clearance rate is [1]:

$$V_J^Y(day, z) = f\pi(l_J/2)^2 u_J(z), \tag{25}$$

with $f$ the filtering efficiency of tactile predators.

**Capture probability:**

The capture probability $\Gamma_X^Y(z, day)$ is based on the maximum swimming speed of organisms and on the prey visual range [6, see figure S3]. During an attack event, the organisms do not swim at their cruising speeds $u$ but at their maximum swimming speed $u_{max}$, defined in m s$^{-1}$ as [12]:

$$u_{max} = \begin{cases} 1.15 w_c^{0.16} & \text{for zooplankton and fish} \\ 0.51 w_c^{0.16} & \text{for jellyfish,} \end{cases} \tag{26}$$

with $w$ in gC.

Once a predator $X$ encounters a prey $Y$, the attack event starts when the distance between the two is the prey detection distance $r_{detec} = \Lambda_Y^X(z, day)$ – if the prey detects the predator before the predator detects the prey, we assume that the prey always escapes before the predator detects it, and as such it is not captured. Similarly, the capture probability of non-motile prey (phytoplankton, detritus) is 1. For simplicity, we do not consider social behaviours and assume that capture probabilities are independent of prey and predator concentrations. The predator jumps towards the prey, and the prey tries to escape by jumping a distance $r_{esc}$ into a random direction. For simplicity, we assume here that the prey jumps for as long as it takes the predator to reach its initial position, so we have:

$$r_{esc} = \frac{r_{detec}}{AS_X(z) u_{max,X}} AS_Y(z) u_{max,Y}. \tag{27}$$

The predator jumps until it reaches the end of the sphere of all possible escape jumps of the prey (figure S3), so:

$$r_{attac} = r_{detec} + r_{esc}. \tag{28}$$

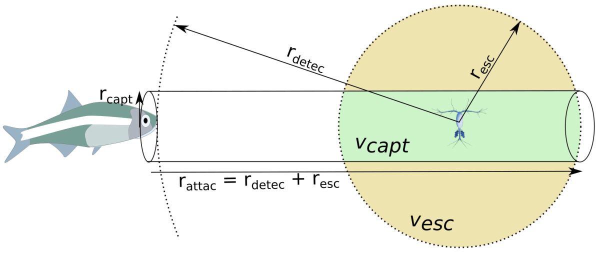

Figure S3: The zooplankton capture probability by the fish is the ratio between the green volume (intersection between the predator capture area and the zooplankton potential escape location) and the beige volume (sphere of potential escapes of the zooplankton).

The predator can capture the prey on a disc of radius $r_{capt} = 0.1 l_X$. As the prey can jump in any direction with no preference *a priori*, we assume that the capture probability is the fraction $v_{capt}$ of the escape sphere $v_{esc}$ swept by the predator as it moves through it:

$$\Gamma_X^Y(z, day) = \frac{v_{capt,X}^Y(z, day)}{v_{esc,X}^Y(z, day)}. \tag{29}$$

##### 1.3.3 Mortality

For all migrating populations, mortality is the sum of a predation mortality rate $\mu_X$ and of a background mortality rate $\mu_{0,X}$. For macro zooplankton and mesopelagic fish, $\mu_{0,X}$ is light-dependent to mimic extra predation risks from non-modelled functional groups. As for metabolic and assimilation rates, the total mortality rate of an individual following a strategy $ij$ is the mean of the mortality rates experienced during day and during night:

$$\mu_{Y,i,j} = \sigma \tilde{\mu}_Y(i, 1) + (1 - \sigma)\tilde{\mu}_Y(j, 0), \tag{30}$$

with $\tilde{\mu}_Y(z, day)$ the mortality rate experienced by $Y(i, j)$ during day (depth $i$) or during night (depth $j$). Note that, contrary to ingestion rates, mortality rates are not dependent on the aerobic scope of the individual, but only on the aerobic scopes of its predators. Therefore, the mortality rate of a prey population during $day$ at $z$ is:

$$\tilde{\mu}_Y(z, day) = \sum_{pred.X} \sum_{\substack{compatible\\ strat.ij}} F_X^Y X_{ij} \frac{1}{Y(z, day)}, \tag{31}$$

$Y(z, day)$ being the concentration of prey $Y$ at depth $z$ during $day$. The compatible strategies are the strategies that overlap with prey at the time and depth considered (so being at $i$ during daytime, or a $j$ during nighttime depending on the value of $day$).

##### 1.3.4 Migration cost

The migration cost $C_{migr,X}$ of an organism $X$ is calculated similarly to Pinti et al. [35], where organisms are simply assumed to be steadily translating spheres. We start by computing the hydrodynamic drag $D_{r,X}$ of the body [24]:

$$D_{r,X} = \frac{1}{2}\pi C_D \rho \frac{((10^{-2}\ \text{m cm}^{-1})l)^2}{4} u_X^2 \qquad [\text{kg m s}^{-2}], \tag{32}$$

with $\rho$ the density of the fluid, $u_X$ the speed of the organism and $C_{D,X}$ its drag coefficient. We can relate this drag coefficient to the Reynolds number $Re_X$ [24]:

$$C_{D,X} = \frac{24}{Re_X} + \frac{5}{\sqrt{Re_X}} + \frac{2}{5}, \tag{33}$$

with the Reynolds number defined as:

$$Re_X = \frac{10^{-2} l_X u_X}{\nu_w}. \tag{34}$$

$\nu_w$ is the kinematic viscosity of sea water. The normalized energy spent migrating is then:

$$\begin{aligned} C_{migr,X} &= 2\frac{1}{w_X}\frac{1}{46\cdot 10^3}\int_{migr\ event}\frac{D_{r,X}}{\epsilon}u_X\mathrm{d}t \\ &= \frac{D_{r,X}}{46\cdot 10^3\epsilon w_X}2\Delta Z \qquad [\mathrm{day}^{-1}]. \end{aligned} \tag{35}$$

$\epsilon$ is the efficiency with which internal energy is converted to motion [52] and $\Delta Z$ is the distance migrated. The factor 2 is there because the migration cost accounts for the migration at dawn and at dusk. The migration cost is converted to grams of carbon using a generic ratio of 46 kJ gC$^{-1}$ [42].

#### 1.4 Detritus

In this model, detritus are fecal pellets created by organisms which are consumed by detrivores or degraded by bacteria. Each population creates fecal pellets of different sizes and sinking speeds. $D_X(z)$ refers to the concentration at depth $z$ of fecal pellets created by population $X$. Fecal pellets are the part of the food ingested that is not assimilated, and is therefore created at the rate

$$D_{crea,X}(z) = (1-\varphi_X)\sum_{preyY} F_X^Y(z). \tag{36}$$

Large pelagic fish are assumed to be distributed evenly in the water column as a closure term, the even distribution means that fecal pellets or respiration could originate from depths where the organisms spend little or virtually no time in reality (so as at 1000 m depth). As a pragmatic solution to this problem, we redistribute the carbon respired and excreted at the depths at which large pelagic prey are present (forage fish, mesopelagic fish and tactile predators). Therefore, we have:

$$D_{crea,A}(z) = \quad (1-\varphi_A)\int_0^{Z_{MAX}}\sum_{preyY}F_X^Y(z')dz'\frac{\sigma\sum_{X=M,F,J}X(z,1)+(1-\sigma)\sum_{X=M,F,J}X(z,0)}{\int_0^{Z_{MAX}}\sigma\sum_{X=M,F,J}X(z',1)+(1-\sigma)\sum_{X=M,F,J}X(z',0)dz'} \tag{37}$$

Macro-zooplankton organisms can also feed on detritus, and their consumption rate of detritus is

$$D_{conso,X}(z) = F_P^{D_X}(z). \tag{38}$$

It is worth noting that in our model with no population or detritus dynamics (i.e. populations do not get depleted as they are eaten), the daily consumption of detritus in a water layer can be larger than than the concentration of detritus in that layer. As a pragmatic solution to this artefact, we impose a limit $\psi$ of the proportion of the detritus in a water layer that can be consumed daily:

$$\tilde{F}_P^{D_X}(z) = \min\left[\frac{\psi D_X(z)}{D_{conso,X}(z)}F_P^{D_X}(z), F_P^{D_X}(z)\right]. \tag{39}$$

Further, if a rescaling is used for macro-zooplankton detritus ingestion rates, the rescaling factor $\frac{\psi D_X(z)}{D_{conso,X}(z)}$ is also applied to detritus encounter rates, so that it modifies the macro-zooplankton ingestion rate of phytoplankton and meso-zooplankton (section 1.3.2 eq. 18).

All this considered, the creation of detritus $\zeta^X(z)$ is:

$$\zeta^X(z) = D_{crea,X}(z) - \tilde{F}_P^{D_X}(z). \tag{40}$$

The steady state concentration of detritus $D^X$ in the water column is obtained by solving the following transport equation:

$$\frac{\partial D^X}{\partial t} = -\alpha(z)D^X - \omega_X\frac{\partial D^X}{\partial z} + \zeta^X = 0, \tag{41}$$

where $\omega_X$ is the sinking speed of fecal pellets and $\alpha$ is the depth-dependent bacterial degradation rate of detritus:

$$\alpha(z) = \alpha_0 Q_{bac}^{\frac{T-T_{ref}}{10}}\frac{p_{O_2}}{K_{O_2}+p_{O_2}}, \tag{42}$$

with $\alpha_0$ the maximum degradation rate, $Q_{bac}$ the $Q_{10}$ factor of bacterial respiration and $K_{O_2}$ the half-saturation constant of the oxygen dependency of the degradation rate [11].

Eq. 41 is solved numerically using an Euler scheme.

##### 1.5 Nash equilibrium

The optimal vertical migration patterns of all organisms is attained when the system is at its Nash equilibrium [32]. At this point in the strategy space, no organism can increase its fitness by changing unilaterally its behaviour. Only a subset of the available strategies (i.e. set of depths $ij$) may be populated at this point, and all the populated strategies of a population will have identical fitness. Formally, this translates in:

$$\begin{aligned} W_X(i,j) &= W_X^0 \text{ for all } \textit{(i,j)} \text{ such that } X_{i,j} > 0, \\ W_X(i,j) &\leq W_X^0 \text{ for all } \textit{(i,j)} \text{ such that } X_{i,j} = 0. \end{aligned} \tag{43}$$

The Nash equilibrium of the system is found using the replicator equation [43]. The replicator equation is a two-step process, during which we allow the proportion of individuals following strategy $ij$ of a population to grow proportionally to its fitness, before renormalisation to ensure that populations do not grow in size:

$$\begin{aligned} X'_{ij}(t+\delta t) &= X_{ij}(t) + \lambda_t W_X(i,j) X_{i,j}(t), \\ X_{ij}(t+\delta t) &= \frac{X'_{ij}(t+\delta t)}{\sum_k \sum_l X'_{kl}(t+\delta t)}. \end{aligned} \tag{44}$$

$\lambda_t$ is a factor chosen to ensure a rapid transition to equilibrium. As a practical compromise, $\lambda_t$ is chosen so that $\lambda_t \max(W_X(i,j)) = 0.05$.

At each time step, the detritus flux and concentrations are also updated, and taken as the weighted mean of the equilibrium flux and concentrations given by the population distribution at time $t$ (90%) and $t + \delta t$ (10%).

### 2 Global model

In order to get global DVM patterns of the considered population, we run the previous model on the global ocean discretised in 1x1 degree cells. The water column model is run independently in each cell, ignoring possible interactions between the different cells, though physical properties and plankton biomasses do take into account the influence of currents (interacting cells), whether from in situ measurements (WOA) or an Earth System Model (COBALT).

To run the model on a global scale, physical and biological inputs are needed:

- Global 3D temperature field ;
- Global 3D oxygen concentration field ;
- Global surface irradiance and light attenuation coefficient ;
- Global biomasses (not resolved vertically – but only horizontally) of the different populations of the model: phytoplankton and microzooplankton, meso-zooplankton, macro-zooplankton, forage fish, mesopelagic fish, large pelagic fish and jellyfish ;
- Global mixed layer depth estimate, to assess the vertical distribution of phytoplankton in the water column.

We ignore seasonal fluctuations and use only annually averaged inputs. The global temperature and oxygen fields are averaged from the world ocean atlas [27, 15]. The light attenuation coefficient comes from the MODIS GMIS-AQUA climatology of the European Commission [30]. These three physical parameters were averaged over the period 2003-2017.

The day surface irradiance is calculated for each latitude following Naraghi and Etienne [31]:

$$L_{max} = \frac{1}{365} G_{SG} \sum_{d=1}^{365} \sin(\beta) \left[ 1 + 0.033 \cos(360 \frac{d-3}{365}) \right], \tag{45}$$

with $G_{SG}$ the solar constant equal to 1367 $W/m^2$, $d$ the day number of the year and $\beta$ the solar altitude angle, defined as

$$\sin(\beta) = \cos(lat)\cos(\delta)\cos(h) + \sin(lat)\sin(\delta), \tag{46}$$

with $lat$ the latitude considered, $h$ the hour angle (equal to 0 here as we are looking for the irradiance at solar midday), and $\delta$ the solar inclination angle equal to $23.45 \sin(360 \frac{d+284}{365})$. The resulting yearly-averaged surface irradiance is pictured figure S4.

Mixed layer depth is calculated using a temperature criterion of $\pm$ 0.2 degrees from temperature at 10 $m$ [9, figure S4]. The mixed layer depth is calculated to assess the distribution of the non-migrating phytoplankton in the water column. If $z_0$ is the mixed layer depth, phytoplankton are distributed following [22]:

$$R(z) \propto 1 - \tanh\left(\frac{4(z - z_0)}{z_0}\right). \tag{47}$$

The global estimates of resources and zooplankton are the outputs of the COBALT model [48] (mesozooplankton and macrozooplankton correspond to the medium and large zooplankton size classes in COBALT), that are also used in the FEISTY model that provides the abundances of forage fish and large predators [34]. Fish in FEISTY do not interact between grid cells, but the zooplankton and detritus they experience in their individual grid cells is the result of advection-diffusion in COBALT.

The predicted global median mesopelagic fish biomass of 3.8 Pg [39] (wet weight, assuming all fish retain gas-filled swimbladders throughout their life-cycles) is distributed proportionally by mesopelagic province [38] using the province values of predicted 38 kHz mesopelagic echo energy (i.e. the predicted total amount of echo-energy backscattered by the mesopelagic community when insonified using an echosounder operating at 38 kHz):

$$\text{Mesopelagic fish province biomass} = \frac{(\text{total biomass}) \cdot (\text{province 38 kHz mesopelagic echo energy})}{(\text{global 38 kHz mesopelagic echo energy})}. \tag{48}$$

Global uncertainty in mesopelagic fish biomass is about 1 order of magnitude, but local uncertainty cannot be quantified for the time being. Local NASC uncertainty would have to be combined with local species assemblage uncertainty. Estimating local variations in mesopelagic fish biomass estimates would require knowledge on siphonophore densities, depth distributions and gas bladder size distributions, and knowledge on fish swimbladder size distribution. Proud et al. [40] summarise current knowledge level on that issue.

For forage fish we use output from the FEISTY model [34]. FEISTY does not distinguish between forage and mesopelagic fish as it does not have vertical migrations implemented. Forage fish are only those feeding in the top 100m of the water column, and as such could also account for some mesopelagic fish – resulting in overestimating the importance of forage fish for carbon export. In practice, this risk is limited as mesopelagic fish biomass estimates are typically much higher than forage fish biomass estimates from FEISTY (figure S5).

The global abundance of jellyfish is likely to vary, both temporally and spatially. As such, we set their biomass to a constant and conservative estimate of $0.1\,\text{gC m}^{-2}$ [28]. Global biomasses of zooplankton and fish are pictured figure S5.

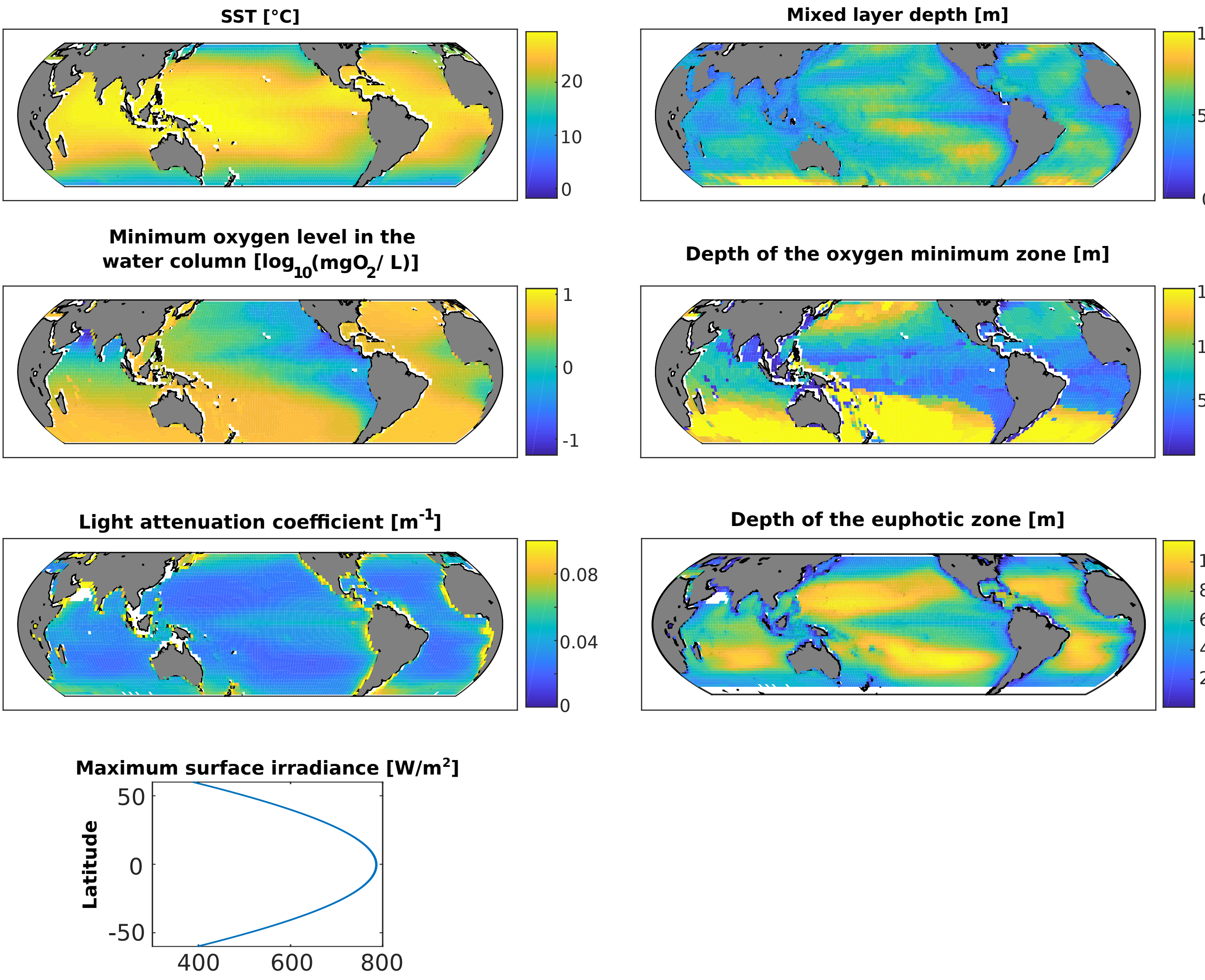

Figure S4: Physical inputs for the global model. In this figure, the depth of the oxygen minimum zone is capped to 1500 $m$ and the light attenuation coefficient to 0.1 $day^{-1}$.

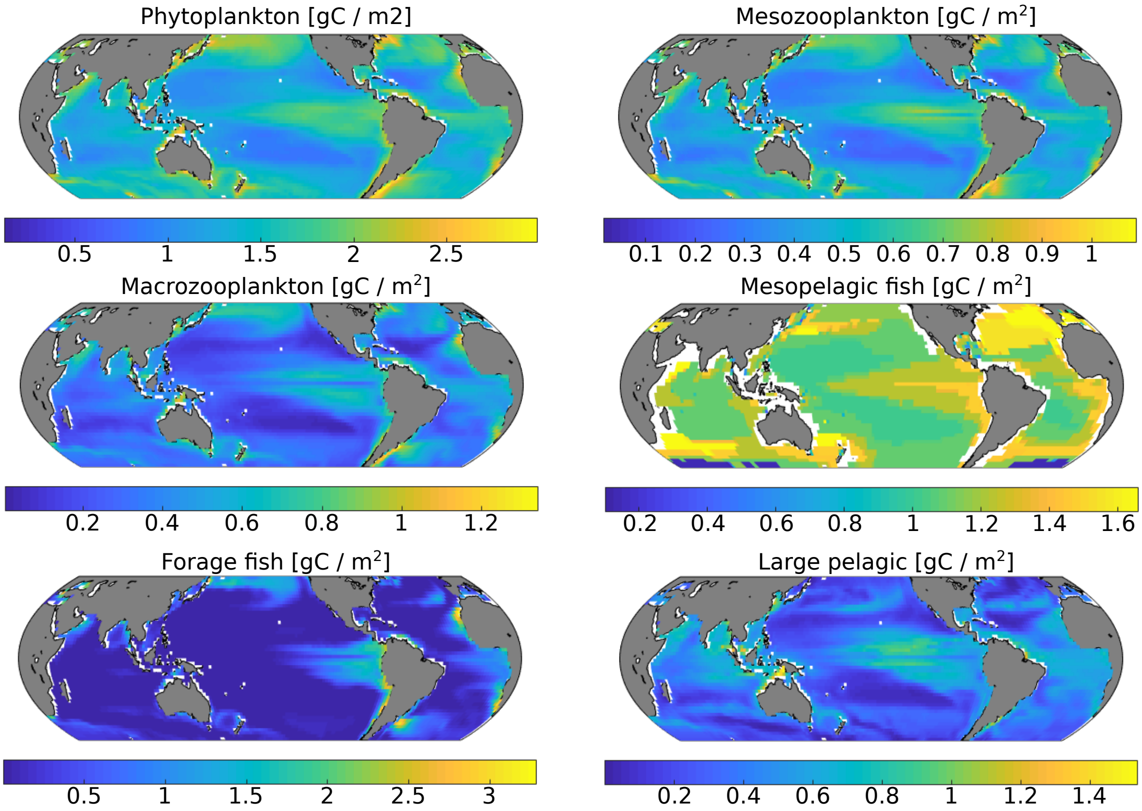

Figure S5: Biomasses of the different functional groups considered in gC / m$^2$.

#### 3 Carbon injection and sequestration

For each functional group, carbon can be sequestered in the oceans via 4 pathways: via fecal pellet degradation, via basal respiration, via carcass degradation (carcasses are produced at a rate equal to the background mortality of each functional group $\mu_{0,X}$), and via other losses (a term accounting for all processes not included in the model, including reproduction and specific dynamic action). The carbon budget is balanced in the model for each functional group, through the "other losses" pathway that acts as a buffer encompassing all processes unaccounted for explicitly in the model (e.g. expenditure due to reproduction, social behaviours). As a conservative estimate, we assume that these other losses are respired by organisms – a reasonable assumption when considering that carbon not currently considered in the bioenergetic budget would either be respired or used for reproduction. They are calculated, for all functional groups as the energy left when all metabolic and mortality costs are paid: $g_X(i,j) - m_X(i,j)$, in gC. This term can be negative, meaning in this case that the parametrization is such that the concerned population is not viable in the model as it suffers a permanent energy deficit. In practice, this happens only at the extremes of the parameter range and then indicates that the combination of parameters is not viable for the population in focus.

The global model outputs provide us with the 3D distribution of organisms, along with their ingestion, egestion and respiration rates. This enables us to compute how much carbon is respired by organisms in the global ocean, and how much carbon is egested as fecal pellets, as well as the natural mortality rate for organisms of all functional groups. Assuming the sinking rate and degradation rates of detritus (table S25 and eq. 42), we compute the flux of fecal material below any depth, but also the amounts of fecal material and carcasses that are turned into DIC (Dissolved Inorganic Carbon) by bacterial respiration (i.e. carbon injection).

The depth of the water column in the global model is set to 1000 m, and we extend here the water column to the *real* depth of the seafloor at the location considered. Detrital particles then sink (and get degraded) below 1000 m and down to the seafloor. We consider that all material reaching the seafloor is respired. Figure S6 shows the globally average DIC source from respiration generated by the bacterial respiration of sinking fecal pellets and by animal respiration.

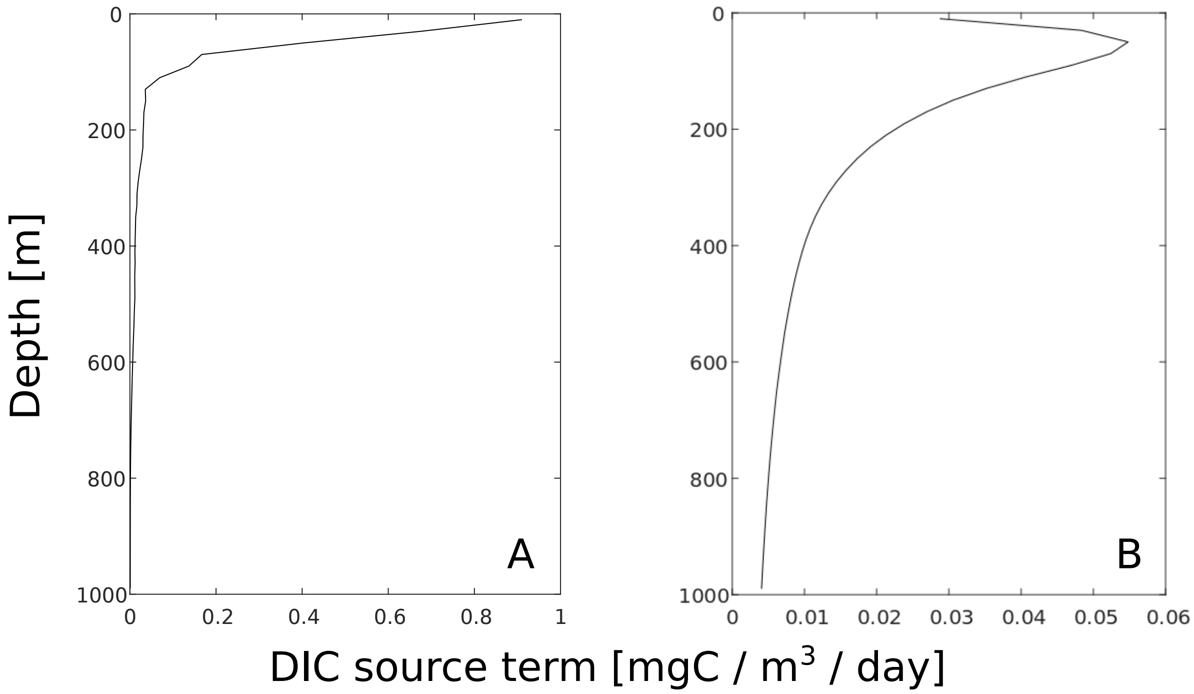

Figure S6: Globally averaged DIC production from (A) animal respiration and (B) bacterial respiration due to fecal pellets degradation.

We couple the DIC production terms to OCIM, the Ocean Circulation Inverse Model [10, 11]. OCIM is a non-seasonal global ocean circulation model based on a transport matrix, that enables us to assess how much carbon is stored in the oceans through the different pathways. Sequestration estimates are more representative than a global flux below an arbitrarily chosen depth [5]. First, a depth chosen arbitrarily has no biological meaning, unless it is chosen as the mixed layer depth below which carbon is effectively removed from phytoplankton [5]. Second, a high carbon flux does not necessarily mean that a high amount of carbon will be stored. For example, carbon sinking below the euphotic zone in upwelling areas may return to the surface quickly. Consequently, computing how much carbon the global ocean sequesters is a more robust way of assessing the efficiency of the biological carbon pump.

The concentration of biologically sequestered DIC is computed by finding the equilibrium $C_{in}$ of the following equation:

$$\frac{d\mathrm{C}_{in}}{dt} = \mathbf{A}\mathrm{C}_{in} + J_{res}, \tag{49}$$

where $\mathbf{A}$ is the advection-diffusion matrix transport operator from the OCIM, $J_{res}$ is the source of DIC, and $\mathrm{C}_{in}$ is dissolved inorganic carbon due to respiration. This equation is solved subject to a boundary condition of $C_{in} = 0$ at the sea surface, mimicking instantaneous air-sea $CO_2$ equilibration, and this calculation is repeated for each considered pathway.

After solving for $\mathrm{C}_{in}$ in equation (49), we volume integrate to obtain the total sequestered carbon inventory due to each mechanism (in Pg C). Finally, dividing the total sequestered carbon inventory for each pathway by the source term of carbon (in PgC $yr^{-1}$) yields a sequestration time (in yr), an indication of the efficiency of the considered pathway.

### 4 Difference between carbon export and injection

Export refers to carbon that is transported as organic carbon (here, below the euphotic zone). The carbon can be transported passively (i.e. sinking) or actively (i.e. by migrating organisms that then respire or egest it). Active export refers to the (organic) carbon that is transported by metazoans from above the euphotic zone to below. It is then defined as respiration of metazoans plus deadfalls and fecal pellet production, plus predation mortality minus ingestion (all of these processes considered below the euphotic zone only). This can be simplified by noting that active export is then active injection plus predation mortality (below the euphotic zone) minus ingestion (below the euphotic zone). Passive carbon export refers to organic carbon (fecal pellets or carcasses) passively sinking below the euphotic zone.

Injection refers to carbon that is transformed into DIC (below the euphotic zone), and that is then sequestered in the ocean's interior. In our model, carbon injection and carbon export differ, because organisms can consume detritus and bring it back to the euphotic zone while migrating. Additionally, the organism exporting carbon below the euphotic zone (the migrant) may not be the one actually injecting it as DIC, for example if it is preyed upon. Carbon injection is a more straightforward way to consider carbon transport out of the system, as it considers carbon that is turned into DIC and so rendered inaccessible to metazoans. In addition, it is what makes more sense when computing carbon sequestration and sequestration time scale as it is the carbon that is directly injected in the ocean at a specific depth.

Passive injection is passive export minus the fraction of particulate organic carbon (POC) sinking from above the euphotic zone that is consumed by metazoans below the euphotic zone. Active (or direct) injection refers to the carbon turned into DIC by metazoan processes happening below the euphotic zone (i.e. respiration, and bacterial degradation of fecal pellets and deadfalls produced below the euphotic zone).

This distinction between export and injection is hard to make in observational studies that generally ignore these complexities since they don't have enough information to parse out all the terms in the food-web. Our model allows to compare the two (table S1). The difference between injection and export, 1.1 (0.3-1.7) PgC/yr in total comes from the fact that there can be production below the euphotic zone in our model, as the distribution of resources is not directly linked to the limit of the euphotic zone (eq. 47). This difference can be attributed to mixing export, or, also, to diel migrations of microzooplankton not modelled here. At the global scale, global export and global injection are very similar. All carbon injected below the euphotic zone is exported, and carbon exported below the euphotic zone ends up being injected, unless it is brought back to the surface and respired in the euphotic zone – something that can happen but only represents a small flux compared to total export and injection.

Table S1: Comparison between global export and injection. All values are reported in PgC / yr.

| Functional group | Export | | | Injection | | |
|---|---|---|---|---|---|---|
| | Active | Passive | Total | Active | Passive | Total |
| Meso zooplankton | 0.3 (0.1 - 0.4) | 0.6 (0.2 - 0.9) | 0.9 (0.3 - 1.3) | 0.2 (0.06 - 0.3) | 0.5 (0.2 - 0.7) | 0.6 (0.3 - 1.0) |
| Macro zooplankton | 0.4 (0.2 - 0.6) | 0.3 (0.2 - 0.4) | 0.7 (0.4 - 1.0) | 0.9 (0.4 - 1.4) | 0.2 (0.1 - 0.3) | 1.1 (0.4 - 1.7) |
| Mesopelagic fish | 0.05 (0.01 - 0.3) | 0.2 (0.05 - 0.5) | 0.3 (0.06 - 0.8) | 0.8 (0.2 - 1.6) | 0.2 (0.04 - 0.4) | 1.0 (0.2 - 2.0) |
| Forage fish | -0.01 (-0.02 - -0.005) | 0.1 (0.05 - 0.2) | 0.1 (0.03 - 0.2) | 0.06 (0.03 - 0.1) | 0.09 (0.04 - 0.1) | 0.2 (0.1 - 0.2) |
| Large pelagic | 0.03 (7e-03 - 0.05) | 9e-04 (3e-04 - 2e-03) | 0.04 (7e-03 - 0.05) | 0.04 (0.01 - 0.1) | 8e-04 (3e-04 - 2e-03) | 0.04 (0.01 - 0.1) |
| Jellyfish | 0.02 (6e-03 - 0.05) | 0.04 (0.01 - 0.07) | 0.06 (0.02 - 0.1) | 0.07 (0.02 - 0.1) | 0.04 (0.01 - 0.07) | 0.1 (0.04 - 0.2) |
| Total | 0.8 (0.4 - 1.1) | 1.2 (0.8 - 1.9) | 2.0 (1.2 - 3.0) | 2.0 (0.9 - 3.2) | 1.0 (0.6 - 1.5) | 3.1 (1.5 - 4.7) |

### 5 Additional results

#### 5.1 DSL depth and computed POC flux comparison with data

Archived 38 kHz echosounder data were collated and weighted mean depth (WMD) of water column back-scattering intensity was extracted following the approach of [26]. See figure S7 for an example of raw echosounder data processing. The majority of collated raw data that were collected in the Indian and Pacific oceans were downloaded from the Integrated Marine Observing System [21], and most of the data collated in the Atlantic Ocean were obtained through the British Oceanographic Data Center (BODC, www.bodc.ac.uk) and the British Antarctic Survey [37]. Only daytime on-transect calibrated data were considered. WMD was calculated between 20 and 1000 m depth for 10 km along track segments and results were visually checked against plotted echograms. WMD values were excluded if echograms were deemed too noisy (e.g. when false bottoms were present). In most cases, WMD aligned with the depth of the strongest deep scattering layer. However, in some instances, particularly when epipelagic (0-200m) backscattering intensity was relatively high (or mesopelagic backscattering intensity relatively low), WMD was shallower than the DSL depth.

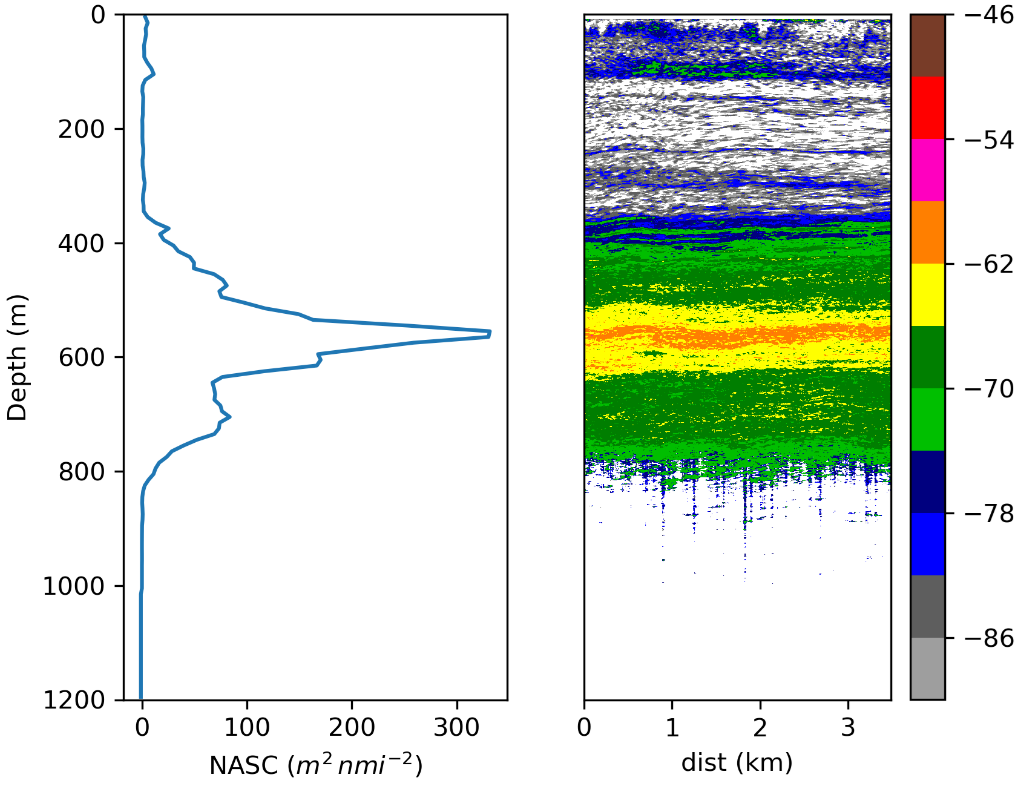

Figure S7: Example of raw echosounder data processing. Left plot: Nautical-area scattering coefficient (NASC, $m^2$ $nmi^{-2}$, average echo intensity per square nautical mile over a given depth range) depth profile (0 to 1200m by 10m intervals). Right plot: Echogram showing volume backscattering strength (Sv, dB re $m^{-1}$, average echo intensity per $m^3$) by depth and along-track distance (dist, km). Data collected from the R/V *Hesperides* during the *Malaspina* 2010 Spanish Circumnavigation Expedition using a EK60 echosounder operating at 38 kHz. In this example, daytime observations made on 8/4/2011 (Latitude = -33.9, Longitude = 156.8, duration = 1 hour) are shown. Calculated NASC-weighted mean depth (WMD) was 572 m.

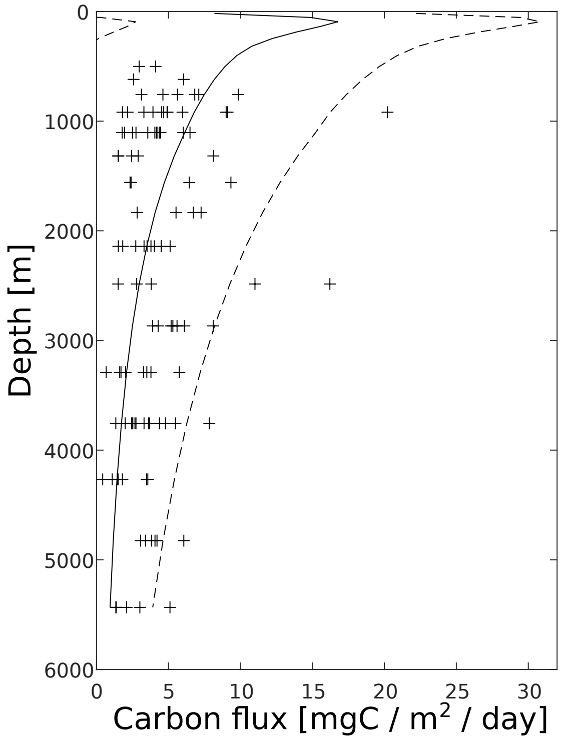

Figure S8: Globally averaged computed carbon flux at any depth (line), and observed carbon flux (crosses) from sediment traps data [29]. Dashed lines represent the minimum and maximum globally averaged carbon flux across sensitivity scenarios.

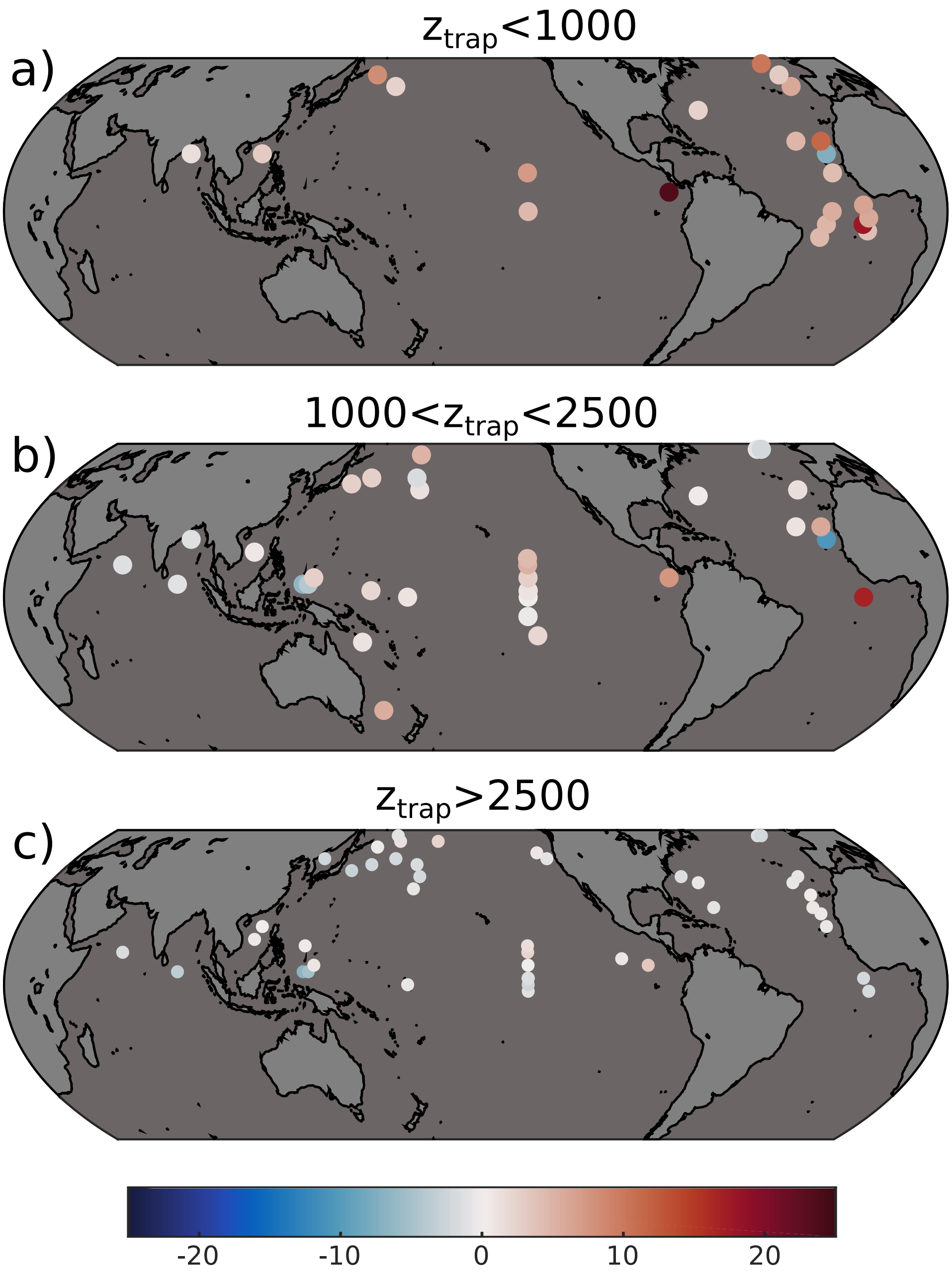

Figure S9: Difference between observed (from sediment traps data, [29]) and modeled POC flux. (a) Sediment traps between 0 and 1000 m, (b) sediment traps between 1000 and 2500 m, (c) sediment traps deeper than 2500 m.

#### 5.2 Regional sequestration potential

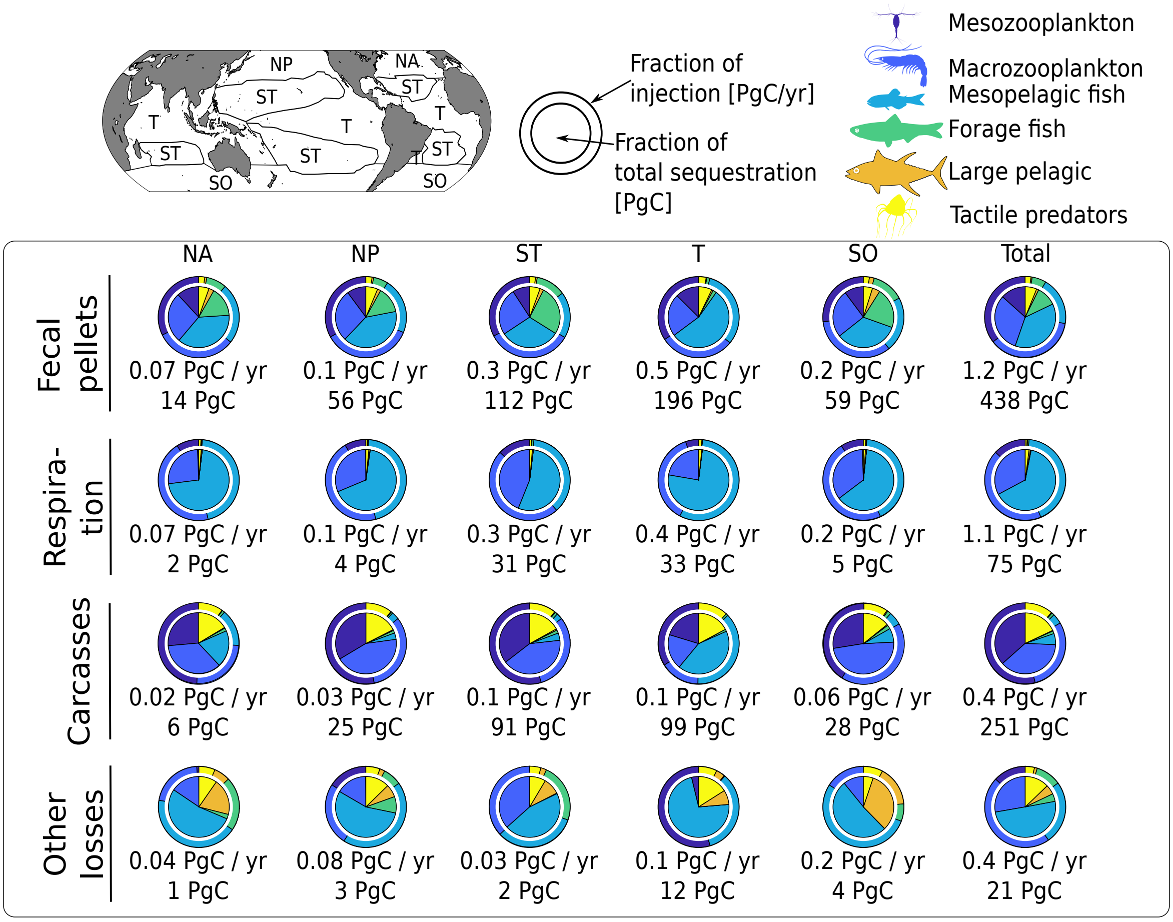

Figure S10: Geographical and functional breakout of carbon exported and sequestrated. NA stands for North Atlantic, NP for North Pacific, ST for subtropical gyres, T for tropics and upwelling zones, and SO for Southern Oceans. The figures below each pie chart represent the global contribution of the geographical zone in focus to carbon export and sequestration.

#### 5.3 Respiration and excretion below the euphotic zone

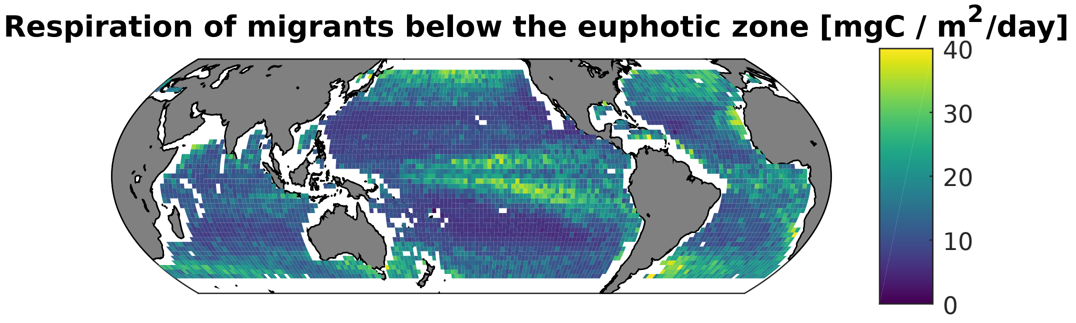

Figure S11: Simulated total respiration of organisms below the euphotic zone (i.e. injection via respiration).

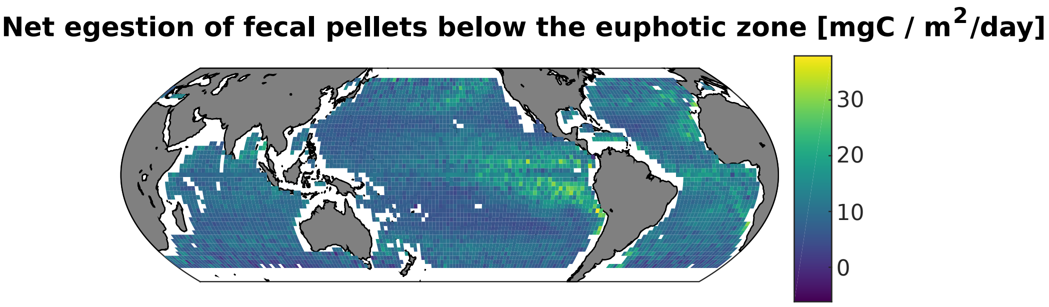

Figure S12: Simulated net egestion (= fecal pellet excretion - consumption) of carbon below the euphotic zone (i.e. injection via fecal pellets egestion directly below the euphotic zone).

#### 5.4 Comparison with apparent oxygen utilization

The apparent oxygen utilization (AOU) is the difference between the oxygen solubility and the measured oxygen concentration in water [15]. As such, it can be seen as a proxy of the total respiration (animal and bacterial) that happened in a water mass, and can be related to the sequestration of DIC in the water mass considered (assuming a molar ratio of 1.4 mol $O_2$ per mol C during respiration). The global carbon sequestration derived from the World Ocean Atlas' AOU (WOA AOU) data [15] is 1765 PgC, but a recent study estimated that the interior oceans stores 1300 ($\pm$ 230) PgC [7]. When investigating ocean transects (figure S13), we see that we do not predict more oxygen utilization than there actually is. This is consistent with the fact that we do not model the entire food-web and do not account for all ocean processes. The qualitative pattern differences in the Pacific and in the Indian Ocean may be explained by the fact that there is no source term above (and below) 45°N (45°S) in our model due to its spatial coverage. Moreover, our simulated AOU has a deeper maximum than the observed AOU, consistent with the fact that we are resolving the processes with faster sinking speed, whereas remaining processes (e.g. remineralization of DOC, aggregates and small fecal pellets from micro-zooplankton) would be concentrated in the upper oceans.

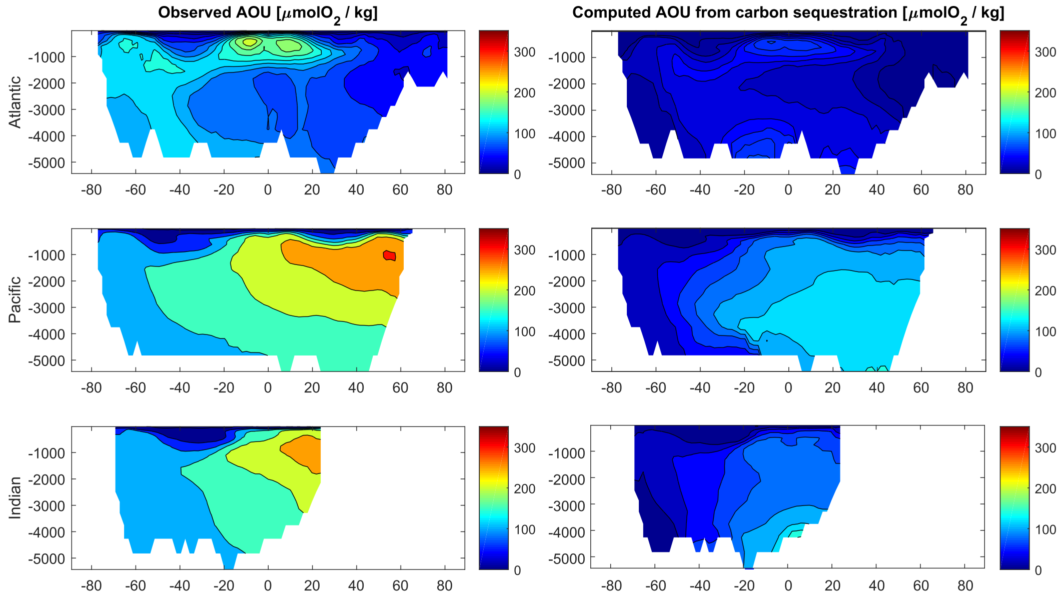

Figure S13: Transects of observed AOU (WOA, left column) and simulated AOU corresponding to the carbon sequestration computed by our model (right column) in the Atlantic, Pacific and Indian oceans.

### 6 Sensitivity analysis

#### 6.1 Global model

Due to the very high number of parameters and high computational cost of each simulation, a complete sensitivity analysis on the parameter space could not be completed. Instead, we performed an analysis of the most sensitive parameters with regards to carbon export, i.e. fecal pellets sinking rates, bacterial degradation rate, biomass of all functional groups, biomass of mesopelagic fish only, assimilation efficiencies, assimilation efficiency for detritus only, swimming speeds of all organisms, swimming speeds of mesopelagic fish only, and reference and maximum temperatures for all temperature-dependent rates.

We give the detailed results of the sensitivity analysis below. In summary, we find that our results are quite robust. Despite small relative variations, the global trends in DSL depths are consistent between the different simulations (figure S14).

The results are the most sensitive (in terms of carbon export and sequestration) to variations in biomass, assimilation efficiencies and fecal pellets sinking rates (as well as organisms swimming speed for carbon export). Details of the variations for the different parameters are given below.

In addition, other organisms and behaviours not included in our model would modify trophic coupling and the estimated strength of the biological carbon pump, particularly at depth. Our study is focused on a global view which necessarily requires that certain details are omitted, and consequently the sensitivity of our model to these processes could not be quantified. For example, mesopelagic fish in our model have a low trophic transfer efficiency (∼1 %), potentially because some of their predators (piscivorous mesopelagic fish, bathypelagic fish, deep-sea squids) are not represented in this model. However, higher order predators repackage carbon into larger faster-sinking pellets, so not including these organisms means that we are providing a conservative estimate of carbon sequestration for mesopelagic fish and their predators. In addition, because of its daily temporal resolution, our model predicts that the entire mesopelagic community migrates to the surface at night. While this is counter to what happens in nature, as acoustic backscatter at depth is often detected at night by echosounders [26], the exact proportion of migrating organisms is unknown. The inclination of individual organisms to migrate to the surface may be species-specific, but also depends on the current satiation level of organisms [4].

Table S2: Injection via basal respiration, net excretion, carcasse production, and other losses in the different scenarios, in PgC / yr.

| Parameter | % variation | Respiration | Fecal pellets | Carcasses | Other losses |
|---|---|---|---|---|---|
| Reference | - | 1.1 | 1.2 | 0.4 | 0.4 |
| Total biomass | 50 % | 0.6 | 0.5 | 0.3 | 0.1 |
| | 150 % | 1.7 | 1.8 | 0.5 | 0.6 |
| Mesopelagic fish | 20 % | 0.7 | 1.0 | 0.3 | 0.8 |
| biomass | 50 % | 0.9 | 1.0 | 0.3 | 0.7 |
| | 150 % | 1.4 | 1.3 | 0.4 | 0.4 |
| | 200 % | 1.7 | 1.4 | 0.4 | 0.5 |
| Sinking rate | 50 % | 1.1 | 1.0 | 0.3 | 0.4 |
| | 150 % | 1.2 | 1.3 | 0.4 | 0.4 |
| Bacterial degradation | 50 % | 1.1 | 1.2 | 0.4 | 0.4 |
| rate | 150 % | 1.2 | 1.1 | 0.4 | 0.4 |
| Assimilation efficiency | 90 % | 1.1 | 1.7 | 0.4 | 0.3 |
| | 110 % | 1.2 | 0.6 | 0.4 | 0.6 |
| Detritus assimilation | 80 % | 1.2 | 1.2 | 0.4 | 0.5 |
| efficiency | 120 % | 1.1 | 1.1 | 0.4 | 0.4 |
| Swimming speeds | 50 % | 0.7 | 1.1 | 0.3 | 0.6 |
| | 150% | 1.6 | 1.2 | 0.8 | 0.2 |
| Mesopelagic swimming | 50 % | 1.1 | 1.1 | 0.6 | 0.3 |
| speed | 150 % | 1.2 | 1.2 | 0.3 | 0.4 |
| Reference and maximum | 90 % | 1.2 | 1.2 | 0.5 | 0.3 |
| temperatures | 110 % | 1.1 | 1.2 | 0.4 | 0.4 |

Contrarily to other parameters that were varied of ± 50 %, the mesopelagic fish biomass varied between 20 and 200 % of the reference value, following its current uncertainty estimates [39].

In addition, assimilation efficiencies levels cannot be set to 150 % of the reference levels as they

Table S3: Total export, corresponding sequestration and sequestration time for the different pathways considered in the model. Model run with biomasses equal to 50 % the reference value.

| Organism | Respiration pathway | | | Fecal pellets pathway | | | Carcasses pathway | | | Other losses | | | Total | | |
|---|---|---|---|---|---|---|---|---|---|---|---|---|---|---|---|
| | Injection [PgC $yr^{-1}$] | Sequestr. [PgC] | Sequestr. time [yr] | Injection [PgC $yr^{-1}$] | Sequestr. [PgC] | Sequestr. time [yr] | Injection [PgC $yr^{-1}$] | Sequestr. [PgC] | Sequestr. time [yr] | Injection [PgC $yr^{-1}$] | Sequestr. [PgC] | Sequestr. time [yr] | Injection [PgC $yr^{-1}$] | Sequestr. [PgC] | Sequestr. time [yr] |
| Meso zoopl. | 0.1 | 0.4 | 6 | 0.2 | 24 | 152 | 0.1 | 33 | 414 | -2e-02 | -0.14 | 6 | 0.3 | 58 | 210 |
| Macro zoopl. | 0.2 | 10 | 39 | 0.2 | 59 | 298 | 0.1 | 43 | 788 | 0.1 | 2.0 | 31 | 0.6 | 114 | 201 |
| Meso-pelagic | 0.2 | 25 | 104 | 0.1 | 78 | 601 | 1e-01 | 124 | 945 | 0.04 | 3 | 62 | 0.6 | 231 | 416 |
| Forage fish | 1e-03 | 0.01 | 7 | 0.05 | 33 | 722 | 2e-03 | 2 | 968 | 0.03 | 0.1 | 5 | 0.1 | 35 | 457 |
| Large pelagic | 1e-03 | 0.2 | 156 | 4e-03 | 4 | 999 | 5e-04 | 0.4 | 1003 | 0.01 | 1 | 152 | 0.01 | 5 | 426 |
| Jellyfish | 5e-03 | 0.5 | 95 | 0.01 | 7 | 833 | 0.02 | 21 | 1015 | 2e-03 | 0.1 | 50 | 0.04 | 28 | 781 |
| **Total** | **0.6** | **36** | **65** | **0.5** | **205** | **376** | **0.3** | **223** | **774** | **0.1** | **6** | **49** | **1.5** | **471** | **310** |

Table S4: Total export, corresponding sequestration and sequestration time for the different pathways considered in the model. Model run with biomasses equal to 150% the reference value.

| Organism | Respiration pathway | | | Fecal pellets pathway | | | Carcasses pathway | | | Other losses | | | Total | | |
|---|---|---|---|---|---|---|---|---|---|---|---|---|---|---|---|
| | Injection [PgC $yr^{-1}$] | Sequestr. [PgC] | Sequestr. time [yr] | Injection [PgC $yr^{-1}$] | Sequestr. [PgC] | Sequestr. time [yr] | Injection [PgC $yr^{-1}$] | Sequestr. [PgC] | Sequestr. time [yr] | Injection [PgC $yr^{-1}$] | Sequestr. [PgC] | Sequestr. time [yr] | Injection [PgC $yr^{-1}$] | Sequestr. [PgC] | Sequestr. time [yr] |
| Meso zoopl. | 0.2 | 0.8 | 5 | 0.6 | 76 | 133 | 0.2 | 99 | 413 | -1e-02 | -0.05 | 4 | 1.0 | 176 | 185 |
| Macro zoopl. | 0.8 | 30 | 39 | 0.6 | 175 | 298 | 0.2 | 123 | 786 | 0.1 | 3.5 | 35 | 1.6 | 332 | 206 |
| Meso-pelagic | 0.8 | 82 | 103 | 0.4 | 246 | 599 | 5e-02 | 39 | 841 | 0.3 | 20 | 65 | 1.6 | 386 | 248 |
| Forage fish | 6e-03 | 0.09 | 16 | 0.1 | 105 | 722 | 6e-03 | 6 | 969 | 0.1 | 1.1 | 12 | 0.2 | 113 | 454 |
| Large pelagic | 4e-03 | 0.6 | 156 | 0.03 | 25 | 985 | 1e-03 | 1 | 1003 | 0.06 | 9 | 153 | 0.09 | 37 | 399 |
| Jellyfish | 2e-02 | 2 | 127 | 0.06 | 53 | 829 | 0.06 | 62 | 1019 | 0.07 | 6 | 76 | 0.2 | 122 | 569 |
| **Total** | **1.7** | **115** | **66** | **1.8** | **680** | **377** | **0.5** | **330** | **647** | **0.6** | **39** | **64** | **4.7** | **1165** | **249** |

Table S5: Total export, corresponding sequestration and sequestration time for the different pathways considered in the model. Model run with mesopelagic biomass equal to 20% the reference value.

| Organism | Respiration pathway | | | Fecal pellets pathway | | | Carcasses pathway | | | Other losses | | | Total | | |
|---|---|---|---|---|---|---|---|---|---|---|---|---|---|---|---|
| | Injection [PgC $yr^{-1}$] | Sequestr. [PgC] | Sequestr. time [yr] | Injection [PgC $yr^{-1}$] | Sequestr. [PgC] | Sequestr. time [yr] | Injection [PgC $yr^{-1}$] | Sequestr. [PgC] | Sequestr. time [yr] | Injection [PgC $yr^{-1}$] | Sequestr. [PgC] | Sequestr. time [yr] | Injection [PgC $yr^{-1}$] | Sequestr. [PgC] | Sequestr. time [yr] |
| Meso zoopl. | 0.1 | 0.5 | 5 | 0.4 | 51 | 140 | 0.2 | 66 | 414 | 4e-02 | 0.19 | 5 | 0.7 | 118 | 175 |
| Macro zoopl. | 0.5 | 20 | 39 | 0.4 | 116 | 293 | 0.1 | 83 | 784 | 0.7 | 22.3 | 34 | 1.7 | 241 | 145 |
| Meso-pelagic | 0.1 | 11 | 103 | 0.1 | 34 | 596 | 8e-03 | 7 | 871 | 0.04 | 3 | 62 | 0.2 | 53 | 255 |
| Forage fish | 3e-03 | 0.02 | 6 | 0.1 | 72 | 722 | 4e-03 | 4 | 968 | 0.1 | 0.2 | 4 | 0.2 | 76 | 461 |
| Large pelagic | 3e-03 | 0.4 | 156 | 0.01 | 11 | 975 | 9e-04 | 1 | 1003 | 0.02 | 4 | 154 | 0.04 | 16 | 405 |
| Jellyfish | 1e-02 | 1 | 113 | 0.03 | 24 | 831 | 0.04 | 41 | 1017 | 0.03 | 2 | 65 | 0.1 | 68 | 641 |
| **Total** | **0.7** | **32** | **44** | **1.0** | **307** | **321** | **0.3** | **202** | **634** | **0.8** | **31** | **37** | **2.9** | **573** | **201** |

Table S6: Total export, corresponding sequestration and sequestration time for the different pathways considered in the model. Model run with mesopelagic biomass equal to 50% the reference value.

| Organism | Respiration pathway | | | Fecal pellets pathway | | | Carcasses pathway | | | Other losses | | | Total | | |
|---|---|---|---|---|---|---|---|---|---|---|---|---|---|---|---|
| | Injection [PgC $yr^{-1}$] | Sequestr. [PgC] | Sequestr. time [yr] | Injection [PgC $yr^{-1}$] | Sequestr. [PgC] | Sequestr. time [yr] | Injection [PgC $yr^{-1}$] | Sequestr. [PgC] | Sequestr. time [yr] | Injection [PgC $yr^{-1}$] | Sequestr. [PgC] | Sequestr. time [yr] | Injection [PgC $yr^{-1}$] | Sequestr. [PgC] | Sequestr. time [yr] |
| Meso zoopl. | 0.1 | 0.5 | 5 | 0.4 | 51 | 140 | 0.2 | 66 | 414 | 2e-02 | 0.08 | 4 | 0.6 | 118 | 181 |
| Macro zoopl. | 0.5 | 20 | 39 | 0.4 | 116 | 295 | 0.1 | 87 | 788 | 0.4 | 14.7 | 34 | 1.4 | 237 | 165 |
| Meso-pelagic | 0.3 | 27 | 102 | 0.1 | 83 | 597 | 3e-02 | 23 | 865 | 0.1 | 6 | 62 | 0.5 | 139 | 265 |
| Forage fish | 3e-03 | 0.02 | 7 | 0.1 | 70 | 721 | 4e-03 | 4 | 968 | 0.1 | 0.2 | 4 | 0.2 | 75 | 474 |
| Large pelagic | 3e-03 | 0.4 | 156 | 0.01 | 11 | 981 | 9e-04 | 1 | 1003 | 0.03 | 4 | 154 | 0.04 | 17 | 406 |
| Jellyfish | 1e-02 | 1 | 111 | 0.03 | 24 | 828 | 0.04 | 41 | 1018 | 0.03 | 2 | 64 | 0.1 | 69 | 634 |
| **Total** | **0.9** | **48** | **54** | **1.0** | **356** | **344** | **0.3** | **222** | **650** | **0.7** | **27** | **41** | **2.9** | **654** | **224** |

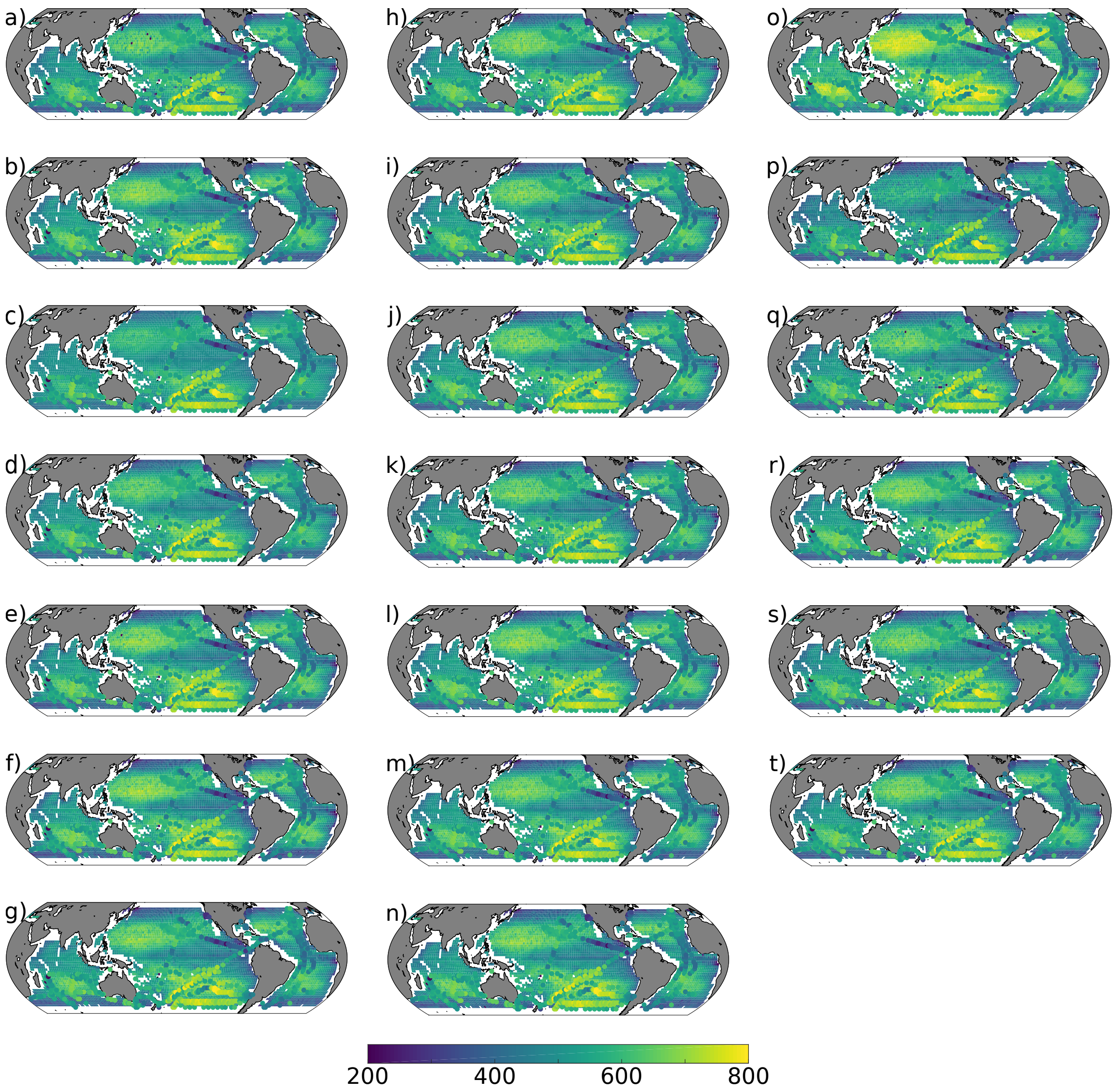

Figure S14: Predicted mean depth during daytime, weighted by biomass for the different sensitivity scenarios. Circles overlaid are the observed weighted mean depth recorded at 38 kHz [26]. (a) 50% of the biomass reference value, (b) 150% of biomass, (c) 20% of mesopelagic biomass, (d) 50% of mesopelagic biomass, (e) 150% of mesopelagic biomass, (f) 200% of mesopelagic biomass, (g) 50% of sinking rates, (h) 150% of sinking rates, (i) 50% of bacterial degradation rates, (j) 150% of bacterial degradation rate, (k) 90% of assimilation efficiencies, (l) 110% of assimilation efficiencies, (m) 80% of detritus assimilation efficiency, (n) 120% of detritus assimilation efficiency, (o) 50% of swimming speeds, (p) 150% of swimming speeds, (q) 50% of mesopelagic swimming speed, (r) 150% of mesopelagic swimming speed, (s) 90% of reference and maximum temperatures, (t) 110% of reference and maximum temperatures.

cannot be greater than 1. Moreover, variations of 50 % or more for the assimilation efficiencies are highly unrealistic. As such, when conducting this sensitivity analysis, reference assimilation efficiencies were varied of 10 %.

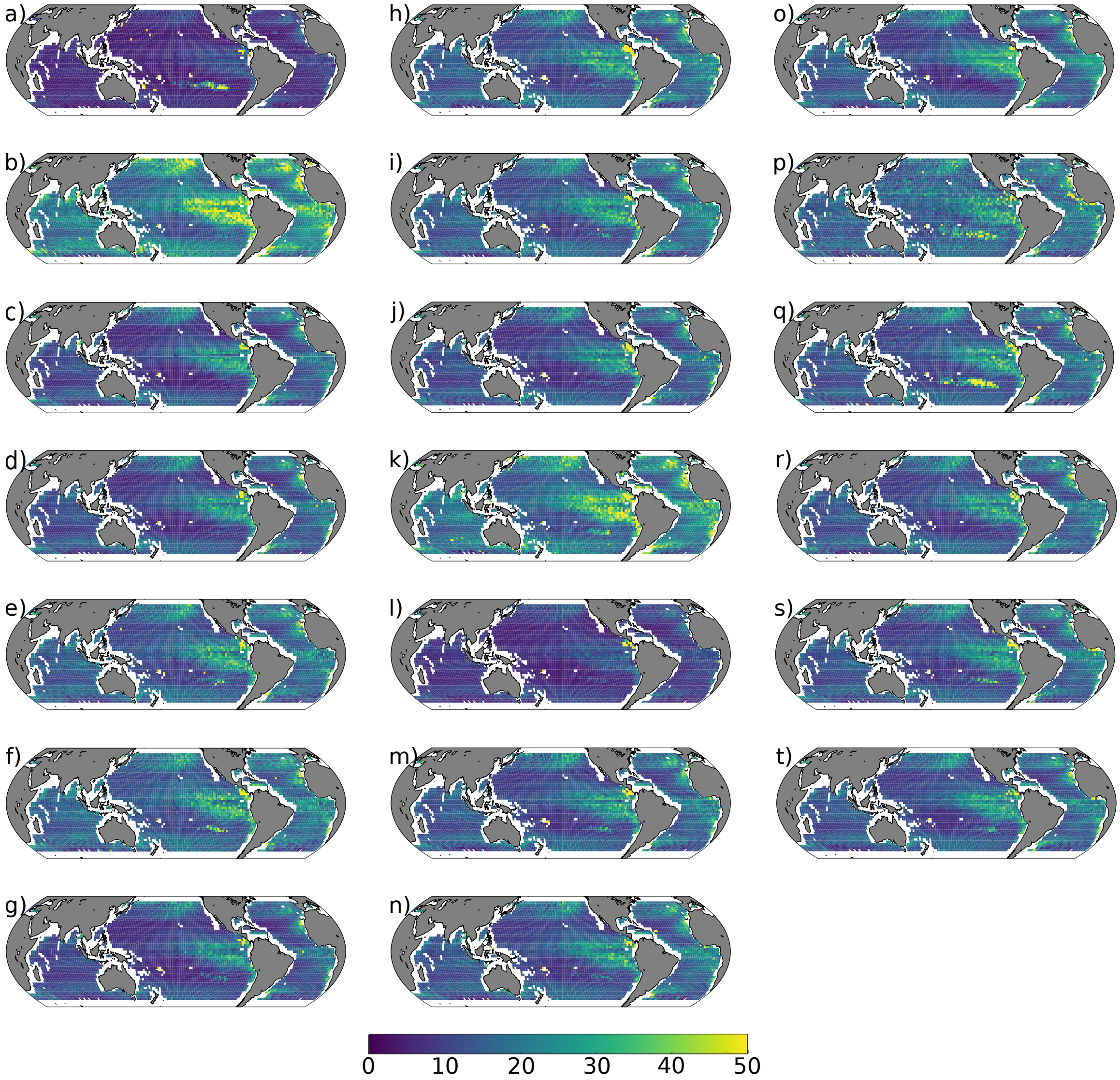

Figure S15: Predicted passive sinking flux at the base of the euphotic zone for the different scenarios, in mgC/m$^2$/day. (a) 50% of the biomass reference value, (b) 150% of biomass, (c) 20% of mesopelagic biomass, (d) 50% of mesopelagic biomass, (e) 150% of mesopelagic biomass, (f) 200% of mesopelagic biomass, (g) 50% of sinking rates, (h) 150% of sinking rates, (i) 50% of bacterial degradation rates, (j) 150% of bacterial degradation rate, (k) 90% of assimilation efficiencies, (l) 110% of assimilation efficiencies, (m) 80% of detritus assimilation efficiency, (n) 120% of detritus assimilation efficiency, (o) 50% of swimming speeds, (p) 150% of swimming speeds, (q) 50% of mesopelagic swimming speed, (r) 150% of mesopelagic swimming speed, (s) 90% of reference and maximum temperatures, (t) 110% of reference and maximum temperatures.

Table S7: Total export, corresponding sequestration and sequestration time for the different pathways considered in the model. Model run with mesopelagic biomass equal to 150 % the reference value.

| Organism | Respiration pathway | | | Fecal pellets pathway | | | Carcasses pathway | | | Other losses | | | Total | | |
|---|---|---|---|---|---|---|---|---|---|---|---|---|---|---|---|
| | Injection [PgC $yr^{-1}$] | Sequestr. [PgC] | Sequestr. time [yr] | Injection [PgC $yr^{-1}$] | Sequestr. [PgC] | Sequestr. time [yr] | Injection [PgC $yr^{-1}$] | Sequestr. [PgC] | Sequestr. time [yr] | Injection [PgC $yr^{-1}$] | Sequestr. [PgC] | Sequestr. time [yr] | Injection [PgC $yr^{-1}$] | Sequestr. [PgC] | Sequestr. time [yr] |
| Meso zoopl. | 0.1 | 0.5 | 5 | 0.4 | 51 | 141 | 0.2 | 66 | 414 | -2e-02 | -0.09 | 4 | 0.6 | 118 | 194 |
| Macro zoopl. | 0.5 | 20 | 39 | 0.4 | 118 | 299 | 0.1 | 87 | 788 | -0.3 | -8.5 | 32 | 0.7 | 216 | 289 |
| Meso-pelagic | 0.8 | 80 | 103 | 0.4 | 244 | 599 | 1e-01 | 101 | 899 | 0.3 | 17 | 63 | 1.6 | 443 | 282 |
| Forage fish | 3e-03 | 0.05 | 15 | 0.1 | 69 | 729 | 4e-03 | 4 | 969 | 0.1 | 0.6 | 11 | 0.2 | 74 | 457 |
| Large pelagic | 3e-03 | 0.4 | 156 | 0.01 | 13 | 996 | 9e-04 | 1 | 1003 | 0.03 | 4 | 154 | 0.04 | 18 | 410 |
| Jellyfish | 1e-02 | 1 | 114 | 0.03 | 25 | 829 | 0.04 | 41 | 1018 | 0.03 | 2 | 67 | 0.1 | 69 | 636 |
| **Total** | **1.4** | **102** | **72** | **1.3** | **520** | **399** | **0.4** | **300** | **702** | **0.4** | **24** | **60** | **3.5** | **946** | **270** |

Table S8: Total export, corresponding sequestration and sequestration time for the different pathways considered in the model. Model run with mesopelagic biomass equal to 200 % the reference value.

| Organism | Respiration pathway | | | Fecal pellets pathway | | | Carcasses pathway | | | Other losses | | | Total | | |
|---|---|---|---|---|---|---|---|---|---|---|---|---|---|---|---|
| | Injection [PgC $yr^{-1}$] | Sequestr. [PgC] | Sequestr. time [yr] | Injection [PgC $yr^{-1}$] | Sequestr. [PgC] | Sequestr. time [yr] | Injection [PgC $yr^{-1}$] | Sequestr. [PgC] | Sequestr. time [yr] | Injection [PgC $yr^{-1}$] | Sequestr. [PgC] | Sequestr. time [yr] | Injection [PgC $yr^{-1}$] | Sequestr. [PgC] | Sequestr. time [yr] |
| Meso zoopl. | 0.1 | 0.5 | 5 | 0.4 | 52 | 141 | 0.2 | 66 | 413 | -4e-02 | -0.17 | 4 | 0.6 | 118 | 201 |
| Macro zoopl. | 0.5 | 20 | 39 | 0.4 | 119 | 301 | 0.1 | 90 | 786 | -0.6 | -18.5 | 32 | 0.4 | 210 | 474 |
| Meso-pelagic | 1.0 | 107 | 103 | 0.5 | 317 | 602 | 1e-01 | 93 | 863 | 0.3 | 20 | 64 | 2.0 | 537 | 271 |
| Forage fish | 4e-03 | 0.08 | 19 | 0.1 | 66 | 735 | 4e-03 | 4 | 970 | 0.1 | 0.9 | 16 | 0.2 | 71 | 457 |
| Large pelagic | 3e-03 | 0.4 | 156 | 0.01 | 13 | 999 | 9e-04 | 1 | 1003 | 0.03 | 4 | 153 | 0.05 | 19 | 410 |
| Jellyfish | 1e-02 | 1 | 116 | 0.03 | 25 | 831 | 0.04 | 41 | 1018 | 0.03 | 2 | 69 | 0.1 | 69 | 642 |
| **Total** | **1.7** | **129** | **77** | **1.4** | **592** | **416** | **0.4** | **295** | **691** | **0.5** | **27** | **54** | **4.0** | **1043** | **261** |

Table S9: Total export, corresponding sequestration and sequestration time for the different pathways considered in the model. Model run with fecal pellets sinking rates equal to 50 % the reference value.

| Organism | Respiration pathway | | | Fecal pellets pathway | | | Carcasses pathway | | | Other losses | | | Total | | |
|---|---|---|---|---|---|---|---|---|---|---|---|---|---|---|---|
| | Injection [PgC $yr^{-1}$] | Sequestr. [PgC] | Sequestr. time [yr] | Injection [PgC $yr^{-1}$] | Sequestr. [PgC] | Sequestr. time [yr] | Injection [PgC $yr^{-1}$] | Sequestr. [PgC] | Sequestr. time [yr] | Injection [PgC $yr^{-1}$] | Sequestr. [PgC] | Sequestr. time [yr] | Injection [PgC $yr^{-1}$] | Sequestr. [PgC] | Sequestr. time [yr] |
| Meso zoopl. | 0.1 | 0.5 | 5 | 0.3 | 12 | 43 | 0.1 | 25 | 177 | 2e-02 | 0.06 | 4 | 0.5 | 37 | 70 |
| Macro zoopl. | 0.5 | 19 | 37 | 0.4 | 41 | 115 | 0.1 | 53 | 488 | 0.05 | 1.5 | 32 | 1.0 | 114 | 112 |
| Meso-pelagic | 0.5 | 53 | 103 | 0.2 | 66 | 271 | 5e-02 | 31 | 622 | 0.2 | 12 | 63 | 1.0 | 161 | 162 |
| Forage fish | 3e-03 | 0.04 | 12 | 0.1 | 34 | 415 | 4e-03 | 3 | 757 | 0.1 | 0.5 | 10 | 0.1 | 38 | 262 |
| Large pelagic | 3e-03 | 0.4 | 156 | 0.01 | 11 | 878 | 9e-04 | 1 | 956 | 0.03 | 4 | 154 | 0.04 | 16 | 374 |
| Jellyfish | 1e-02 | 1 | 117 | 0.03 | 13 | 501 | 0.04 | 34 | 854 | 0.03 | 2 | 71 | 0.1 | 50 | 492 |
| **Total** | **1.1** | **74** | **65** | **1.0** | **177** | **178** | **0.3** | **147** | **428** | **0.4** | **20** | **56** | **2.8** | **417** | **147** |

Table S10: Total export, corresponding sequestration and sequestration time for the different pathways considered in the model. Model run with fecal pellets sinking rates equal to 150 % the reference value.

| Organism | Respiration pathway | | | Fecal pellets pathway | | | Carcasses pathway | | | Other losses | | | Total | | |
|---|---|---|---|---|---|---|---|---|---|---|---|---|---|---|---|
| | Injection [PgC $yr^{-1}$] | Sequestr. [PgC] | Sequestr. time [yr] | Injection [PgC $yr^{-1}$] | Sequestr. [PgC] | Sequestr. time [yr] | Injection [PgC $yr^{-1}$] | Sequestr. [PgC] | Sequestr. time [yr] | Injection [PgC $yr^{-1}$] | Sequestr. [PgC] | Sequestr. time [yr] | Injection [PgC $yr^{-1}$] | Sequestr. [PgC] | Sequestr. time [yr] |
| Meso zoopl. | 0.1 | 0.7 | 6 | 0.4 | 111 | 262 | 0.2 | 103 | 613 | -2e-02 | -0.08 | 5 | 0.7 | 214 | 310 |
| Macro zoopl. | 0.5 | 20 | 40 | 0.4 | 196 | 472 | 0.1 | 105 | 908 | 0.1 | 3.6 | 34 | 1.1 | 324 | 284 |
| Meso-pelagic | 0.5 | 54 | 103 | 0.3 | 229 | 792 | 6e-02 | 59 | 927 | 0.2 | 12 | 63 | 1.1 | 354 | 333 |
| Forage fish | 3e-03 | 0.03 | 11 | 0.1 | 88 | 856 | 4e-03 | 4 | 1028 | 0.1 | 0.6 | 9 | 0.2 | 93 | 541 |
| Large pelagic | 3e-03 | 0.4 | 156 | 0.01 | 12 | 1015 | 9e-04 | 1 | 1016 | 0.03 | 4 | 154 | 0.04 | 18 | 416 |
| Jellyfish | 1e-02 | 1 | 110 | 0.03 | 29 | 951 | 0.04 | 44 | 1064 | 0.03 | 2 | 63 | 0.1 | 76 | 681 |
| **Total** | **1.2** | **76** | **65** | **1.3** | **666** | **523** | **0.4** | **316** | **803** | **0.4** | **22** | **56** | **3.2** | **1080** | **335** |

Table S11: Total export, corresponding sequestration and sequestration time for the different pathways considered in the model. Model run with bacterial degradation rate equal to 50 % the reference value.

| Organism | Respiration pathway | | | Fecal pellets pathway | | | Carcasses pathway | | | Other losses | | | Total | | |
|---|---|---|---|---|---|---|---|---|---|---|---|---|---|---|---|
| | Export [PgC $yr^{-1}$] | Sequestr. [PgC] | Sequestr. time [yr] | Export [PgC $yr^{-1}$] | Sequestr. [PgC] | Sequestr. time [yr] | Export [PgC $yr^{-1}$] | Sequestr. [PgC] | Sequestr. time [yr] | Export [PgC $yr^{-1}$] | Sequestr. [PgC] | Sequestr. time [yr] | Export [PgC $yr^{-1}$] | Sequestr. [PgC] | Sequestr. time [yr] |
| Meso zoopl. | 0.1 | 0.5 | 5 | 0.4 | 138 | 360 | 0.2 | 127 | 749 | -2e-04 | -6e-04 | 4 | 0.7 | 266 | 408 |
| Macro zoopl. | 0.5 | 20 | 39 | 0.4 | 254 | 617 | 0.1 | 103 | 972 | 0.1 | 2.3 | 34 | 1.1 | 379 | 347 |
| Meso-pelagic | 0.5 | 53 | 102 | 0.3 | 242 | 896 | 7e-02 | 71 | 984 | 0.2 | 10 | 63 | 1.0 | 376 | 366 |
| Forage fish | 3e-03 | 0.04 | 12 | 0.1 | 89 | 909 | 4e-03 | 5 | 1055 | 0.1 | 0.5 | 8 | 0.2 | 94 | 555 |
| Large pelagic | 3e-03 | 0.4 | 156 | 0.01 | 13 | 1031 | 9e-04 | 1 | 1021 | 0.03 | 4 | 154 | 0.04 | 19 | 420 |
| Jellyfish | 1e-02 | 1 | 117 | 0.03 | 30 | 1005 | 0.04 | 44 | 1084 | 0.02 | 2 | 70 | 0.1 | 77 | 727 |
| **Total** | **1.1** | **75** | **66** | **1.2** | **766** | **635** | **0.4** | **351** | **891** | **0.4** | **19** | **55** | **3.1** | **1211** | **392** |

Table S12: Total export, corresponding sequestration and sequestration time for the different pathways considered in the model. Model run with bacterial degradation rate equal to 150 % the reference value.

| Organism | Respiration pathway | | | Fecal pellets pathway | | | Carcasses pathway | | | Other losses | | | Total | | |
|---|---|---|---|---|---|---|---|---|---|---|---|---|---|---|---|
| | Injection [PgC $yr^{-1}$] | Sequestr. [PgC] | Sequestr. time [yr] | Injection [PgC $yr^{-1}$] | Sequestr. [PgC] | Sequestr. time [yr] | Injection [PgC $yr^{-1}$] | Sequestr. [PgC] | Sequestr. time [yr] | Injection [PgC $yr^{-1}$] | Sequestr. [PgC] | Sequestr. time [yr] | Injection [PgC $yr^{-1}$] | Sequestr. [PgC] | Sequestr. time [yr] |
| Meso zoopl. | 0.1 | 0.6 | 5 | 0.3 | 27 | 78 | 0.2 | 39 | 257 | -6e-03 | -0.03 | 5 | 0.6 | 66 | 110 |
| Macro zoopl. | 0.5 | 20 | 39 | 0.4 | 67 | 177 | 0.1 | 74 | 617 | 0.1 | 2.7 | 33 | 1.1 | 163 | 150 |
| Meso-pelagic | 0.5 | 54 | 103 | 0.3 | 108 | 396 | 5e-02 | 38 | 736 | 0.2 | 12 | 63 | 1.0 | 212 | 205 |
| Forage fish | 3e-03 | 0.03 | 10 | 0.1 | 53 | 551 | 4e-03 | 4 | 865 | 0.1 | 0.5 | 8 | 0.2 | 57 | 346 |
| Large pelagic | 3e-03 | 0.4 | 156 | 0.01 | 11 | 934 | 9e-04 | 1 | 981 | 0.03 | 4 | 154 | 0.04 | 17 | 392 |
| Jellyfish | 1e-02 | 1 | 110 | 0.03 | 19 | 654 | 0.04 | 38 | 939 | 0.03 | 2 | 64 | 0.1 | 60 | 552 |
| **Total** | **1.2** | **76** | **66** | **1.1** | **285** | **253** | **0.4** | **193** | **525** | **0.4** | **21** | **55** | **3.0** | **575** | **190** |

Table S13: Total export, corresponding sequestration and sequestration time for the different pathways considered in the model. Model run with assimilation efficiencies equal to 90 % the reference values.

| Organism | Respiration pathway | | | Fecal pellets pathway | | | Carcasses pathway | | | Other losses | | | Total | | |
|---|---|---|---|---|---|---|---|---|---|---|---|---|---|---|---|
| | Injection [PgC $yr^{-1}$] | Sequestr. [PgC] | Sequestr. time [yr] | Injection [PgC $yr^{-1}$] | Sequestr. [PgC] | Sequestr. time [yr] | Injection [PgC $yr^{-1}$] | Sequestr. [PgC] | Sequestr. time [yr] | Injection [PgC $yr^{-1}$] | Sequestr. [PgC] | Sequestr. time [yr] | Injection [PgC $yr^{-1}$] | Sequestr. [PgC] | Sequestr. time [yr] |
| Meso zoopl. | 0.1 | 0.4 | 4 | 0.5 | 73 | 141 | 0.2 | 65 | 413 | -1e-02 | -0.04 | 3 | 0.8 | 139 | 183 |
| Macro zoopl. | 0.5 | 19 | 38 | 0.6 | 183 | 303 | 0.1 | 89 | 776 | -0.1 | -2.2 | 32 | 1.2 | 289 | 251 |
| Meso-pelagic | 0.5 | 51 | 101 | 0.4 | 227 | 602 | 9e-02 | 81 | 919 | 0.1 | 6 | 62 | 1.1 | 365 | 342 |
| Forage fish | 3e-03 | 0.03 | 11 | 0.1 | 107 | 721 | 4e-03 | 4 | 968 | 0.1 | 0.4 | 8 | 0.2 | 112 | 534 |
| Large pelagic | 3e-03 | 0.4 | 156 | 0.02 | 17 | 993 | 9e-04 | 1 | 1003 | 0.03 | 4 | 152 | 0.05 | 22 | 475 |
| Jellyfish | 1e-02 | 1 | 119 | 0.04 | 34 | 832 | 0.04 | 41 | 1017 | 0.02 | 1 | 71 | 0.1 | 78 | 699 |
| **Total** | **1.1** | **72** | **65** | **1.7** | **641** | **376** | **0.4** | **282** | **693** | **0.3** | **11** | **46** | **3.5** | **1006** | **287** |

Table S14: Total export, corresponding sequestration and sequestration time for the different pathways considered in the model. Model run with assimilation efficiencies equal to 110 % the reference values.

| Organism | Respiration pathway | | | Fecal pellets pathway | | | Carcasses pathway | | | Other losses | | | Total | | |
|---|---|---|---|---|---|---|---|---|---|---|---|---|---|---|---|
| | Injection [PgC $yr^{-1}$] | Sequestr. [PgC] | Sequestr. time [yr] | Injection [PgC $yr^{-1}$] | Sequestr. [PgC] | Sequestr. time [yr] | Injection [PgC $yr^{-1}$] | Sequestr. [PgC] | Sequestr. time [yr] | Injection [PgC $yr^{-1}$] | Sequestr. [PgC] | Sequestr. time [yr] | Injection [PgC $yr^{-1}$] | Sequestr. [PgC] | Sequestr. time [yr] |
| Meso zoopl. | 0.1 | 0.8 | 6 | 0.2 | 30 | 141 | 0.2 | 67 | 415 | -6e-03 | -0.03 | 5 | 0.5 | 98 | 195 |
| Macro zoopl. | 0.5 | 21 | 41 | 0.2 | 52 | 294 | 0.1 | 91 | 789 | 0.3 | 9.4 | 34 | 1.1 | 173 | 162 |
| Meso-pelagic | 0.5 | 56 | 105 | 0.2 | 98 | 595 | 4e-02 | 37 | 873 | 0.2 | 16 | 65 | 1.0 | 207 | 209 |
| Forage fish | 4e-03 | 0.04 | 10 | 0.04 | 30 | 728 | 4e-03 | 4 | 968 | 0.1 | 0.4 | 7 | 0.1 | 34 | 312 |
| Large pelagic | 3e-03 | 0.4 | 156 | 0.01 | 8 | 985 | 9e-04 | 1 | 1003 | 0.03 | 5 | 154 | 0.04 | 14 | 339 |
| Jellyfish | 1e-02 | 1 | 108 | 0.02 | 15 | 831 | 0.04 | 42 | 1018 | 0.04 | 2 | 60 | 0.1 | 60 | 558 |
| **Total** | **1.2** | **79** | **66** | **0.6** | **233** | **373** | **0.4** | **242** | **661** | **0.6** | **33** | **51** | **2.8** | **587** | **208** |

Table S15: Total export, corresponding sequestration and sequestration time for the different pathways considered in the model. Model run with detritus assimilation efficiency equal to 80 % the reference value.

| Organism | Respiration pathway | | | Fecal pellets pathway | | | Carcasses pathway | | | Other losses | | | Total | | |
|---|---|---|---|---|---|---|---|---|---|---|---|---|---|---|---|
| | Injection [PgC $yr^{-1}$] | Sequestr. [PgC] | Sequestr. time [yr] | Injection [PgC $yr^{-1}$] | Sequestr. [PgC] | Sequestr. time [yr] | Injection [PgC $yr^{-1}$] | Sequestr. [PgC] | Sequestr. time [yr] | Injection [PgC $yr^{-1}$] | Sequestr. [PgC] | Sequestr. time [yr] | Injection [PgC $yr^{-1}$] | Sequestr. [PgC] | Sequestr. time [yr] |
| Meso zoopl. | 0.1 | 0.7 | 6 | 0.4 | 53 | 143 | 0.2 | 67 | 414 | -3e-02 | -0.14 | 5 | 0.6 | 120 | 192 |
| Macro zoopl. | 0.5 | 20 | 39 | 0.5 | 139 | 304 | 0.1 | 94 | 791 | 0.1 | 4.0 | 34 | 1.2 | 256 | 214 |
| Meso-pelagic | 0.5 | 54 | 103 | 0.3 | 167 | 597 | 7e-02 | 60 | 861 | 0.2 | 12 | 62 | 1.1 | 292 | 275 |
| Forage fish | 4e-03 | 0.04 | 11 | 0.1 | 70 | 726 | 4e-03 | 4 | 968 | 0.1 | 0.5 | 9 | 0.2 | 75 | 457 |
| Large pelagic | 3e-03 | 0.4 | 156 | 0.01 | 12 | 988 | 9e-04 | 1 | 1003 | 0.03 | 4 | 155 | 0.04 | 18 | 408 |
| Jellyfish | 1e-02 | 1 | 109 | 0.03 | 24 | 832 | 0.04 | 41 | 1018 | 0.03 | 2 | 62 | 0.1 | 69 | 629 |
| **Total** | **1.2** | **76** | **65** | **1.2** | **465** | **375** | **0.4** | **267** | **676** | **0.5** | **23** | **49** | **3.3** | **831** | **259** |

Table S16: Total export, corresponding sequestration and sequestration time for the different pathways considered in the model. Model run with detritus assimilation efficiency equal to 120 % the reference value.

| Organism | Respiration pathway | | | Fecal pellets pathway | | | Carcasses pathway | | | Other losses | | | Total | | |
|---|---|---|---|---|---|---|---|---|---|---|---|---|---|---|---|
| | Injection [PgC $yr^{-1}$] | Sequestr. [PgC] | Sequestr. time [yr] | Injection [PgC $yr^{-1}$] | Sequestr. [PgC] | Sequestr. time [yr] | Injection [PgC $yr^{-1}$] | Sequestr. [PgC] | Sequestr. time [yr] | Injection [PgC $yr^{-1}$] | Sequestr. [PgC] | Sequestr. time [yr] | Injection [PgC $yr^{-1}$] | Sequestr. [PgC] | Sequestr. time [yr] |
| Meso zoopl. | 0.1 | 0.4 | 5 | 0.4 | 50 | 138 | 0.2 | 65 | 413 | 1e-02 | 0.04 | 4 | 0.6 | 116 | 185 |
| Macro zoopl. | 0.5 | 19 | 38 | 0.3 | 94 | 288 | 0.1 | 86 | 782 | 0.1 | 1.9 | 32 | 1.0 | 200 | 199 |
| Meso-pelagic | 0.5 | 53 | 102 | 0.3 | 159 | 599 | 7e-02 | 58 | 820 | 0.2 | 11 | 64 | 1.0 | 280 | 275 |
| Forage fish | 3e-03 | 0.04 | 12 | 0.1 | 70 | 720 | 4e-03 | 4 | 968 | 0.1 | 0.6 | 9 | 0.2 | 74 | 445 |
| Large pelagic | 3e-03 | 0.4 | 156 | 0.01 | 12 | 983 | 9e-04 | 1 | 1003 | 0.03 | 4 | 153 | 0.04 | 18 | 405 |
| Jellyfish | 1e-02 | 1 | 119 | 0.03 | 25 | 830 | 0.04 | 41 | 1018 | 0.03 | 2 | 73 | 0.1 | 69 | 652 |
| **Total** | **1.1** | **74** | **65** | **1.1** | **408** | **375** | **0.4** | **255** | **664** | **0.4** | **19** | **55** | **3.0** | **757** | **255** |

Table S17: Total export, corresponding sequestration and sequestration time for the different pathways considered in the model. Model run with animal swimming speeds equal to 50 % the reference value.

| Organism | Respiration pathway | | | Fecal pellets pathway | | | Carcasses pathway | | | Other losses | | | Total | | |
|---|---|---|---|---|---|---|---|---|---|---|---|---|---|---|---|
| | Injection [PgC $yr^{-1}$] | Sequestr. [PgC] | Sequestr. time [yr] | Injection [PgC $yr^{-1}$] | Sequestr. [PgC] | Sequestr. time [yr] | Injection [PgC $yr^{-1}$] | Sequestr. [PgC] | Sequestr. time [yr] | Injection [PgC $yr^{-1}$] | Sequestr. [PgC] | Sequestr. time [yr] | Injection [PgC $yr^{-1}$] | Sequestr. [PgC] | Sequestr. time [yr] |
| Meso zoopl. | 0.04 | 0.1 | 2 | 0.4 | 58 | 155 | 0.1 | 61 | 412 | -3e-03 | -1e-04 | 1 | 0.6 | 119 | 215 |
| Macro zoopl. | 0.4 | 13 | 34 | 0.4 | 116 | 298 | 0.1 | 80 | 791 | 0.2 | 5.8 | 31 | 1.0 | 214 | 205 |
| Meso-pelagic | 0.3 | 33 | 130 | 0.3 | 154 | 600 | 2e-02 | 21 | 844 | 0.3 | 26 | 95 | 0.8 | 233 | 290 |
| Forage fish | 6e-03 | 0.09 | 16 | 0.1 | 79 | 725 | 4e-03 | 4 | 968 | 0.1 | 0.8 | 9 | 0.2 | 84 | 410 |
| Large pelagic | 3e-03 | 0.4 | 156 | 0.01 | 6 | 983 | 9e-04 | 1 | 1003 | 0.01 | 2 | 158 | 0.02 | 10 | 428 |
| Jellyfish | 4e-03 | 1 | 142 | 0.02 | 14 | 836 | 0.04 | 40 | 1020 | 3e-03 | 0.3 | 105 | 0.1 | 55 | 873 |
| **Total** | **0.7** | **47** | **70** | **1.1** | **427** | **371** | **0.3** | **207** | **649** | **0.6** | **34** | **62** | **2.7** | **715** | **265** |

Table S18: Total export, corresponding sequestration and sequestration time for the different pathways considered in the model. Model run with animal swimming speeds equal to 150 % the reference value.

| Organism | Respiration pathway | | | Fecal pellets pathway | | | Carcasses pathway | | | Other losses | | | Total | | |
|---|---|---|---|---|---|---|---|---|---|---|---|---|---|---|---|
| | Injection [PgC $yr^{-1}$] | Sequestr. [PgC] | Sequestr. time [yr] | Injection [PgC $yr^{-1}$] | Sequestr. [PgC] | Sequestr. time [yr] | Injection [PgC $yr^{-1}$] | Sequestr. [PgC] | Sequestr. time [yr] | Injection [PgC $yr^{-1}$] | Sequestr. [PgC] | Sequestr. time [yr] | Injection [PgC $yr^{-1}$] | Sequestr. [PgC] | Sequestr. time [yr] |
| Meso zoopl. | 0.1 | 1.3 | 9 | 0.4 | 54 | 144 | 0.2 | 67 | 419 | 4e-02 | 0.32 | 7 | 0.7 | 123 | 169 |
| Macro zoopl. | 0.6 | 23 | 37 | 0.4 | 120 | 299 | 0.1 | 92 | 773 | -0.1 | -2.0 | 31 | 1.1 | 233 | 215 |
| Meso-pelagic | 0.8 | 76 | 91 | 0.3 | 186 | 608 | 5e-01 | 456 | 980 | -0.2 | -9 | 50 | 1.4 | 708 | 500 |
| Forage fish | 3e-03 | 0.08 | 28 | 0.1 | 66 | 735 | 4e-03 | 4 | 971 | 0.1 | 0.9 | 17 | 0.2 | 71 | 472 |
| Large pelagic | 3e-03 | 0.4 | 156 | 0.03 | 30 | 1036 | 9e-04 | 1 | 1003 | 0.07 | 10 | 145 | 0.10 | 41 | 409 |
| Jellyfish | 2e-02 | 2 | 114 | 0.04 | 31 | 832 | 0.04 | 41 | 1018 | 0.04 | 3 | 63 | 0.1 | 77 | 558 |
| **Total** | **1.6** | **103** | **63** | **1.2** | **487** | **393** | **0.8** | **661** | **837** | **0.2** | **14** | **66** | **3.8** | **1265** | **333** |

Table S19: Total export, corresponding sequestration and sequestration time for the different pathways considered in the model. Model run with mesopelagic fish swimming speed equal to 50 % the reference value.

| Organism | Respiration pathway | | | Fecal pellets pathway | | | Carcasses pathway | | | Other losses | | | Total | | |
|---|---|---|---|---|---|---|---|---|---|---|---|---|---|---|---|
| | Injection [PgC $yr^{-1}$] | Sequestr. [PgC] | Sequestr. time [yr] | Injection [PgC $yr^{-1}$] | Sequestr. [PgC] | Sequestr. time [yr] | Injection [PgC $yr^{-1}$] | Sequestr. [PgC] | Sequestr. time [yr] | Injection [PgC $yr^{-1}$] | Sequestr. [PgC] | Sequestr. time [yr] | Injection [PgC $yr^{-1}$] | Sequestr. [PgC] | Sequestr. time [yr] |
| Meso zoopl. | 0.1 | 0.5 | 5 | 0.4 | 53 | 143 | 0.2 | 66 | 414 | 3e-02 | 0.13 | 4 | 0.7 | 119 | 180 |
| Macro zoopl. | 0.5 | 19 | 37 | 0.4 | 117 | 299 | 0.1 | 92 | 774 | 0.1 | 2.8 | 32 | 1.1 | 230 | 208 |
| Meso-pelagic | 0.5 | 51 | 101 | 0.2 | 145 | 600 | 3e-01 | 252 | 931 | 0.1 | 4 | 60 | 1.1 | 451 | 419 |
| Forage fish | 3e-03 | 0.05 | 14 | 0.1 | 69 | 724 | 4e-03 | 4 | 969 | 0.1 | 0.5 | 9 | 0.2 | 74 | 460 |
| Large pelagic | 3e-03 | 0.4 | 156 | 0.01 | 14 | 998 | 9e-04 | 1 | 1003 | 0.03 | 5 | 151 | 0.05 | 20 | 408 |
| Jellyfish | 1e-02 | 1 | 115 | 0.03 | 24 | 829 | 0.04 | 41 | 1018 | 0.03 | 2 | 68 | 0.1 | 69 | 643 |
| **Total** | **1.1** | **72** | **64** | **1.1** | **422** | **369** | **0.6** | **455** | **767** | **0.3** | **14** | **47** | **3.2** | **963** | **304** |

Table S20: Total export, corresponding sequestration and sequestration time for the different pathways considered in the model. Model run with mesopelagic fish swimming speed equal to 150 % the reference value.

| Organism | Respiration pathway | | | Fecal pellets pathway | | | Carcasses pathway | | | Other losses | | | Total | | |
|---|---|---|---|---|---|---|---|---|---|---|---|---|---|---|---|
| | Injection [PgC $yr^{-1}$] | Sequestr. [PgC] | Sequestr. time [yr] | Injection [PgC $yr^{-1}$] | Sequestr. [PgC] | Sequestr. time [yr] | Injection [PgC $yr^{-1}$] | Sequestr. [PgC] | Sequestr. time [yr] | Injection [PgC $yr^{-1}$] | Sequestr. [PgC] | Sequestr. time [yr] | Injection [PgC $yr^{-1}$] | Sequestr. [PgC] | Sequestr. time [yr] |
| Meso zoopl. | 0.1 | 0.6 | 5 | 0.4 | 51 | 140 | 0.2 | 66 | 414 | -3e-02 | -0.10 | 4 | 0.6 | 118 | 193 |
| Macro zoopl. | 0.5 | 20 | 39 | 0.4 | 117 | 296 | 0.1 | 83 | 792 | 0.1 | 3.7 | 34 | 1.1 | 223 | 201 |
| Meso-pelagic | 0.5 | 55 | 104 | 0.3 | 173 | 597 | 3e-02 | 25 | 851 | 0.2 | 14 | 64 | 1.1 | 268 | 250 |
| Forage fish | 3e-03 | 0.03 | 9 | 0.1 | 70 | 722 | 4e-03 | 4 | 968 | 0.1 | 0.4 | 8 | 0.2 | 74 | 466 |
| Large pelagic | 3e-03 | 0.4 | 156 | 0.01 | 12 | 986 | 9e-04 | 1 | 1003 | 0.03 | 4 | 155 | 0.04 | 17 | 408 |
| Jellyfish | 1e-02 | 1 | 114 | 0.03 | 24 | 830 | 0.04 | 41 | 1018 | 0.03 | 2 | 67 | 0.1 | 69 | 640 |
| **Total** | **1.2** | **77** | **66** | **1.2** | **447** | **377** | **0.3** | **221** | **649** | **0.4** | **24** | **59** | **3.1** | **769** | **248** |

Table S21: Total export, corresponding sequestration and sequestration time for the different pathways considered in the model. Model run with reference and maximum temperatures equal to 90 % the reference value.

| Organism | Respiration pathway | | | Fecal pellets pathway | | | Carcasses pathway | | | Other losses | | | Total | | |
|---|---|---|---|---|---|---|---|---|---|---|---|---|---|---|---|
| | Injection [PgC $yr^{-1}$] | Sequestr. [PgC] | Sequestr. time [yr] | Injection [PgC $yr^{-1}$] | Sequestr. [PgC] | Sequestr. time [yr] | Injection [PgC $yr^{-1}$] | Sequestr. [PgC] | Sequestr. time [yr] | Injection [PgC $yr^{-1}$] | Sequestr. [PgC] | Sequestr. time [yr] | Injection [PgC $yr^{-1}$] | Sequestr. [PgC] | Sequestr. time [yr] |
| Meso zoopl. | 0.1 | 0.4 | 5 | 0.4 | 49 | 134 | 0.2 | 65 | 413 | 4e-04 | 5e-04 | 4 | 0.6 | 114 | 184 |
| Macro zoopl. | 0.5 | 20 | 37 | 0.4 | 116 | 289 | 0.1 | 100 | 781 | 0.1 | 3.0 | 33 | 1.2 | 238 | 206 |
| Meso-pelagic | 0.5 | 53 | 101 | 0.3 | 155 | 596 | 1e-01 | 107 | 858 | 0.1 | 8 | 61 | 1.0 | 323 | 311 |
| Forage fish | 4e-03 | 0.04 | 12 | 0.1 | 69 | 721 | 4e-03 | 4 | 968 | 0.1 | 0.6 | 9 | 0.2 | 74 | 447 |
| Large pelagic | 3e-03 | 0.5 | 157 | 0.01 | 13 | 991 | 9e-04 | 1 | 1003 | 0.03 | 4 | 152 | 0.05 | 19 | 406 |
| Jellyfish | 1e-02 | 1 | 107 | 0.02 | 19 | 829 | 0.04 | 41 | 1016 | 0.02 | 1 | 51 | 0.1 | 61 | 685 |
| **Total** | **1.2** | **75** | **64** | **1.2** | **420** | **362** | **0.5** | **317** | **698** | **0.3** | **17** | **51** | **3.1** | **829** | **266** |

Table S22: Total export, corresponding sequestration and sequestration time for the different pathways considered in the model. Model run with reference and maximum temperatures equal to 110 % the reference value.

| Organism | Respiration pathway | | | Fecal pellets pathway | | | Carcasses pathway | | | Other losses | | | Total | | |
|---|---|---|---|---|---|---|---|---|---|---|---|---|---|---|---|
| | Injection [PgC $yr^{-1}$] | Sequestr. [PgC] | Sequestr. time [yr] | Injection [PgC $yr^{-1}$] | Sequestr. [PgC] | Sequestr. time [yr] | Injection [PgC $yr^{-1}$] | Sequestr. [PgC] | Sequestr. time [yr] | Injection [PgC $yr^{-1}$] | Sequestr. [PgC] | Sequestr. time [yr] | Injection [PgC $yr^{-1}$] | Sequestr. [PgC] | Sequestr. time [yr] |
| Meso zoopl. | 0.1 | 0.5 | 5 | 0.4 | 53 | 147 | 0.2 | 66 | 414 | 3e-02 | 0.11 | 4 | 0.6 | 120 | 187 |
| Macro zoopl. | 0.5 | 19 | 38 | 0.4 | 119 | 305 | 0.1 | 87 | 781 | 0.1 | 2.6 | 32 | 1.1 | 227 | 211 |
| Meso-pelagic | 0.5 | 52 | 103 | 0.3 | 159 | 605 | 5e-02 | 40 | 880 | 0.2 | 11 | 65 | 1.0 | 263 | 266 |
| Forage fish | 3e-03 | 0.04 | 12 | 0.1 | 69 | 727 | 4e-03 | 4 | 969 | 0.1 | 0.5 | 8 | 0.2 | 74 | 445 |
| Large pelagic | 2e-03 | 0.3 | 155 | 0.01 | 11 | 984 | 9e-04 | 1 | 1003 | 0.03 | 4 | 153 | 0.04 | 17 | 407 |
| Jellyfish | 1e-02 | 1 | 123 | 0.03 | 26 | 831 | 0.04 | 41 | 1018 | 0.03 | 2 | 84 | 0.1 | 70 | 652 |
| **Total** | **1.1** | **73** | **66** | **1.2** | **438** | **380** | **0.4** | **239** | **661** | **0.4** | **21** | **52** | **3.0** | **771** | **255** |

#### 6.2 Monte-Carlo analysis of water columns

To have a better estimate of the model's robustness in terms of behaviour and fluxes below the euphotic zone, a Monte-Carlo sensitivity analysis was performed on 5 water columns located in 5 different ocean regions (North Atlantic, North Pacific, Tropics, Subtropical gyres and Southern Ocean – cf. figure S10). For each of these water columns, we performed 500 different simulations, with all input parameters randomly selected from a normal distribution with means equal to the reference values at that location and standard deviations ranging from 10 to 50% of the reference value. Contrarily to the previous section, multiple parameters varied simultaneously.

The results proved to be very robust in terms of behaviour, with limited variations around the reference values. Passive and active injections are fairly robust to small changes in parameters. Respiration because of other losses, followed by passive injection are more sensitive to small changes in parameters than basal respiration rates and production of fecal pellets and carcasses below the euphotic zone (figures S16, S17, S18, S19, S20).

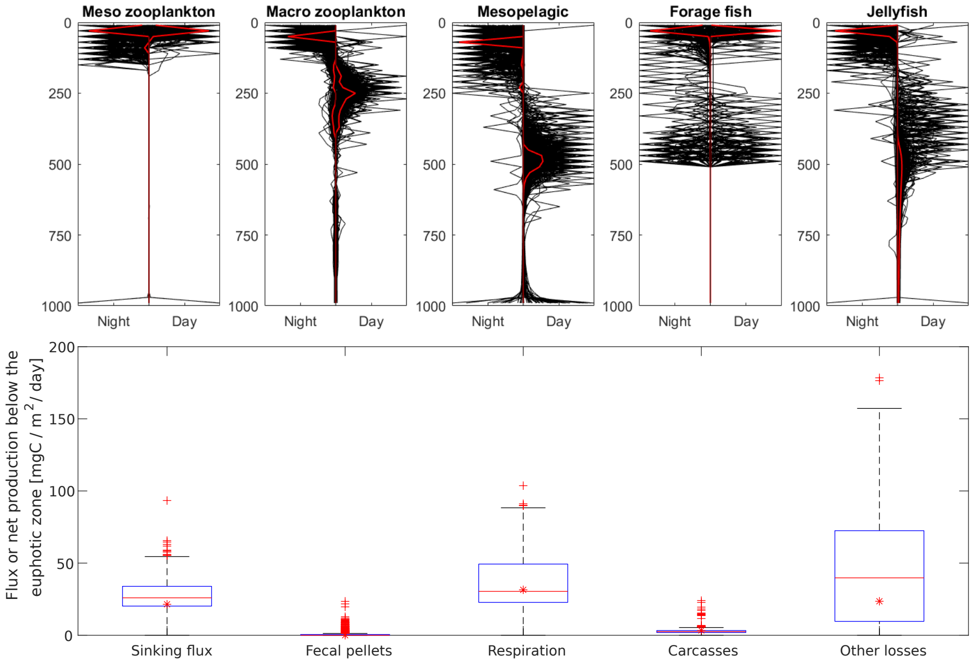

Figure S16: Monte-Carlo analysis of the model in the North Atlantic. Top panel: Distribution of the different functional groups in the North Atlantic (37°N, 20°W). Red curves are the reference vertical distributions. Bottom panel: Box plots of passive sinking flux and active injection below the euphotic zone (fecal pellet, respiration, carcasses and other losses). The red stars represent the reference exports, the red lines the medians of the Monte-Carlo simulations.

#### 6.3 Impact of deep chlorophyll maxima on behavior

Our model assumes that the distribution of phytoplankton resources is maximum at the surface. However, some locations such as subtropical gyres have deep chlorophyll maxima. To test for the sensitivity of the model to this simplification, we compare the simulated DVM patterns in a subtropical gyre (figure S21). The deep chlorophyll maximum impacts the day and night distributions of surface residents, most notably forage fish (and a little meso-zooplankton), and the night distribution of some migrants (jellyfish and macro-zooplankton). Organisms at the surface tend to gather around the chlorophyll maximum, but this has a limited impact on the residence depths of organisms in the mesopelagic (macro-zooplankton, mesopelagic fish and jellyfish during daytime). Thus, we conclude that our simplification has no major impact of the resulting sequestration estimates.

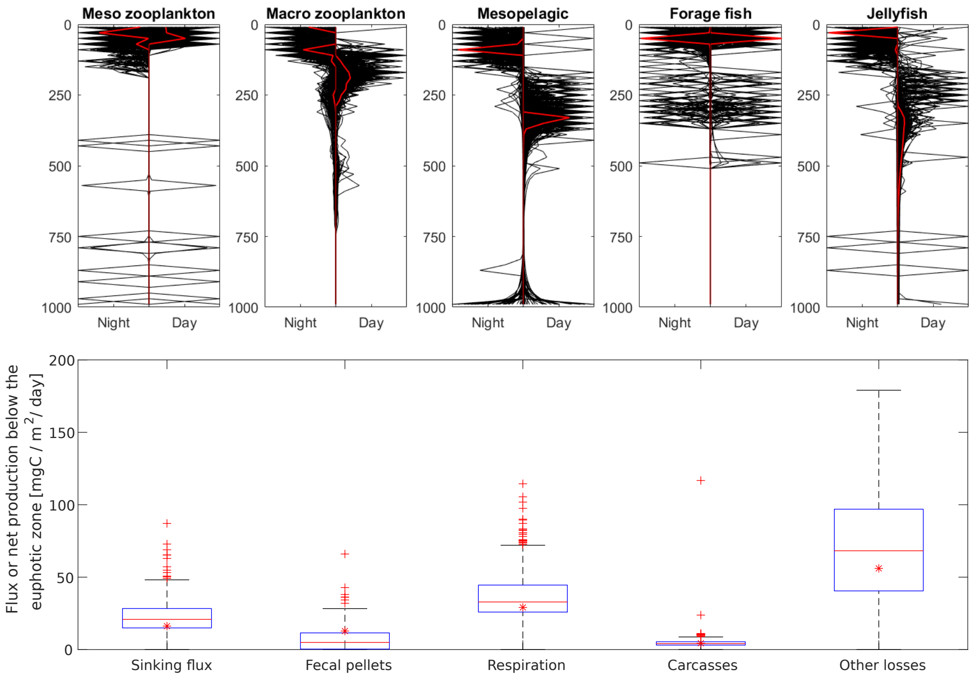

Figure S17: Monte-Carlo analysis of the model in the North Pacific. Top panel: Distribution of the different functional groups in the North Pacific (43°N, 170°W). Red curves are the reference vertical distributions. Bottom panel: Box plots of passive sinking flux and active injection below the euphotic zone (fecal pellet, respiration, carcasses and other losses). The red stars represent the reference exports, the red lines the medians of the Monte-Carlo simulations.

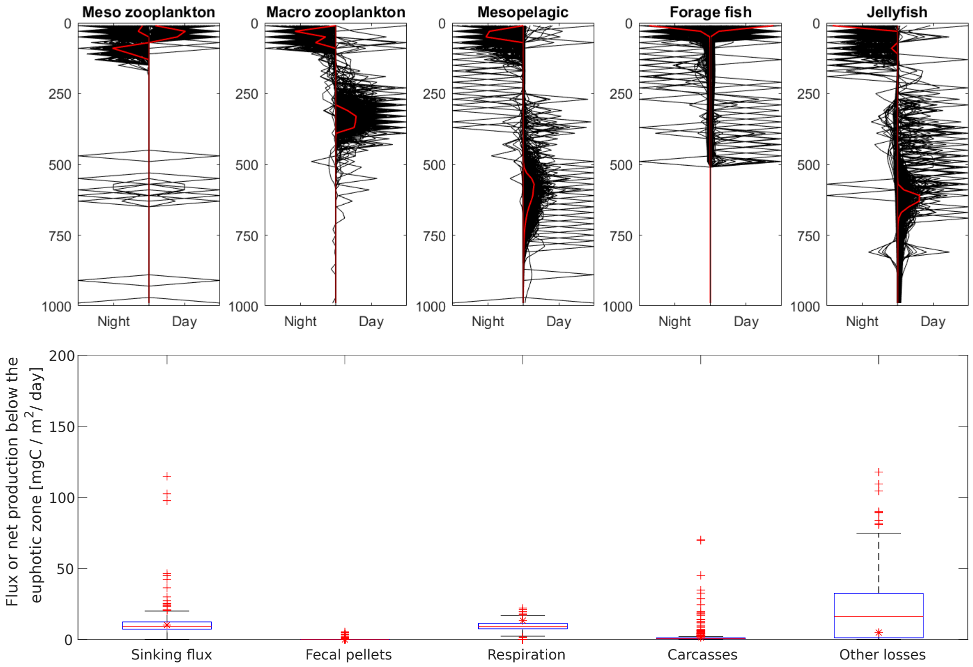

Figure S18: Monte-Carlo analysis of the model in a Subtropical gyre. Top panel: Distribution of the different functional groups in a subtropical gyre (25°N, 152°W). Red curves are the reference vertical distributions. Bottom panel: Box plots of passive sinking flux and active injection below the euphotic zone (fecal pellet, respiration, carcasses and other losses). The red stars represent the reference exports, the red lines the medians of the Monte-Carlo simulations.

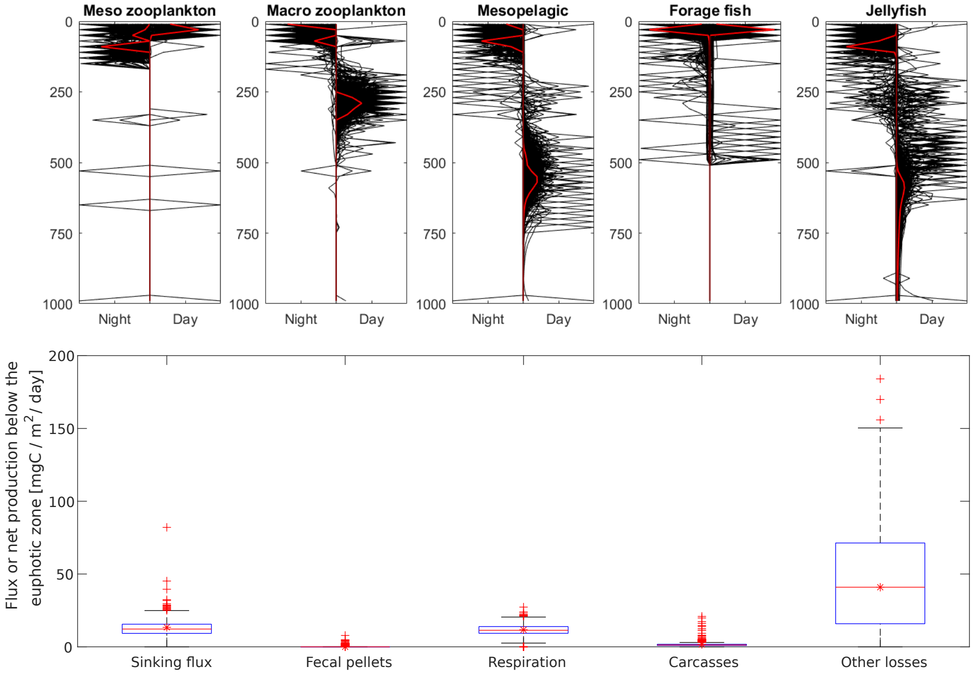

Figure S19: Monte-Carlo analysis of the model in the tropics. Top panel: Distribution of the different functional groups in the tropics (1°N, 90°E). Red curves are the reference vertical distributions. Bottom panel: Box plots of passive sinking flux and active injection below the euphotic zone (fecal pellet, respiration, carcasses and other losses). The red stars represent the reference exports, the red lines the medians of the Monte-Carlo simulations.

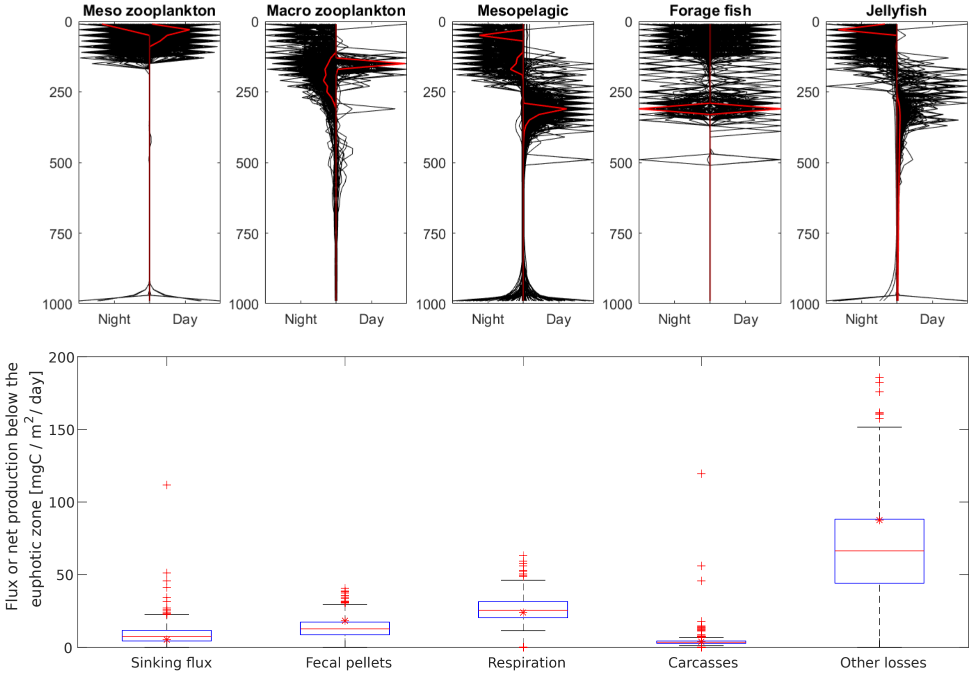

Figure S20: Monte-Carlo analysis of the model in the Southern Ocean. Top panel: Distribution of the different functional groups in the Southern Ocean (43°S, 69°E). Red curves are the reference vertical distributions. Bottom panel: Box plots of passive sinking flux and active injection below the euphotic zone (fecal pellet, respiration, carcasses and other losses). The red stars represent the reference exports, the red lines the medians of the Monte-Carlo simulations.

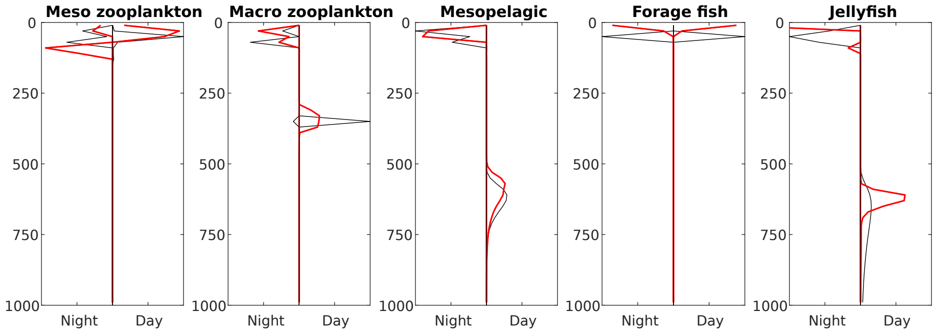

Figure S21: Simulated DVM patterns in a subtropical gyre (25N, 152W). In red is the reference run (no deep chlorophyll maximum), and in black is the run with a deep chlorophyll maximum around 50m depth.

### 7 Glossary of parameters

Table S23: Glossary of non-population-specific parameters and values

| Parameter | Signification | Value | Unit |
|---|---|---|---|
| $\sigma$ | Fraction of daylight hours in a day | 0.5 | - |
| $z$ | Depth | - | m |
| $i, j$ | Daytime (Nighttime) depth or corresponding water layer | - | m/- |
| $day$ | Boolean for dayttime (1) or nighttime (0) | - | - |
| $n$ | Number of water layers | 50 | - |
| $\Delta Z$ | Width of each water layer | 20 | m |
| $Z_{MAX}$ | Maximum depth of the 1D model | $n \cdot \Delta Z = 1000$ | m |
| $T$ | Temperature | - | °C |
| $O_2$ | Oxygen concentration | - | $mgO_2$ $L^{-1}$ |
| $p_{O_2}$ | Oxygen partial pressure | - | kPa |
| $light$ | Light level | eq. 22 | W $m^{-2}$ |
| $L_{max}$ | Maximum irradiance at the surface | eq. 45 | W $m^{-2}$ |
| $\rho_l$ | Attenuation coefficient between day and night | 1 or $10^{-5}$ | - |
| $\kappa$ | Light attenuation coefficient of the water | see figure S4 | $m^{-1}$ |
| $\rho$ | Density of seawater | 1028 | kg $m^{-3}$ |
| $\nu_w$ | Kinematic viscosity of seawater | $1.3 \cdot 10^{-6}$ | $m^2$ $s^{-1}$ |
| $G_{SG}$ | Solar constant | 1367 | W $m^{-2}$ |
| $d$ | Day of the year | - | - |
| $\beta$ | Solar altitude angle | eq. 46 | rad |
| $h$ | Hour angle | 0 | rad |
| $\delta$ | Solar inclination angle | $23.45 \sin(360 \frac{d+284}{365})$ | rad |
| $z_0$ | Mixed layer depth | see figure S4 | m |
| $R$ | Vertical profile of phytoplankton | eq. 47 | gC $m^{-3}$ |
| $\psi$ | Maximum fraction of detritus that can be consumed daily | 0.8 | $day^{-1}$ |
| $\tilde{F}_X^{D_Y}$ | Corrected $F$ with the maximum ingestion of detritus. Only valid for $X = C$ and $X = P$ | eq. 39 | |
| $\alpha$ | Bacterial degradation rate of fecal matter | eq. 42 | $day^{-1}$ |
| $\alpha_0$ | Maximum bacterial degradation rate | 0.25 | $day^{-1}$ |
| $Q_{bac}$ | $Q_{10}$ factor for bacterial degradation | 2 | - |
| $T_{ref,bac}$ | Reference temperature for bacterial degradation rate | 10 | ° C |
| $K_{O_2}$ | Half-saturation constant for bacterial degradation | 20 | $mgO_2$ $L^{-1}$ |
| $\lambda_t$ | Rescaling factor in the Replicator equation | $0.05/\max(W_X(i,j))$ | - |
| $\delta_t$ | Time step of the Replicator equation | - | - |
| $\mathbf{A}$ | Advection-diffusion matrix transport operator from OCIM | - | - |

Table S24: Glossary of population-specific functions and rates

| Symbol | Signification | Expression | Unit |
|---|---|---|---|
| $X$ | Placeholder for the different populations: $C$ (mesozooplankton), $P$ (macrozooplankton), $M$ (mesopelagic), $F$ (forage), $A$ (large pelagic), $J$ (jellyfish), or $D_X$ (fecal pellets produced by $X$) | - | - |
| $X_{ij}$ | Fraction of population $X$ following strategy $ij$ | eq. 44 | - |
| $D_X$ | Concentration of fecal pellets created by $X$ in the water column | eq. 41 | gC $m^{-3}$ |
| $X(i, day)$ | Concentration of organisms $X$ at $i$ during $day$ | eq. 2 | gC $m^{-3}$ |
| $W_X(i,j)$ | Fitness of an individual $X$ with strategy $ij$ | eq. 9 | - |
| $g_X(i,j)$ | Growth rate of an individual $X$ with strategy $ij$ | $\nu_X - Q_X - C_{migr,X}$ | $day^{-1}$ |
| $m_X(i,j)$ | Mortality rate of an individual $X$ with strategy $ij$ | $\mu_X + \mu_{0X}$ | $day^{-1}$ |
| $Q_X(i,j)$ | Strategy-dependent standard metabolic rate | eq. 10 | $day^{-1}$ |
| $C_{migr,X}(i,j)$ | Migration cost | eq. 35 | $day^{-1}$ |
| $\tilde{Q}_X(z)$ | Depth-dependent standard metabolic cost | eq. 11 | $day^{-1}$ |
| $\nu_X(i,j)$ | Strategy-dependent assimilation rate | eq. 13 | $day^{-1}$ |
| $\tilde{\nu}_{X,i,j}(z)$ | Depth-dependent assimilation rate | eq. 17 | $day^{-1}$ |
| $u_{X,i,j}(z)$ | (Cruising) swimming speed | eq. 14 | m $day^{-1}$ |
| $u_{max,X}$ | Maximum swimming speed | eq. 26 | m $s^{-1}$ |
| $I_{max,X,i,j}(z)$ | Maximum ingestion rate | eq. 14 | gC $day^{-1}$ |
| $SMR_{0,X}$ | Standard metabolic cost at $T_{ref}$ | eq. 12 | $day^{-1}$ |
| $u_{0,X}$ | Reference swimming speed | eq. 15 | m $day^{-1}$ |
| $s$ | Function used for $SMR$ and $MMR$ | eq. 3 | $day^{-1}$ |
| $SMR$ | Standard Metabolic Rate | eq. 4 and 6 | $day^{-1}$ |
| $MMR$ | Maximum Metabolic Rate | eq. 5 and 7 | $day^{-1}$ |
| $AS$ | Aerobic scope | $MMR - SMR$ | $day^{-1}$ |
| $\tilde{S}_X(i,j,day)$ | $AS(z=i)$ if $day=1$, $AS(z=j)$ if $day=0$ | - | $day^{-1}$ |
| $S_X(i,j,z)$ | Available aerobic scope | eq. 8 | $day^{-1}$ |
| $S_{0,X}$ | Reference metabolic scope for $u_0$ & $I_{max0}$ | $\max S_X$ | $day^{-1}$ |
| $F_X^Y(z)$ | Depth-dependent specific ingestion rate of prey Y by pred. X | eq. 18, 20 | $day^{-1}$ |
| $E_X^Y(z)$ | Depth-dependent encounter rate of $Y$ by $X$ | eq. 19 | gC $day^{-1}$ |
| $\Phi(X,Y)$ | Preference function of $X$ for $Y$ | table S26 | - |
| $\Gamma_X^Y(z, day)$ | Capture probability of $Y$ by $X$ during an attack event | eq. 29 | - |
| $r_{esc}$ | Length of escape jump by prey | eq. 27 | m |
| $r_{detec}$ | Prey detection distance of predator | $\Lambda_Y^X(z, day)$ | m |
| $r_{capt}$ | Capture distance for predator | $0.1 l_X$ | m |
| $r_{attac}$ | Length of attack jump for predator | eq. 28 | m |
| $v_{esc}$ | Volume of the escape sphere | $\frac{4}{3}\pi r_{esc}^3$ | $m^3$ |
| $v_{capt}$ | Volume of the escape sphere swept by predator | - | $m^3$ |
| $V_X^Y(z)$ | Clearance rate of $X$ for $Y$ | eq. 23, 24, 25 | $m^3$ $day^{-1}$ |
| $\Lambda_X^Y(z, day)$ | Visual range of $X$ | eq. 21 | m |
| $\mu_{X,i,j}$ | Mortality rate due to predation for strategy $ij$ | eq. 30 | $day^{-1}$ |
| $\tilde{\mu}_X(z, day)$ | Mortality rate due to predation at (z,day) | eq. 31 | $day^{-1}$ |
| $Dr_X$ | Hydrodynamic drag | eq. 32 | kg m $s^{-2}$ |
| $C_D$ | Drag coefficient | eq. 33 | - |
| $Re_X$ | Reynolds number | eq. 34 | - |
| $W_X^0$ | Fitness of population $X$ at the Nash equilibrium | eq. 43 | - |
| $X'_{ij}$ | Intermediate value in the Replicator equation | eq. 44 | - |
| $D_{crea,X}$ | Creation rate of fecal pellets by $X$ | eq. 36, 37 | gC $m^{-3}$ $day^{-1}$ |
| $D_{conso,X}$ | Consumption rate of fecal pellets $X$ | eq. 38 | gC $m^{-3}$ $day^{-1}$ |
| $\zeta_X$ | Source term of detritus $X$ in each water layer | eq. 40 | gC $m^{-3}$ $day^{-1}$ |
| $J_{res}$ | Source of DIC | - | gC $m^{-3}$ $day^{-1}$ |
| $C_{in}$ | DIC due to respiration | eq. 49 | gC $m^{-3}$ |

Table S25: Glossary of population-dependent parameters. Subscript $_X$ was omitted for readability.

| Param. | Signification | Unit | Mesozoo-plankton | Macrozoo-plankton | Mesopelagic fish | Forage fish | Large pelagic fish | Tactile predators |
|---|---|---|---|---|---|---|---|---|
| $l$ | Length of a typical individual | m | $5 \cdot 10^{-4}$ | $5 \cdot 10^{-3}$ | 0.03 | 0.2 | 1 | 0.1 |
| $w$ | Carbon weight of a typical individual | gC | $2 \cdot 10^{-5}$ | $1.5 \cdot 10^{-4}$ | 0.18 | 15.2 | 1900 | 11.8 |
| $\tilde{X}$ | Biomass in the water column | gC m$^{-2}$ | fig. S5 | fig. S5 | fig. S5 | fig. S5 | fig. S5 | $10^{-3}$ |
| $Q_{10}$ | Temperature coefficient | - | 2 | 2 | 2 | 1.5 | 2 | 3 |
| $T_{ref}$ | Reference temperature | °C | $\frac{1}{100}\int_0^{100} T(z)dz$ | $\frac{1}{950}\int_{50}^{1000} T(z)dz$ | $\frac{1}{950}\int_{50}^{1000} T(z)dz$ | $\frac{1}{100}\int_0^{100} T(z)dz$ | $\frac{1}{950}\int_{50}^{1000} T(z)dz$ | $\frac{1}{400}\int_{300}^{700} T(z)dz$ |
| $T_{max}$ | Maximum temperature | °C | $\max_{z<20} T(z)$ | $\max_{z>20} T(z)$ | $\max_{z>50} T(z)$ | $\max T(z)$ | $\max T(z)$ | $\max_{z>50} T(z)$ |
| $\varphi$ | Assimilation efficiency | - | 0.8 | 0.85 (0.7 if detritus) | 0.8 | 0.85 | 0.75 | 0.8 |
| $R_0$ | Maximum detection distance | m | $2l_C$ | $2l_P$ | table S27 | table S27 | table S27 | 0.2 |
| $K_e$ | Half-saturation constant for light | W m$^{-2}$ | - | - | $10^{-6}$ | 1 | 1 | - |
| $\gamma$ | Cross-sectional area efficiently scanned | - | - | 0.5 | 0.5 | 0.5 | 0.5 | - |
| $f$ | Filtering efficiency of typical predators | - | - | - | - | - | - | 0.089 |
| $\mu_0$ | Background mortality rate | day$^{-1}$ | 0.005 | $0.003 + 0.5\frac{light(z_i,1)+light(z_j,0)}{light(0,1)+light(0,0)}$ | $0.005 + 0.8\frac{light(z_i,1)+light(z_j,0)}{light(0,1)+light(0,0)}$ | $1.3 \cdot 10^{-4}$ | $0.7 \cdot 10^{-4}$ | 0.02 |
| $p_{crit}$ | Critical oxygen partial pressure | kPa | 3.5 | 0.1 | 0.2 | 5 | 0.5 | 1 |
| $p_{reg}$ | Transition between oxy-regulation and conformism | % oxy. satur. | 0.6 | 0.3 | - | - | - | 0.6 |
| $\Delta_{MR}$ | Maximum $AS$ compared to $SMR$ | - | 3 | 3 | 3 | 6 | 6 | 2 |
| $\omega$ | Fecal pellet sinking speed | m day$^{-1}$ | 100 | 150 | 300 | 400 | 800 | 500 |
| $\omega_{carc}$ | Carcasses sinking speed | m day$^{-1}$ | 200 | 400 | 600 | 800 | 1500 | 800 |
| $\epsilon$ | Swimming efficiency | - | 0.01 | 0.01 | 0.01 | 0.01 | 0.01 | 0.01 |
| $I_{max0}$ | Reference max. ingestion rate | gC day$^{-1}$ | $3 \cdot 10^{-6}$ | $3.6 \cdot 10^{-5}$ | 0.013 | 3 | 227 | - |

Table S26: Preference function values. Predators are in line, prey in column.

| | Phytoplankton | Detritus | MesoZPK | MacroZPK | Forage | Meso | Tact |
|---|---|---|---|---|---|---|---|
| MesoZPK | 1 | 0 | - | 0 | 0 | 0 | 0 |
| MacroZPK | 1 | 1 | 1 | - | 0 | 0 | 0 |
| Forage | 0 | 0 | 1 | 1 | - | 1 | 0 |
| Meso | 0 | 0 | 0.1 | 1 | 0 | - | 0 |
| Tact | 0 | 0 | 1 | 1 | 0 | 0 | - |
| Top | 0 | 0 | 0 | 0 | 1 | 0.2 | 0.1 |

Table S27: Maximum visual range in m. Visual "viewers" (predators or prey) are in lines, and their targets are in columns.

| | MesoZPK | MacroZPK | Meso | Forage | Tact | Large pelagic |
|---|---|---|---|---|---|---|
| Mesopelagic | 0.04 | 0.05 | - | 0.3 | 0.3 | 1 |
| Forage fish | 0.2 | 0.2 | 2 | - | - | 3 |
| Large pelagic | - | - | 2 | 2 | 4 | - |

### References

[1] José Luis Acuña, Ángel López-Urrutia, and Sean Colin. Supplementary material- Faking giants: The evolution of high prey clearance rates in jellyfishes. *Science*, 333(6049):1627–1629, 2011.

[2] Ken H. Andersen. *Fish Ecology, Evolution, and Exploitation : A New Theoretical Synthesis*. Princeton University Press, monographs edition, 2019.

[3] Daniele Bianchi, Charles Stock, Eric D. Galbraith, and Jorge L. Sarmiento. Diel vertical migration: Ecological controls and impacts on the biological pump in a one-dimensional ocean model. *Global Biogeochemical Cycles*, 27(2):478–491, 2013.

[4] Ryan P Bos, Tracey T Sutton, and Tamara M Frank. State of Satiation Partially Regulates the Dynamics of Vertical Migration. *Frontiers in Marine Science*, 8:607228, 2021.

[5] Ken O. Buesseler, Philip W. Boyd, Erin E. Black, and David A. Siegel. Metrics that matter for assessing the ocean biological carbon pump. *Proceedings of the National Academy of Sciences*, 117(18):201918114, 2020.

[6] P Caparroy, Uffe H. Thygesen, and André W. Visser. Modelling the attack success of planktonic predators: patterns and mechanisms of prey size selectivity. *Journal of Plankton Research*, 22(10):1871–1900, 2000.

[7] B. R. Carter, R. A. Feely, S. K. Lauvset, A. Olsen, T. DeVries, and R. Sonnerup. Preformed Properties for Marine Organic Matter and Carbonate Mineral Cycling Quantification. *Global Biogeochemical Cycles*, 35(1), 2021.

[8] G Claireaux, D M Webber, J-P Lagardère, and S R Kerr. Influence of water temperature and oxygenation on the aerobic metabolic scope of Atlantic cod ( Gadus morhua ). *Journal of Sea Research*, 44:257–265, 2000.

[9] Clément de Boyer Montégut, Gurvan Madec, Albert S. Fischer, Alban Lazar, and Daniele Iudicone. Mixed layer depth over the global ocean: An examination of profile data and a profile-based climatology. *Journal of Geophysical Research C: Oceans*, 109(12):1–20, 2004.

[10] Tim DeVries. The oceanic anthropogenic CO2 sink: Storage, air-sea fluxes, and transports over the industrial era. *Global Biogeochemical Cycles*, 28(7):631–647, 2014.

[11] Tim DeVries and Thomas Weber. The export and fate of organic matter in the ocean: New constraints from combining satellite and oceanographic tracer observations. *Global Biogeochemical Cycles*, 31(3):535–555, 2017.

[12] Paolo Domenici. The scaling of locomotor performance in predator–prey encounters: from fish to killer whales. *Comparative Biochemistry and Physiology Part A: Molecular & Integrative Physiology*, 131(1):169–182, dec 2001.

[13] W. Ekau, H. Auel, H. O. Portner, and D. Gilbert. Impacts of hypoxia on the structure and processes in pelagic communities (zooplankton, macro-invertebrates and fish). *Biogeosciences*, 7(5):1669–1699, 2010.

[14] Rasmus Ern, Tommy Norin, A. Kurt Gamperl, and Andrew J. Esbaugh. Oxygen dependence of upper thermal limits in fishes. *The Journal of Experimental Biology*, 219(21):3376–3383, 2016.

[15] H.E. Garcia, C.R. Weathers, C.R. Paver, I. Smolyar, T.P. Boyer, R.A. Locarnini, M.M. Zweng, A.V. Mishonov, O.K. Baranova, D. Seidov, and J.R. Reagan. WORLD OCEAN ATLAS 2018 Volume 3: Dissolved Oxygen, Apparent Oxygen Utilization, and Dissolved Oxygen Saturation. Technical report, Silver Spring, MD, 2019.

[16] James F. Gilliam and Douglas F. Fraser. Habitat Selection Under Predation Hazard: Test of a Model with Foraging Minnows. *Ecology*, 68(6):1856–1862, dec 1987.

[17] Josef Hofbauer and Karl Sigmund. Evolutionary Game Dynamics. *Bulletin (New Series) of the American mathematical society*, 40(403):479–519, 2003.

[18] Kim N. Holland, Richard W. Brill, Randolph K.C. Chang, John R. Sibert, and David A. Fournier. Physiological and behavioural thermoregulation in bigeye tuna (Thunnus obesus). *Nature*, 358(6385):410–412, 1992.

[19] Alasdair I. Houston, John M. McNamara, and John M. C. Hutchinson. General results concerning the trade-off between gaining energy and avoiding predation. *Philosophical Transactions of the Royal Society of London. Series B: Biological Sciences*, 341(1298):375–397, sep 1993.

[20] Mark E. Huntley and Meng Zhou. Influence of animal on turbulence in the sea. *Marine Ecology Progress Series*, 273:65–79, 2004.

[21] IMOS. IMOS BASOOP sub facility, 2021.

[22] Rubao Ji and PJS Franks. Vertical migration of dinoflagellates: model analysis of strategies, growth, and vertical distribution patterns. *Marine Ecology Progress Series*, 344:49–61, aug 2007.

[23] Rainer Kiko, Helena Hauss, Friedrich Buchholz, and Frank Melzner. Ammonium excretion and oxygen respiration of tropical copepods and euphausiids exposed to oxygen minimum zone conditions. *Biogeosciences*, 13(8):2241–2255, apr 2016.

[24] Thomas Kiørboe, Anders Andersen, V. J. Langlois, and H. H. Jakobsen. Unsteady motion: escape jumps in planktonic copepods, their kinematics and energetics. *Journal of The Royal Society Interface*, 7(52):1591–1602, nov 2010.

[25] Thomas Kiørboe and Andrew G. Hirst. Shifts in Mass Scaling of Respiration, Feeding, and Growth Rates across Life-Form Transitions in Marine Pelagic Organisms. *The American Naturalist*, 183(4):E118–E130, 2014.

[26] T. A. Klevjer, X. Irigoien, A. Røstad, E. Fraile-Nuez, V. M. Benítez-Barrios, and Stein Kaartvedt. Large scale patterns in vertical distribution and behaviour of mesopelagic scattering layers. *Scientific Reports*, 6(1):19873, apr 2016.

[27] R.A. Locarnini, A.V. Mishonov, O.K. Baranova, T.P. Boyer, M.M. Zweng, H.E. Garcia, J.R. Reagan, D. Seidov, K.W. Weathers, C.R. Paver, and I.V. Smolyar. World Ocean Atlas, Volume 1: Temperature, 2019.

[28] Cathy H. Lucas, Daniel O.B. Jones, Catherine J. Hollyhead, Robert H. Condon, Carlos M. Duarte, William M. Graham, Kelly L. Robinson, Kylie A. Pitt, Mark Schildhauer, and Jim Regetz. Gelatinous zooplankton biomass in the global oceans: Geographic variation and environmental drivers. *Global Ecology and Biogeography*, 23(7):701–714, 2014.

[29] Michael J. Lutz, Ken Caldeira, Robert B. Dunbar, and Michael J. Behrenfeld. Seasonal rhythms of net primary production and particulate organic carbon flux to depth describe the efficiency of biological pump in the global ocean. *Journal of Geophysical Research: Oceans*, 112(10), 2007.

[30] Frederic Melin. GMIS - MODIS-AQUA Monthly climatology sea surface diffuse attenuation coefficient at 490nm (9km) in mˆ-1. Technical report, European Commision, Joint Research Centre (JRC), 2013.

[31] Mohammad H. Naraghi and Gregory Etienne. Solar Panel Orientation and Modeling Based on Hourly Clearness Index. *ASME 2012 6th International Conference on Energy Sustainability, Parts A and B*, (July 2012):105, 2012.

[32] John Nash. Non-Cooperative Games. *The Annals of Mathematics*, 54(2):286, sep 1951.

[33] Göran E. Nilsson and Sara Östlund-Nilsson. Does size matter for hypoxia tolerance in fish? *Biological Reviews*, 83(2):173–189, 2008.

[34] Colleen M. Petrik, Charles A. Stock, Ken Haste Andersen, P. Daniël van Denderen, and James R. Watson. Bottom-up drivers of global patterns of demersal, forage, and pelagic fishes. *Progress in Oceanography*, 176(June):102124, sep 2019.

[35] Jérôme Pinti, Thomas Kiørboe, Uffe H. Thygesen, and André W. Visser. Trophic interactions drive the emergence of diel vertical migration patterns: a game-theoretic model of copepod communities. *Proceedings of the Royal Society B: Biological Sciences*, 286(1911):20191645, sep 2019.

[36] Jérôme Pinti and André W. Visser. Predator-Prey Games in Multiple Habitats Reveal Mixed Strategies in Diel Vertical Migration. *The American Naturalist*, 193(3):E65–E77, mar 2019.

[37] Polar Data Centre. British Antarctic Survey. *Raw acoustic data collected by ship-borne EK60 echosounder in the Atlantic Ocean (AMT24, AMT25, AMT26, AMT29).*, 2020.

[38] Roland Proud, Martin J. Cox, and Andrew S. Brierley. Biogeography of the Global Ocean's Mesopelagic Zone. *Current Biology*, 27(1):113–119, 2017.

[39] Roland Proud, Nils Olav Handegard, Rudy J Kloser, Martin J Cox, and Andrew S Brierley. From siphonophores to deep scattering layers: uncertainty ranges for the estimation of global mesopelagic fish biomass. *ICES Journal of Marine Science*, 76(3):718–733, may 2019.

[40] Roland Proud, Nils Olav Handegard, Rudy J Kloser, Martin J Cox, and Andrew S Brierley. From siphonophores to deep scattering layers: uncertainty ranges for the estimation of global mesopelagic fish biomass. *ICES Journal of Marine Science*, 76(3):718–733, may 2019.

[41] Nicholas J. Rogers, Mauricio A. Urbina, Erin E. Reardon, David J. McKenzie, and Rod W. Wilson. A new analysis of hypoxia tolerance in fishes using a database of critical oxygen level (Pcrit). *Conservation Physiology*, 4(1):1–19, 2016.

[42] K Salonen, J Sarvala, I Hakala, and M L Viljanen. The relation of energy and organic carbon in aquatic invertebrates. *Limnology and Oceanography*, 21 (5)(5):724–730, 1976.

[43] Peter Schuster and Karl Siegmund. Replicator Dynamics. *Journal of Theoretical Biology*, 100:533–538, 1983.

[44] B. A. Seibel. Critical oxygen levels and metabolic suppression in oceanic oxygen minimum zones. *Journal of Experimental Biology*, 214(2):326–336, 2011.

[45] Brad A. Seibel and Curtis Deutsch. Oxygen supply capacity in animals evolves to meet maximum demand at the current oxygen partial pressure regardless of size or temperature. *Journal of Experimental Biology*, 223(12), 2020.

[46] Brad A. Seibel, Jillian L. Schneider, Stein Kaartvedt, Karen F. Wishner, and Kendra L. Daly. Hypoxia Tolerance and Metabolic Suppression in Oxygen Minimum Zone Euphausiids: Implications for Ocean Deoxygenation and Biogeochemical Cycles. *Integrative and Comparative Biology*, 56(4):510–523, 2016.

[47] Ben Speers-Roesch, Milica Mandic, Derrick J.E. Groom, and Jeffrey G. Richards. Critical oxygen tensions as predictors of hypoxia tolerance and tissue metabolic responses during hypoxia exposure in fishes. *Journal of Experimental Marine Biology and Ecology*, 449:239–249, 2013.

[48] Charles A. Stock, Jasmin G. John, Ryan R. Rykaczewski, Rebecca G. Asch, William W.L. Cheung, John P. Dunne, Kevin D. Friedland, Vicky W.Y. Lam, Jorge L. Sarmiento, and Reg A. Watson. Reconciling fisheries catch and ocean productivity. *Proceedings of the National Academy of Sciences of the United States of America*, 114(8):E1441–E1449, 2017.

[49] Uffe H. Thygesen, Lene Sommer, Karen Evans, and Toby A. Patterson. Dynamic optimal foraging theory explains vertical migrations of Bigeye tuna. *Ecology*, 97(7):1852–1861, 2016.

[50] Gordon R. Ultsch and Matthew D. Regan. The utility and determination of Pcrit in fishes. *Journal of Experimental Biology*, 222(22):1–9, 2019.

[51] Hans van Someren Gréve, Rodrigo Almeda, and Thomas Kiørboe. Motile behavior and predation risk in planktonic copepods. *Limnology and Oceanography*, 62(5):1810–1824, 2017.

[52] André W. Visser. Motility of zooplankton: Fitness, foraging and predation. *Journal of Plankton Research*, 29(5):447–461, 2007.
